# A meta-interaction basis for cell-cell communication in tissues

**DOI:** 10.64898/2026.09.21.753369

**Authors:** Junjie Tang, Shaoheng Liang, Shahul Alam, Wenduo Cheng, Yang Zhang, Jian Ma

## Abstract

Tissue function depends on signals exchanged between cells and the responses they elicit. Yet whether diverse cell-cell interactions *in situ* form recurrent sender-receiver programs remains unclear. We present SpiderNet, an interpretable representation-learning framework that discovers such directed programs as a compact basis of cell-cell meta-interactions (MIs) from spatial transcriptomics. SpiderNet jointly learns which sender regulators, ligand-receptor pairs, and receiver targets define each MI and where each program is active across neighboring cell pairs. The resulting representation traces multicellular relays and links communication to cell states, perturbation responses, and phenotypes. SpiderNet recovers ground-truth MIs and their molecular components in simulations and, in real tissues, shows stronger direction-specific agreement with independently curated regulatory programs in senders and receivers than alternative methods. Across more than 5.8 million spatially profiled cells, SpiderNet resolves an SPP1-THBS relay linking monocytes, fibroblasts, and tumor cells within an immune-suppressive ovarian cancer niche, predicts T-cell responses to held-out melanoma-cell perturbations, and identifies a T-cell-associated brain-aging program and age-predictive signals that transfer across regions and platforms. It reveals a recurrent pan-cancer COLLAGEN-linked fibroblast-tumor program whose projected abundance in independent cohorts is associated with poorer survival and non-response to immunotherapy. SpiderNet thus establishes MIs as a reusable organizational layer between molecular interactions and tissue phenotypes, providing a framework to resolve, compare, trace, and perturb multicellular regulation *in situ*.

## Introduction

Cell-cell communication (CCC) in tissues couples signals exchanged between cells to intracellular responses, shaping tissue organization and function [1]. Rather than arising from isolated molecular contacts, CCC is often organized through coordinated combinations of molecular interactions and cellular states [2, 3]. During immune-cell activation, receptor signaling, adhesion, and downstream transcriptional responses are integrated at the immunological synapse [4]; in tumors, stromal signals, extracellular-matrix interactions, and malignant-cell responses likewise form coordinated multicellular programs [5]. A key challenge is therefore not only to determine where communication occurs, but also to identify the organizational unit through which heterogeneous local interactions generate coherent tissue-level behavior. This motivates the search for recurrent, directed cell-pair programs that could underlie such organization by coupling sender state, intercellular signaling, and receiver response.

Spatial transcriptomics (ST) provides an opportunity to study multicellular communication *in situ* by jointly measuring cellular gene expression and spatial context in intact tissues [6, 7]. As ST atlases expand, a critical goal is to move from cataloging cellular neighborhoods and candidate interactions toward uncovering organizing principles of multicellular regulation. Computational methods have resolved complementary aspects of spatial CCC, including screening spatial ligand-receptor (LR) associations [8–10], inferring LR-mediated pathway activity [11, 12], modeling neighbor-induced transcriptional effects [13– 15], and detecting multi-hop relay networks [16]. Yet these approaches generally represent intercellular signals, sender states, and receiver responses separately or sequentially [17], rather than as linked components of jointly learned, recurrent cell-pair programs.

Addressing this gap requires jointly learning which molecular components define each communication program and how that program recurs across directed cell pairs, linking sender state to receiver response through LR signaling. Such a representation would reveal where these programs are active, how they form multicellular relays, and how their activity varies with cell state, perturbation, and phenotype. Learning this representation from ST is challenging for three reasons. First, communication is defined over ordered cell pairs rather than individual cells, creating a large inference space for recurrent programs. Second, cellular gene expression combines cell-type-intrinsic expression with communication-associated transcriptional responses, making the latter difficult to distinguish. Third, each program spans sender regulation, LR signaling, and receiver response, making it difficult to determine which molecular components belong to the same program.

Here we present SpiderNet, an interpretable framework that learns a compact basis of cell-cell meta-interactions (MIs) from ST data. Each MI is a recurrent, directed communication module linking sender-side regulators to receiver-side targets through LR pairs, with activity quantified over ordered neighboring cell pairs across tissue. SpiderNet learns this basis by jointly reconstructing cell-pair LR co-expression and cellular gene expression, so that the same latent MI activities jointly explain intercellular LR signaling and sender- and receiver-side transcriptional variation, while a separate component captures cell-type-intrinsic expression. This directly ties each learned communication program to its molecular composition rather than assigning interpretation only post hoc. We establish recovery of ground-truth MIs in simulations and direction-specific agreement with independently curated regulatory programs in real tissues. Across cancer, genetic perturbation, and aging, the MI representation resolves communication-associated cell states and multicellular relays, predicts non-cell-autonomous responses to held-out perturbations, and identifies recurrent programs linked to tissue phenotypes and clinical out-comes. Together, these analyses establish MIs as an intermediate-scale representation for resolving, comparing, tracing, and perturbing multicellular regulation *in situ*.

## Results

### SpiderNet learns a recurrent, directional basis of cell-cell communication

SpiderNet learns a compact, directional basis of cell-cell meta-interactions (MIs) from ST data. Each MI defines a recurrent communication module with molecular signatures spanning sender-side regulators, ligand-receptor (LR) bridges, and receiver-side targets, while each directed neighboring cell pair receives an MI activity vector specifying which modules are engaged and at what strengths (**Fig**. 1). Here, regulators and targets denote genes associated with the sending and receiving sides of an MI, respectively. This factorization allows the same MI to recur across many cell pairs and multiple MIs to be engaged by a single directed pair. Reversing the sender and receiver can yield a different MI activity profile. By jointly reconstructing LR co-expression and gene expression, SpiderNet integrates intercellular signaling with cellular transcriptional state while separately modeling cell-type-intrinsic expression and MI-associated transcriptional variation.

**Figure 1:**
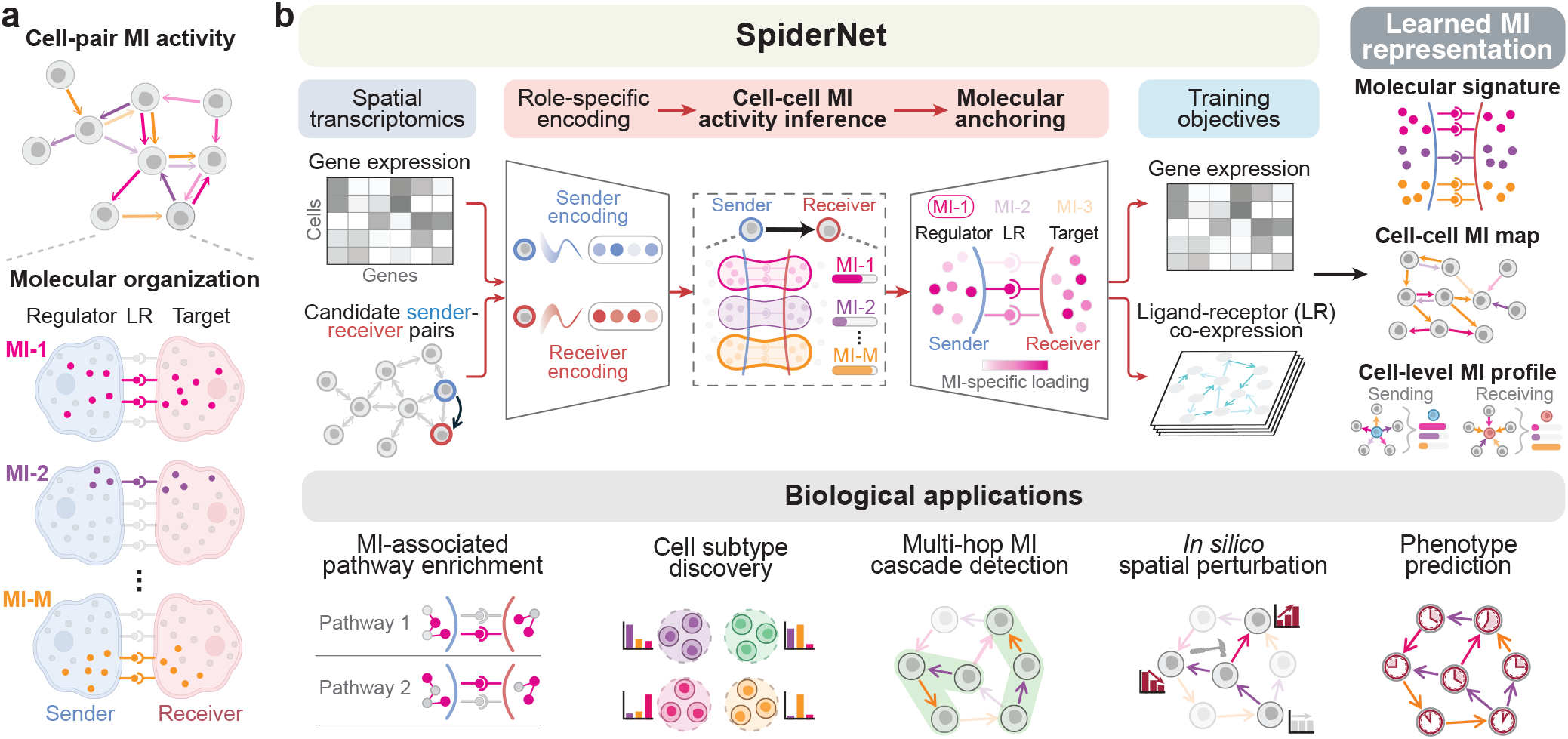
Overview of SpiderNet. **a**, Conceptual organization of recurrent, directional meta-interactions (MIs). Directed neighboring cell pairs carry activity profiles across multiple MIs, while each MI represents a coordinated molecular program linking sender-side regulators to receiver-side targets through ligand-receptor (LR) pairs. **b**, SpiderNet learns cell-cell MIs from spatial transcriptomics by integrating gene expression with spatial context. Spatial proximity defines ordered neighboring cell pairs, with each cell encoded separately in sender and receiver roles to infer which communication programs are engaged and how strongly. A dual reconstruction objective molecularly anchors the inferred MIs by reconstructing LR co-expression and cellular gene expression, yielding interpretable non-negative loadings for sender-side regulators, LR bridges, and receiver-side target genes. Outputs include MI molecular signatures, edge-level MI activities, and cell-level sending and receiving profiles, supporting pathway enrichment, cell subtype discovery, multi-hop MI cascade detection, *in silico* spatial perturbation, and phenotype prediction.

SpiderNet implements this representation through two coupled components: directed cell-cell MI activity inference and molecular anchoring (**Fig**. 1, **Fig**. S1). Starting from spatially proximal cells, SpiderNet defines ordered sender-receiver pairs as candidate interactions and encodes each cell’s expression separately in sender and receiver roles. For each candidate interaction, SpiderNet uses the paired sender and receiver encodings to infer non-negative MI activities, yielding an edge-level map of MI activity across the tissue. A dual reconstruction objective then anchors these latent activities to their molecular programs. Edge-level MI activities reconstruct LR co-expression through MI-specific LR loadings, while outgoing and incoming activities are aggregated into cell-level sending and receiving factors, respectively. Gene expression is then reconstructed from these factors through MI-specific sender-regulator and receiver-target loadings, together with a separate cell-type-intrinsic component. Each MI therefore describes both where a communication program recurs and which sender regulators, LR pairs, and receiver targets characterize it.

The learned MI representation provides complementary spatial outputs at two resolutions. At cell-cell resolution, MI activities on directed neighboring pairs specify the location, direction, and strength of each communication program. At cellular resolution, sending and receiving MI profiles summarize each cell’s engagement across outgoing and incoming interactions, respectively (**Methods**). Together with the MI-specific molecular loadings, these outputs link spatial communication patterns to sender-side regulators, LR pairs, and receiver-side targets. This multiscale representation supports MI-associated pathway enrichment, MI-informed cell subtype discovery, multi-hop MI cascade detection, *in silico* spatial perturbation, and phenotype prediction. Details of model implementation, component ablations, MI-basis stability and robustness, and parameter selection are provided in the **Methods** and **Supplementary Notes**.

### SpiderNet recovers directed meta-interactions in simulation

To test recovery of a known modular communication structure, we simulated ST datasets with ground truth defined at both the directed cell-pair and molecular-program levels (**Fig**. 2a; **Methods**). We generated 2,000 cells from three cell types in two-dimensional space, with directed neighboring cell pairs along a ring assigned to MI-1, MI-2, or a non-interacting state. Gene expression was simulated to encode both cell-intrinsic identity and MI-dependent molecular programs, including MI-associated ligand-receptor pairs, sender regulators, and receiver targets. Additive noise and dropout were introduced to evaluate recoverability across ST-like noise and quality variation (**Supplementary Notes**).

**Figure 2:**
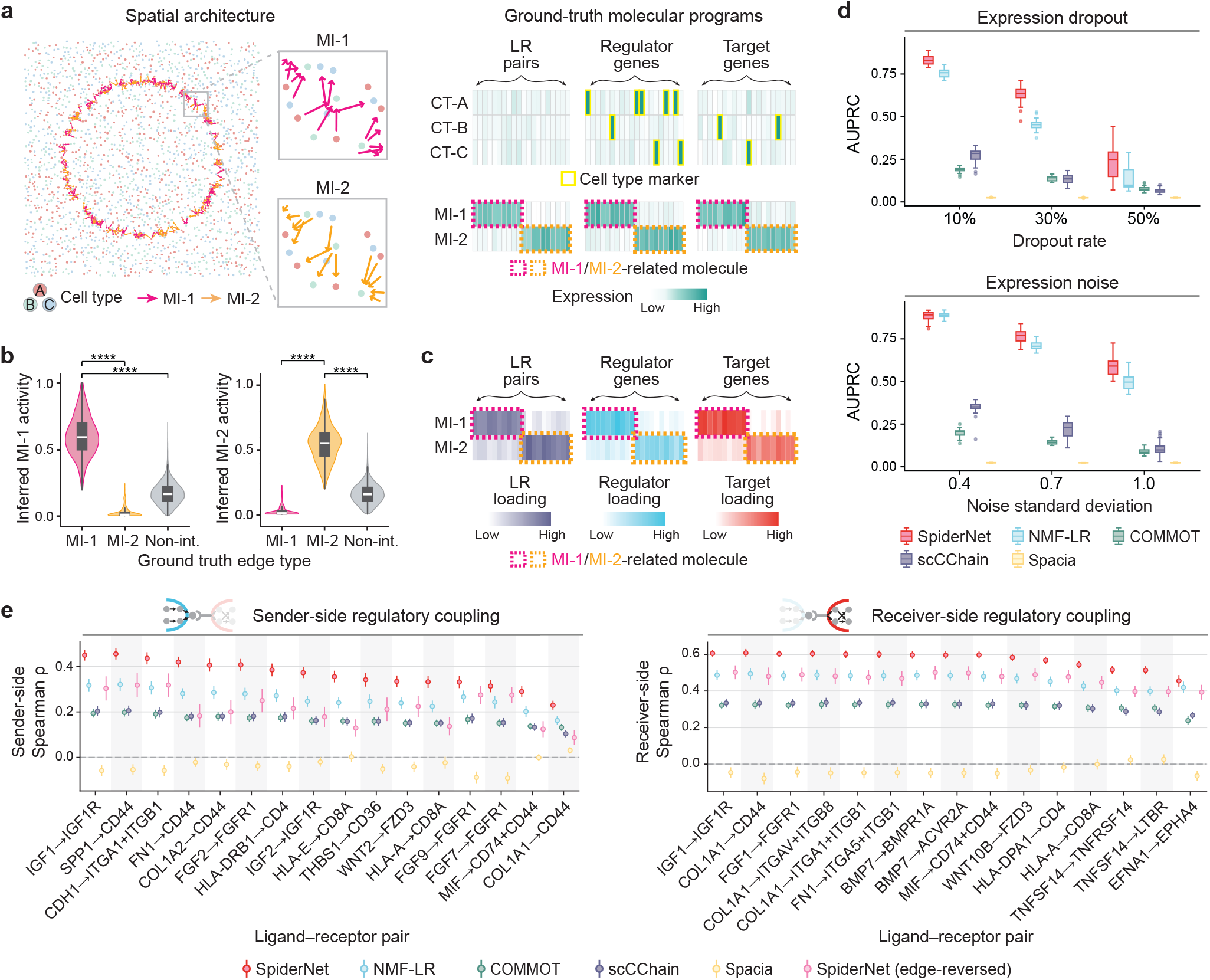
Recovery and real-tissue regulatory concordance of SpiderNet meta-interactions. **a**, Simulated spatial transcriptomics dataset comprising three cell types and two directed ground-truth meta-interactions (MIs). Candidate directed cell pairs along the ring were assigned to MI-1, MI-2, or a non-interacting state, and gene expression combined cell-type markers with MI-associated ligand-receptor (LR) pairs, sender regulators, and receiver targets. **b**, Inferred MI-1 and MI-2 activities stratified by ground-truth edge type. Inferred MIs were matched one-to-one to the ground-truth MIs for evaluation. Non-int., non-interacting pairs. *P* values are from two-sided Wilcoxon rank-sum tests; \*\*\*\**P<*0.0001. **c**, Recovered LR-pair, sender-regulator, and receiver-target loadings for the two MIs. Dashed outlines mark ground-truth MI-associated features. **d**, Recovery of ground-truth MI edges across dropout and noise settings, quantified by the area under the precision-recall curve (AUPRC) across 30 independent simulation replicates. **e**, Regulatory concordance of inferred communication features in the HGSOC CosMx dataset. Left, sender-side regulatory coupling, quantified as the correlation between sender-side communication activity and OmniPath [18] upstream-regulator activity. Right, receiver-side regulatory coupling, quantified as the correlation between receiver-side communication activity and scSeqComm [19] downstream-target activity. For each representative LR pair, points and error bars show the mean and 95% confidence interval across tissue slices, respectively. *SpiderNet (edge-reversed)* denotes a control that exchanges sender and receiver cells while retaining the original MI activity assigned to each edge.

SpiderNet accurately recovered two MIs corresponding to the ground-truth programs at both the cell-pair and molecular levels. Following one-to-one matching for evaluation, inferred MI activities were substantially higher on edges belonging to the corresponding ground-truth MI than on edges from the other MI or non-interacting pairs (**Fig**. 2b). Under the representative simulation setting shown (10% dropout), the median inferred activity was 0.59 (MI-1) and 0.55 (MI-2) on the corresponding ground-truth edges, compared with 0.01 for both MIs on edges from the other MI and 0.16 (MI-1) and 0.15 (MI-2) for non-interacting pairs. Furthermore, the ligand-receptor pairs, sender regulators, and receiver targets associated with each simulated MI were highly ranked in the learned loadings (**Fig**. 2c, **Fig**. S2). Across 30 replicates, median ratios of mean loading ranks for MI-associated versus non-associated features were 1.4-2.4 at 50% dropout and 1.9-2.8 at a noise standard deviation of 1.0 across LR, regulator, and target loadings. Ratios were significantly above 1 under every dropout and noise condition tested (**Fig**. S2). Thus, SpiderNet recovered both the participating cell pairs and each MI’s defining molecular signature.

We next benchmarked SpiderNet against four representative ST-based CCC methods, NMF-LR, COMMOT [11], scCChain [20], and Spacia [14], for recovery of the two ground-truth MI edge maps. To harmonize their distinct outputs, SpiderNet, NMF-LR, and scCChain were fitted with two latent components that were matched one-to-one to MI-1 and MI-2. COMMOT and Spacia LR- or gene-level edge scores were instead aggregated over the corresponding ground-truth LR or gene signature to produce MI-1- and MI-2-specific edge scores, giving these comparators access to the known molecular composition of each MI and representing oracle versions of these methods (**Supplementary Notes**). Across methods, SpiderNet more clearly distinguished true interacting from non-interacting pairs *in situ* (**Fig**. S3) and achieved the highest AUPRC for recovering ground-truth MIs across dropout and noise settings (**Fig**. 2d), with median AUPRCs of 0.56 and 0.79, respectively, versus 0.45 and 0.71 for the next-best method. At 50% dropout and at a noise standard deviation of 1.0, SpiderNet retained median AUPRCs of 0.24 and 0.59, respectively, compared with 0.10 and 0.49 for the next-best method. Its advantage persisted under the most challenging settings, supporting robustness to simulated sparsity and measurement noise.

Together, these controlled simulations show that SpiderNet recovers the spatial deployment of directed MIs and their molecular anchors from noisy, sparse ST-like data.

### Meta-interactions encode direction-specific regulatory structure in real tissues

Having established MI recoverability in simulation, we next asked whether MIs inferred from real tissues aligned with independently curated sender-to-receiver regulatory programs and whether this concordance depended on edge orientation. We evaluated these properties in two ST datasets profiled by distinct technologies: high-grade serous ovarian cancer (HGSOC) CosMx SMI [21] and aging mouse brain MERFISH [22].

For LR pairs with curated regulatory annotations, we tested whether inferred sender- and receiver-side communication activities aligned with activity scores for OmniPath-annotated upstream regulators and scSeqComm-annotated downstream targets, respectively (**Fig**. 2e, **Fig**. S4). Because these regulator and target annotations were not used during model training or CCC inference, this analysis provides an independent benchmark of regulatory concordance. To compare methods with different feature representations, we applied the same best-matching-feature procedure to every method and communication side (**Supplementary Notes**). Across both datasets, SpiderNet ranked first by regulatory-coupling score for all 28 sender-side and 36 of 37 receiver-side curated LR pairs, with mean increases of Δ*ρ*=0.11 and 0.09, respectively, over the next-best method. This indicates that the learned MI representation preserves biologically annotated sender-to-receiver regulatory structure.

To test orientation specificity, we constructed a post hoc edge-reversed control that retained each edge’s MI activity but exchanged its sender and receiver cells (**Supplementary Notes**). Edge reversal weakened both sender-side regulator coupling and receiver-side target coupling, with mean coupling-score decreases of Δ*ρ*=0.15 and 0.11, respectively, in HGSOC, and Δ*ρ*=0.21 and 0.19, respectively, in aging mouse brain (**Fig**. 2e, **Fig**. S4). Moreover, MI activities inferred for the two sender-receiver orientations of the same neighboring-cell pair were generally only weakly to moderately correlated within tissue slices (**Fig**. S5; **Supplementary Notes**). Together, these analyses support the directional specificity of the learned MIs.

SpiderNet communication states also preserved within-cell-type transcriptional similarity more strongly than comparison methods and showed higher MI activity in local neighboring pairs than in cell-type-matched distant cell pairs (**Fig**. S6, S7; **Supplementary Notes**). Taken together, these complementary regulatory, directional, and spatial controls support MIs as molecularly grounded, direction-specific sender-receiver programs with local tissue organization.

### An SPP1-THBS multicellular relay marks an immune-suppressive ovarian cancer niche

To examine whether the learned MI basis resolves malignant niches and their higher-order communication architecture within the tumor microenvironment (TME), we applied SpiderNet to the HGSOC CosMx SMI dataset [21] (276,668 cells from 48 untreated samples) (**Fig**. 3a). Using the MI dimensionality selection procedure (**Methods**), SpiderNet learned a 15-dimensional MI basis whose sender-regulator, LR-pair, and receiver-target signatures were supported by concordant molecular activity patterns (**Fig**. S8). These MIs were highly specific to sender-receiver cell-type pairs and aligned with canonical CCC pathways (**Fig**. 3b). Prominent malignant-cell-centered programs included an SPP1 monocyte-derived MI (MI-12), a THBS fibroblast-to-malignant MI (MI-10), and MHC-I malignant-to-T/NK MIs (MI-8 and MI-4), consistent with known roles of osteopontin/SPP1 signaling in tumor-associated myeloid niches [23, 24], THBS signaling in stromal adhesion and tumor progression [25, 26], and MHC-I-associated tumor-immune interactions [27, 28].

**Figure 3:**
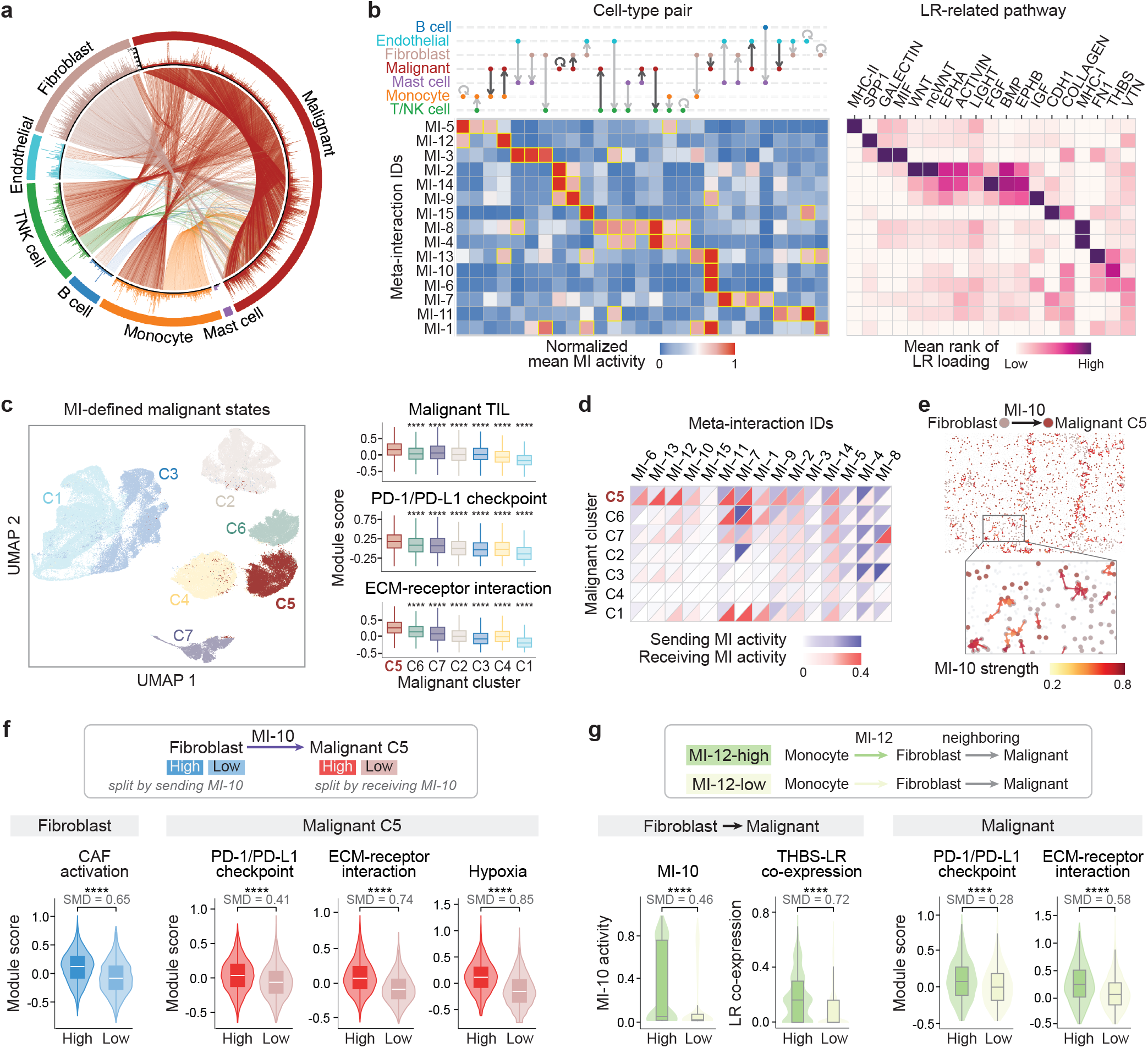
SpiderNet resolves an immune-suppressive CAF-malignant communication niche in HGSOC. **a**, Aggregated cell-cell MI map between non-tumor and malignant cells in the CosMx HGSOC dataset. Links indicate sender-receiver pairs with MI activity*>*0.7 in any MI dimension. **b**, MI landscape across cell-type pairs and LR-related pathways. Left, row-normalized MI activity across sender-receiver cell-type pairs; yellow boxes mark pairs involving malignant cells with normalized mean MI activity*>*0.7. Right, mean rank of LR loadings summarized by CellChat pathways for each MI. **c**, MI-defined malignant states and their functional programs. Left, UMAP of aggregated malignant-cell MI profiles, colored by Louvain cluster. Right, tumor-infiltrating lymphocyte (TIL), PD-1/PD-L1 checkpoint, and ECM-receptor interaction module scores across malignant clusters. C5 was compared with each other cluster using one-sided Wilcoxon rank-sum tests testing for higher scores in C5. **d**, Cluster-level sending and receiving MI activities across malignant clusters. **e**, Spatial distribution of fibroblast → MI-10 → malignant C5 interactions. **f**, CAF activation in fibroblasts and malignant immune-suppressive programs in C5 cells, stratified by MI-10 activity. Fibroblasts were stratified by sending MI-10 activity, whereas malignant C5 cells were stratified by receiving MI-10 activity. **g**, Topology-matched cascade analysis comparing MI-12-high monocyte-fibroblast-malignant triplets with MI-12-low baseline. Violin/box plots compare downstream MI-10 activity, THBS-related LR co-expression, and malignant immune-suppressive programs. For panels **f** and **g**, *P* values are from Wilcoxon rank-sum tests, and standardized mean differences (SMDs) are shown. Significance levels are denoted as \**P<*0.05; \*\**P<*0.01; \*\*\**P<*0.001; and \*\*\*\**P<*0.0001.

Clustering malignant cells by MI profiles identified seven malignant subtypes (C1-C7) (**Fig**. 3c; **Methods**) with distinct tumor-related transcriptional programs (**Fig**. S9a). We assessed their immune context using the malignant TIL (tumor-infiltrating lymphocytes) program, a malignant-cell signature of T/NK-cell infiltration defined in the original HGSOC study [21]. C5 had the highest malignant TIL score and also showed elevated PD-1/PD-L1 checkpoint, hypoxia, and ECM-receptor programs (**Fig**. 3c; **Fig**. S9b), identifying a TIL-associated yet immune-suppressive, ECM-remodeled malignant state [29, 30]. Concordantly, C5-neighboring CD8^+^ T cells and fibroblasts exhibited higher T-cell exhaustion and cancer-associated fibroblast (CAF) activation scores, respectively, than their non-C5-neighboring counterparts (**Fig**. S9c). Compared with Scanpy [31] and Banksy [32], MI-based subtyping was less dominated by sample identity and more clearly grouped cells with high immune-suppressive program scores (**Fig**. S9d).

C5 preferentially received three fibroblast-derived MIs corresponding to FN1 (MI-13), VTN (MI-6), and THBS (MI-10) signaling (**Fig**. 3b,d,e; **Fig**. S10a), all linked to elevated CAF activation in sender fibroblasts (**Fig**. S10b). Among them, THBS-associated MI-10 showed the strongest coupling to PD-1/PD-L1 checkpoint, ECM-receptor interaction, and hypoxia programs in malignant C5 cells (**Fig**. 3f; **Fig**. S10b). This prioritization is consistent with reported roles of THBS signaling in chemoresistance and immune suppression [33]. Compared with alternative CCC inference methods, SpiderNet showed the strongest joint coupling to CAF activation and malignant immune-suppressive programs (SMDs: 0.41-0.85; **Fig**. S10c; **Supplementary Notes**). Moreover, model-based perturbation of top MI-10 LR pairs reduced predicted CAF and C5 immune-suppression program scores relative to random LR perturbation, providing an internal consistency check for the MI-10 interpretation (**Fig**. S10d; **Methods**; **Supplementary Notes**).

We next asked whether this THBS-linked program was embedded within upstream TME signaling. MI-cascade analysis identified MI-12→MI-10 as an enriched monocyte→fibroblast→malignant relay among 18 significant ordered MI cascades (BH-adjusted empirical *P<*0.05; observed-to-expected ratio*>*1.5; **Fig**. S11a-c; **Methods**). This relay was detected in 28 of 48 HGSOC samples (**Fig**. S11c). The molecular signatures of MI-12 and MI-10 (**Fig**. 3b) defined a candidate SPP1-THBS immune-stromal-tumor cascade consistent with reported links among SPP1-high myeloid states, stromal remodeling, and immune-excluded TME [23, 24]. We compared MI-12-high triplets with topology-matched MI-12-low controls, which shared the same monocyte-fibroblast-malignant topology but had high versus low upstream MI-12 activity. MI-12-high triplets showed stronger downstream MI-10 activity (SMD=0.46) and THBS-related LR co-expression (SMD=0.72), accompanied by higher fibroblast CAF scores (SMD=1.08) and elevated malignant immune-suppressive programs (SMDs=0.28-0.58) (**Fig**. 3g; **Fig**. S11d).

Together, these analyses resolve an immune-suppressive CAF-malignant niche characterized by a THBS-linked fibroblast-malignant program coupled to an upstream SPP1-active monocyte-fibroblast program. Thus, the same MI representation links local malignant-cell states to higher-order communication across immune, stromal, and tumor cells.

### SpiderNet predicts T-cell responses to held-out melanoma perturbations

To test whether the MI representation links communication rewiring to cellular responses under direct genetic perturbation, we applied SpiderNet to the Perturb-FISH human melanoma dataset [34] (**Fig**. 4a). The dataset profiles CRISPR perturbations of 35 NF-*κ*B-related genes in melanoma cells, together with spatial transcriptomes of 154,418 melanoma and immune cells across 154 genes, enabling direct assessment of neighboring immune-cell responses to melanoma-cell perturbations. We focused on 11 perturbations previously reported to elicit strongly correlated transcriptional responses in neighboring T cells and evaluated SpiderNet’s ability to predict these non-cell-autonomous responses and resolve the underlying communication rewiring (**Fig**. 4a). Using the MI dimensionality selection procedure (**Methods**), SpiderNet learned a 23-dimensional MI basis spanning melanoma-T-cell communication across genetic perturbations.

**Figure 4:**
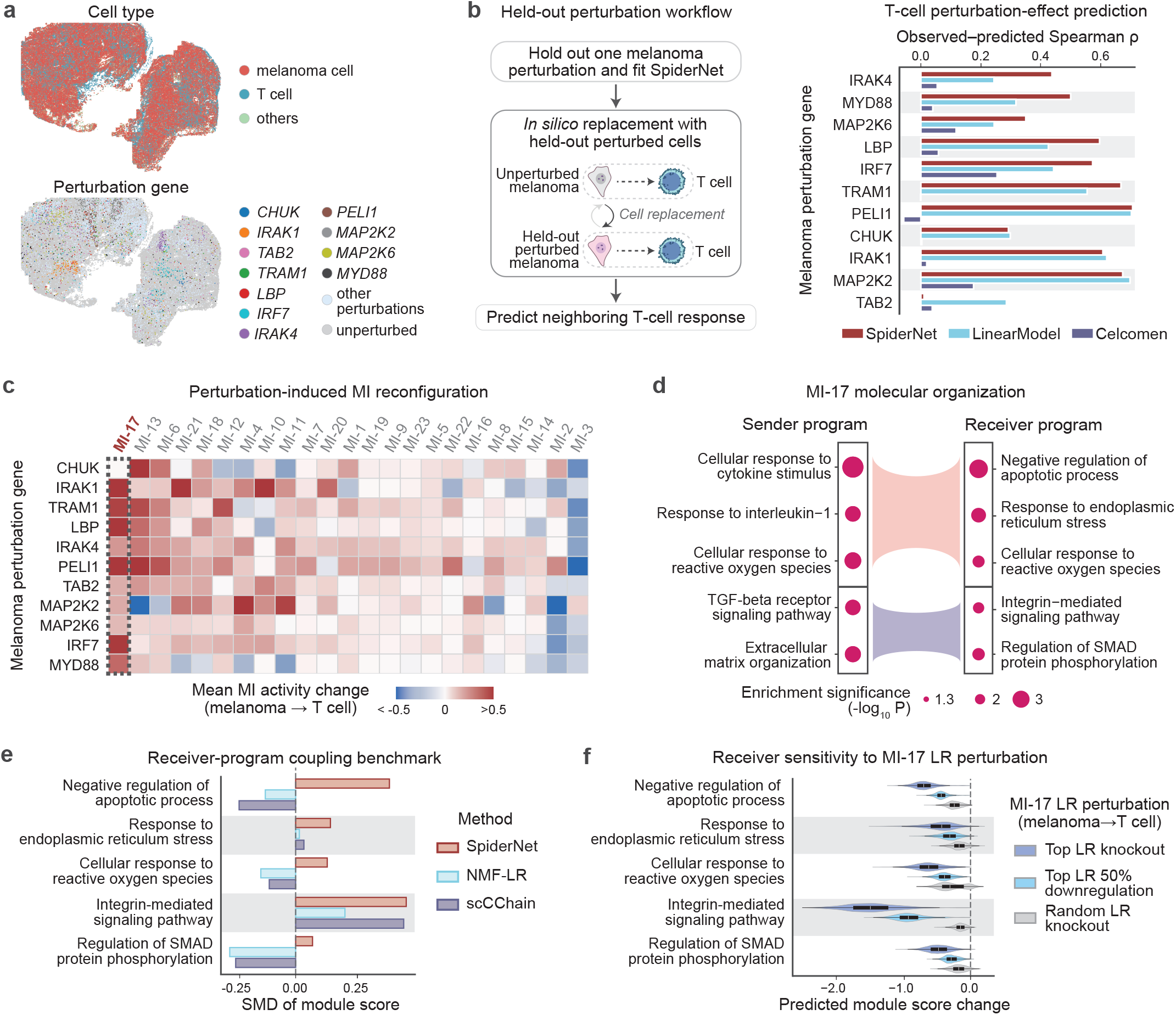
SpiderNet predicts held-out T-cell responses and resolves melanoma-to-T-cell crosstalk rewiring. **a**, Overview of the Perturb-FISH melanoma dataset, showing spatial cell-type annotations and CRISPR perturbation labels in melanoma cells for the 11 NF-*κ*B-associated perturbations analyzed. **b**, Held-out perturbation prediction. For each perturbation, SpiderNet was fitted after excluding melanoma cells carrying that perturbation; measured profiles of the held-out perturbed melanoma cells were then introduced by *in silico* replacement to predict responses in neighboring T cells. Bars show Spearman correlations between predicted and observed gene-wise T-cell responses for SpiderNet, the niche-summary linear model, and Celcomen. **c**, Mean perturbation-induced changes in melanoma-to-T-cell MI activity after *in silico* replacement. **d**, MI-17-associated melanoma sender and T-cell receiver programs. Dot size denotes enrichment significance, and curves indicate functional groupings linking sender and receiver biological processes captured by MI-17. **e**, Receiver-program coupling for each method’s most perturbation-responsive melanoma-to-T-cell communication feature. Coupling was quantified as the standardized mean difference (SMD) in module scores between T cells with high and low receiving activity; positive values indicate higher program scores in the high-activity group. **f**, Fixed-model sensitivity of MI-17 receiver programs in T cells to perturbation of top ligand-receptor (LR) pairs. Top pairs were knocked out or downregulated by 50% and compared with size-matched random LR-pair knockouts; negative values indicate reduced predicted program scores.

We tested predictive generalization using a leave-one-perturbation-out design. For each perturbation, the corresponding melanoma cells were excluded from training, and neighboring T-cell responses were predicted by replacing unperturbed melanoma cells with measured expression profiles from held-out perturbed melanoma cells while retaining the original tissue context (**Fig**. 4b, **Fig**. S12; **Supplementary Notes**). We compared SpiderNet with two niche-aware baselines: Celcomen [35] and a niche-summary linear model (**Supplementary Notes**). Across the 11 held-out predictions, SpiderNet achieved a median predicted-versus-observed T-cell response Spearman correlation of 0.57 versus 0.03 for Celcomen (paired two-sided Wilcoxon signed-rank test, *P* =0.002). Performance was broadly comparable to the niche-summary linear model, with SpiderNet achieving a higher median correlation (0.57 versus 0.42) and higher correlations in 7 of 11 perturbations. SpiderNet additionally resolved communication at directed cell-pair rather than aggregated niche resolution. Thus, the MI representation generalized to held-out perturbations and predicted their non-cell-autonomous effects on neighboring T-cell expression.

We then fitted SpiderNet to the full Perturb-FISH dataset to identify melanoma-T-cell communication programs remodeled across perturbations. Counterfactual MI-17 activity increased relative to the unperturbed state across all 11 perturbations and showed the largest mean increase in melanoma-to-T-cell interaction activity among the inferred MIs (mean MI activity change=0.35; **Fig**. 4c). Its receiver-target loadings were also enriched for T-cell response genes induced by melanoma NF-*κ*B perturbations (**Fig**. S13), nominating MI-17 as a prominent perturbation-responsive crosstalk program. In the observed data, melanoma-T-cell pairs involving perturbed melanoma cells likewise showed higher MI-17 activity, top ligand-receptor co-expression, and T-cell target-gene expression than pairs involving unperturbed melanoma cells (**Fig**. S14), supporting its perturbation responsiveness.

MI-17 linked two broad sender-receiver axes: melanoma cytokine and oxidative-stress programs with T-cell stress responses, and melanoma ECM/TGF-*β* programs with T-cell integrin and SMAD signaling (**Fig**. 4d; **Methods**), consistent with established roles of NF-*κ*B in inflammatory signaling and TGF-*β* in immune regulation [36–39]. MI-17 showed stronger receiver-side coupling to the five T-cell stress and SMAD-linked programs than the most perturbation-responsive programs from alternative CCC inference methods (mean SMD=0.23 versus −0.02 for the next-best method; **Fig**. 4e), while sender-side coupling was comparable (**Fig**. S15). MI-17 showed positive coupling across all five receiver programs, whereas the alternative methods showed weak or negative coupling for several. Moreover, *in silico* perturbation of the top MI-17 ligand-receptor pairs preferentially attenuated the corresponding melanoma sender and T-cell receiver program scores (**Fig**. 4f, **Fig**. S16), supporting the internal coherence of the inferred MI-17 molecular bridge.

Together, these analyses show that the MI representation predicts non-cell-autonomous T-cell responses to held-out melanoma perturbations and resolves perturbation-associated communication rewiring as a molecularly coherent melanoma-to-T-cell program.

### A T-cell-associated communication program tracks brain aging

Beyond perturbation-induced responses, we asked whether MI representation captures phenotype-associated communication variation that transfers across tissue contexts. We applied SpiderNet to an aging mouse brain MERFISH atlas [22] (712,695 cells, ten coronal sections spanning 3.4-33.2 months) to resolve age-associated remodeling of cell-cell communication (**Fig**. 5a). Using the MI dimensionality selection procedure (**Methods**), SpiderNet learned 30 MIs whose activities varied markedly with age (**Fig**. S17). Several MIs linked specific sender-receiver pairs to age-associated communication programs (**Fig**. S17, S18; **Methods**), including a complement-linked T-cell-derived MI (MI-29) and a VCAM-linked VSMC-to-immune MI (MI-12), consistent with reported immune and vascular aging programs [40–42].

**Figure 5:**
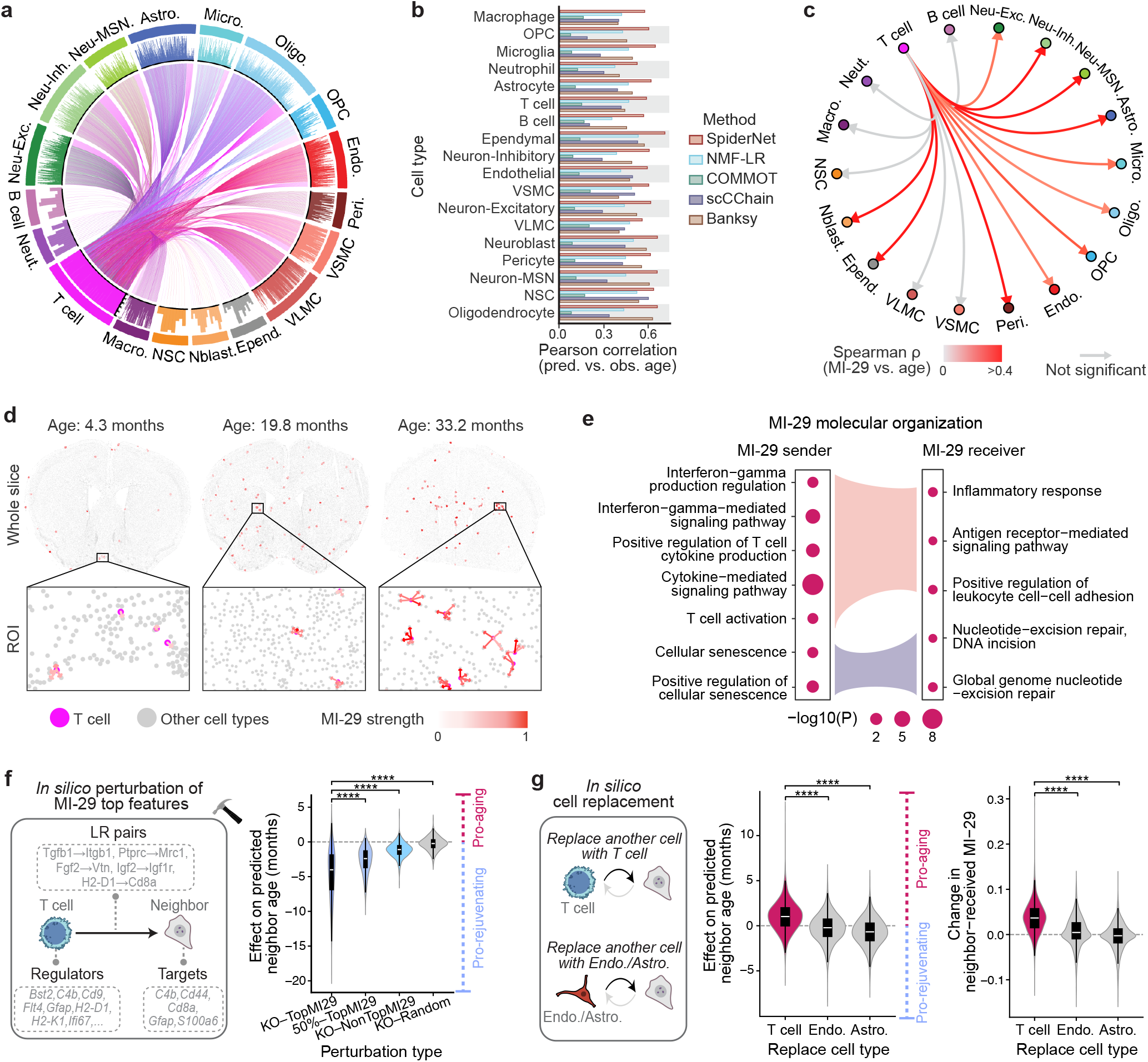
SpiderNet resolves an age-predictive MI landscape and a T-cell-associated aging program. **a**, Aggregated cell-cell MI map from T cells to neighboring cell types in the MERFISH aging brain dataset. Links indicate sender-receiver pairs with MI activity*>*0.7 in any MI dimension. **b**, Cell-level age prediction benchmark across 18 cell types in the coronal MERFISH atlas using SpiderNet, NMF-LR, COMMOT, scCChain, and Banksy representations, measured by out-of-fold Pearson correlation between predicted and observed age under age-stratified 10-fold cross-validation. **c**, Association of T-cell-derived MI-29 activity with tissue-section age across receiver cell types, quantified by Spearman correlation between received MI-29 activity and section age. Gray denotes non-significant correlations with two-sided Spearman test *P*≥0.05. **d**, *In situ* MI-29 interactions from T cells to neighbors across ages. Top: whole-slice views. Bottom: zoomed views at 4.3, 19.8, and 33.2 months. **e**, GO enrichment of MI-29-associated sender regulators (left) and receiver targets (right). Curves indicate functional groupings linking sender and receiver sides, and dot size denotes enrichment significance. **f**, Fixed-model *in silico* perturbation of MI-29-associated LR pairs, sender regulators, and receiver targets in T-cell→neighbor interactions. Violin plots show changes in model-predicted age of neighboring cells. “KO-TopMI29”, MI-29 top-feature knockout (KO); “50%-TopMI29”, 50% downregulation of MI-29 top features; “KO-NonTopMI29”, KO of MI-29 non-top features; and “KO-Random”, size-matched random-feature KO. **g**, Fixed-model *in silico* cell replacement with T cells, astrocytes, or endothelial cells, showing changes in model-predicted neighbor age and received MI-29 activity. For panels **f**,**g**, *P* values are from two-sided Wilcoxon rank-sum tests for the indicated comparisons; \*\*\*\**P<*0.0001.

To test whether the learned MI basis captured aging-relevant variation, we trained cell-type-specific linear models to predict chronological age from SpiderNet-derived cell-level MI profiles (**Supplementary Notes**). Using the same age-stratified 10-fold cross-validation scheme, we compared SpiderNet with NMF-LR, COMMOT [11], scCChain [20], and the spatial-niche-based method Banksy [32]. SpiderNet achieved the highest predicted-versus-observed age correlation in all 18 evaluated cell types, with all correlations exceeding 0.5 (**Fig**. 5b; **Fig**. S19). To test cross-context transfer, we applied the coronal-trained age-prediction models without retraining to sagittal sections from the same atlas [22] and to an independent aging hippocampus Stereo-seq dataset [43], using the 300 genes shared with the MERFISH panel. Across both settings, SpiderNet consistently preserved the expected old-versus-young ordering of predicted age and showed stronger old-versus-young separation than the baseline methods (median old-to-young predicted-age ratio: 1.16 versus 1.06 for the next-best method in sagittal sections and 1.40 versus 1.02 in hippocampus Stereo-seq; **Fig**. S20a; **Supplementary Notes**). Thus, MI-derived age signals transferred across anatomical regions and ST technologies.

MI-29 emerged as a prominent component of this age-predictive landscape. It was upregulated in older brain sections (**Fig**. S17, left), contributed positively to age prediction across cell types (**Fig**. S21a), and predominantly captured T-cell-to-neighbor interactions (**Fig**. S21b). Received MI-29 activity was positively associated with tissue-section age in 11 of 17 neighboring cell types, with all significant correlations exceeding *ρ*=0.25 and five exceeding *ρ*=0.4 (**Fig**. 5c,d). This pattern is consistent with the reported association between T-cell proximity and local aging signatures in this atlas [22].

Gene Ontology enrichment of MI-29-associated sender regulators and receiver targets linked T-cell interferon-gamma signaling and activation to inflammatory responses in neighboring cells (**Fig**. 5e; **Methods**). These programs were consistent with reported immune-inflammation and aging-associated stress programs [22, 44]. Concordantly, high MI-29-sending T cells showed elevated aging module scores in the coronal atlas (**Fig**. S21c), with a similar pattern observed in sagittal sections from the same atlas (**Fig**. S20b). Compared with the most aging-score-coupled programs from alternative CCC representations, MI-29 showed the strongest association with the T-cell aging score (**Fig**. S21d). Together, these results identify MI-29 as a molecularly anchored T-cell-to-neighbor program linking immune activation to aging-associated transcriptional variation in neighboring cells.

To test the model-predicted sensitivity of aging signals to molecular and cellular changes in MI-29, we performed fixed-model gene perturbation and cell-replacement analyses (**Methods** and **Supplementary Notes**). Knockout or partial downregulation of MI-29-associated LR pairs, sender regulators, and receiver targets in T-cell-to-neighbor interactions reduced predicted neighbor-cell age, with a significantly larger decrease following knockout of top MI-29-associated genes than random or non-top genes (**Fig**. 5f). Replacing local cells with tissue-sampled T cells increased both predicted neighbor age and neighbor-received MI-29, whereas replacement with astrocytes or endothelial cells had negligible effects (**Fig**. 5g).

Collectively, these analyses show that the MI basis captures transferable age-associated communication variation and prioritizes a molecularly anchored T-cell-to-neighbor program whose molecular and cellular components influence predicted aging signals within the fitted model.

### Pan-cancer meta-interactions reveal a recurrent COLLAGEN fibroblast-tumor program

Pan-cancer studies have identified recurrent tumor microenvironment (TME) states with context-dependent cellular composition and clinical associations [45, 46]. How these states are encoded by spatial communication programs across cancers remains unclear. We applied SpiderNet to the publicly available MERSCOPE FFPE Human Immuno-Oncology atlas from Vizgen (see **Data availability**), comprising 4,707,751 cells from large-area tumor sections spanning colon, liver, melanoma, ovarian, prostate, lung, breast, and uterine cancers (**Fig**. 6a; **Supplementary Notes**). Joint modeling yielded a shared MI basis for comparing recurrent tumor-associated programs, their molecular signatures, clinical associations, and higher-order relays.

**Figure 6:**
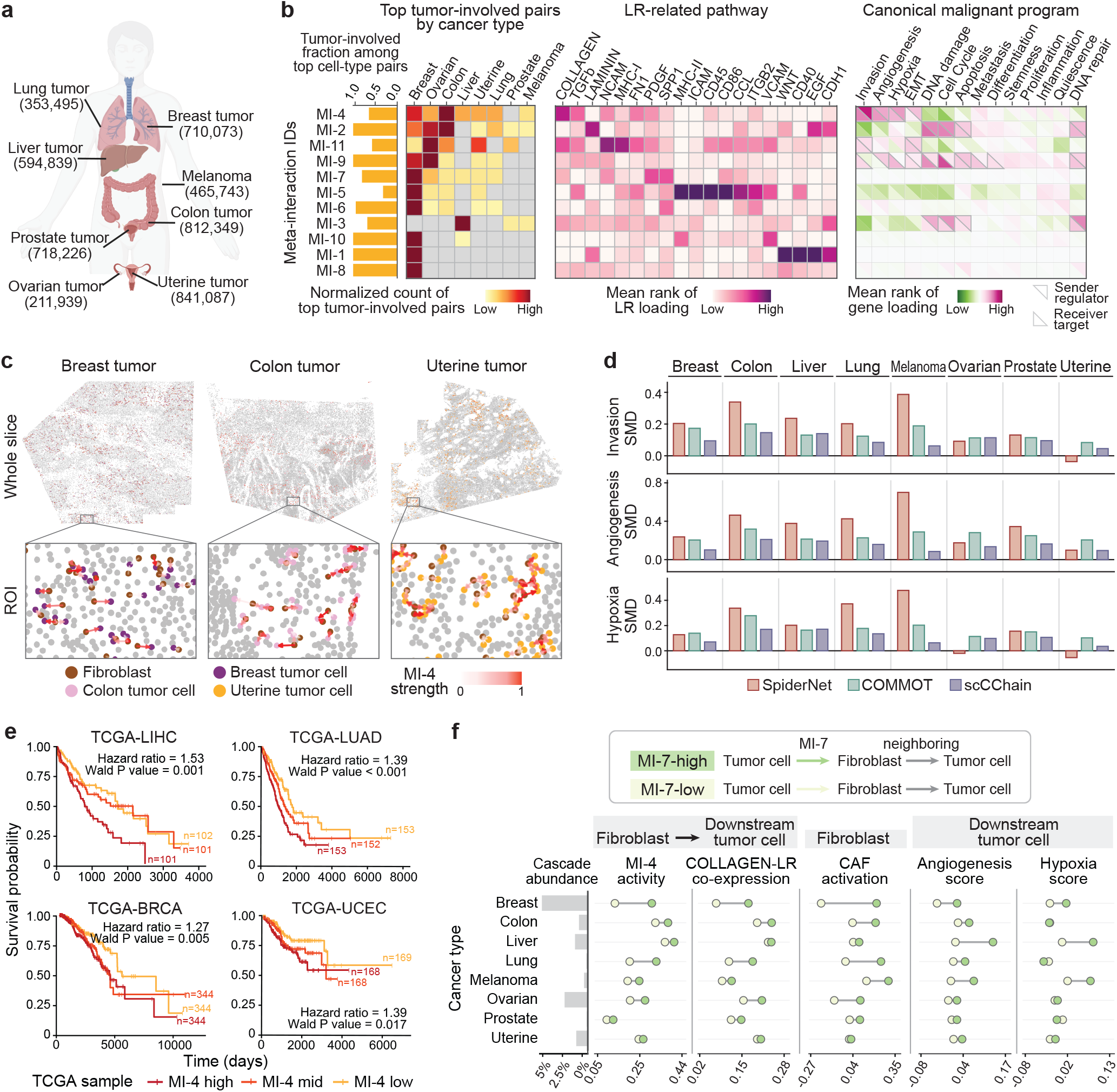
SpiderNet identifies a recurrent COLLAGEN-linked fibroblast-tumor MI associated with adverse survival and embedded in a multi-hop relay. **a**, Pan-cancer MERSCOPE FFPE Human Immuno-Oncology atlas comprising 4,707,751 cells across eight cancer types. **b**, Jointly learned MI basis. Left, fraction of the top 10% sender-receiver cell-type pairs involving tumor cells for each MI; middle, recurrence of top tumor-involved pairs across cancer types; right, mean loading ranks summarized by CellChat LR pathways and canonical malignant programs, with sender-regulator and receiver-target components shown separately. **c**, *In situ* fibroblast→tumor MI-4 interactions in representative breast, colon, and uterine tumor sub-slices. **d**, Malignant-program coupling benchmark across CCC methods. Bars show standardized mean differences (SMDs) in invasion, angiogenesis, and hypoxia between high- and low-communication tumor cells; larger positive SMDs indicate higher malignant-program scores in high-communication tumor cells. Each baseline uses its most strongly coupled fibroblast→tumor communication feature, whereas SpiderNet uses MI-4. **e**, Kaplan-Meier curves for overall survival in the four representative TCGA cohorts with significant adverse MI-4 survival associations, stratified by bulk-projected MI-4 abundance. Hazard ratios and Wald *P* values are from multi-variable Cox models adjusted for age, sex, and tumor stage. **f**, Pan-cancer MI-7→MI-4 relay analysis. Left, abundance of tumor→fibroblast→tumor MI-7→MI-4 cascades across cancer types. Right, topology-matched MI-7-high versus MI-7-low comparisons of median downstream MI-4 activity, COLLAGEN ligand-receptor co-expression, fibroblast CAF activation, angiogenesis, and hypoxia.

SpiderNet resolved TME communication into an 11-dimensional shared MI basis with distinct tumor involvement and recurrence patterns. The strongest sender-receiver cell-type pairs for each MI showed that most MIs were dominated by tumor-involved interactions, with recurrence ranging from broadly shared to context-specialized across cancer types (**Fig**. 6b, left and middle). Integrating CancerSEA enrichment of sender-regulator and receiver-target loadings with CellChat pathway summaries of LR loadings linked each MI to malignant programs and intercellular signaling contexts (**Fig**. 6b, right; **Methods**). For example, MI-4 was anchored by COLLAGEN signaling and aligned with invasion, angio-genesis, and hypoxia, consistent with established links between ECM remodeling, hypoxic tumor niches, and metastatic progression [47], whereas MI-2 linked LAMININ/EGF signaling to stress-response programs consistent with reported ECM/integrin-associated tumor-cell survival responses [48].

Within this shared basis, MI-4 emerged as the most recurrent tumor-involved program across cancer types (**Fig**. 6b) and primarily captured fibroblast-tumor communication (**Fig**. 6c, **Fig**. S23). Tumor cells with high incoming fibroblast→tumor MI-4 activity showed higher invasion, angiogenesis, and hypoxia scores in 21 of 24 cancer-type-by-program comparisons (**Fig**. S24a). Compared with the most strongly coupled fibroblast→tumor programs from alternative CCC representations, SpiderNet MI-4 ranked first for malignant-program coupling in 17 of 24 comparisons versus 6 for the next-best method (**Fig**. 6d; **Supplementary Notes**). Moreover, *in silico* perturbation of the top MI-4 LR pairs attenuated predicted invasion, angiogenesis, and hypoxia scores in receiver tumor cells relative to random perturbations (**Fig**. S24b), indicating model-predicted sensitivity of these malignant programs to MI-4-associated LR features.

We next asked whether the recurrent fibroblast-tumor MI-4 program was linked to survival and immunotherapy response. We projected the spatially learned MI basis onto TCGA bulk transcriptomes to estimate sample-level MI abundances using fixed MI molecular signatures. Pseudo-bulk validation showed that the projection preserved the relative ordering of sub-slice-averaged MI abundances (**Fig**. S25; **Supplementary Notes**). Cox regression models adjusted for age, sex, and stage associated higher projected MI-4 abundance with adverse survival in four of nine TCGA cohorts: TCGA-LIHC, TCGA-LUAD, TCGA-BRCA, and TCGA-UCEC (hazard ratios: 1.27-1.53; all Wald *P<*0.05; **Fig**. 6e; **Fig**. S26a; **Supplementary Notes**). Moreover, nested Cox likelihood-ratio tests showed that MI-4 added prognostic information beyond clinical covariates and MI-4-linked invasion, angiogenesis, and hypoxia programs in all four adverse-survival cohorts (**Fig**. S26b). We further applied bulk MI projection to three published immune-checkpoint blockade (ICB)-treated melanoma and lung cancer cohorts [49– 51]. Projected MI-4 abundance was higher in ICB non-responders than in responders across all four cohort-treatment settings and ranked first for non-response association among the evaluated MIs and established immunotherapy biomarkers when ranks were averaged across settings (**Fig**. S26c; **Supplementary Notes**). These independent-cohort associations are consistent with prior links between CAF-associated collagen/ECM programs, adverse tumor phenotypes, and checkpoint blockade failure [30, 46], supporting the potential clinical relevance of the projected MI-4 fibroblast-tumor program.

We next asked whether the cross-cancer recurrence of this COLLAGEN-linked fibroblast→tumor program extended to a multi-hop relay. MI-cascade analysis identified an MI-7→MI-4 cascade that was significantly enriched in 7 of 8 cancer types (BH-adjusted empirical *P<*0.05; observed-to-expected cascade-count ratio*>*1.5; **Methods**) and recurrently formed tumor→fibroblast→tumor triplets (**Fig**. S27-S28). This topology is consistent with reciprocal tumor-CAF signaling and CAF-mediated matrix remodeling [52, 53]. Compared with topology-matched MI-7-low controls, MI-7-high triplets showed stronger downstream MI-4 activity and COLLAGEN ligand-receptor co-expression, accompanied by higher fibroblast CAF activation scores and elevated angiogenesis and hypoxia scores in downstream tumor cells (**Fig**. 6f). Thus, MI-7→MI-4 delineates a reciprocal tumor-stromal relay linking upstream tumor→fibroblast signaling to the recurrent COLLAGEN fibroblast→tumor program, CAF activation, and malignant remodeling.

Together, these analyses identify a recurrent COLLAGEN-linked fibroblast-tumor MI associated with malignant states, adverse survival, and immunotherapy non-response, and place it within a reciprocal tumor-fibroblast-tumor relay. More broadly, the shared MI basis distinguishes recurrent communication architecture from its context-specific transcriptional and clinical associations across cancers.

## Discussion

A central challenge in tissue communication is to identify the scale at which molecular signaling gives rise to coordinated multicellular regulation. Across the biological settings examined here, recurrent, directed cell-pair programs emerged as an intermediate scale coupling sender regulation, LR signaling, and receiver response. SpiderNet represents this scale as a compact basis of MIs, each defined by a sender-LR-receiver signature and cell-pair-resolved activity across tissue. By learning communication programs together with their molecular signatures and spatial activities, SpiderNet unifies biological interpretation and prediction within the same representation. Simulations and real-tissue benchmarks established MI recoverability and direction-specific regulatory concordance, respectively, while component ablations supported the value of joint molecular anchoring and robustness analyses demonstrated the stability of the learned MI basis. Across applications, MIs recurred across contexts, assembled into multicellular relays, and linked local communication to cell states, tissue phenotypes, and non-cell-autonomous perturbation responses. Together, these findings support MIs as biologically meaningful intermediate units that integrate molecular, cellular, and spatial dimensions of tissue communication.

Biological applications highlight three broader principles. First, pairwise MIs can be linked into higher-order multicellular relays associated with downstream cell states, as illustrated by the SPP1-THBS immune-stromal-tumor relay in HGSOC and the reciprocal tumor-fibroblast-tumor relay across cancers. In ovarian cancer, the THBS-linked niche combines signatures of lymphocyte infiltration, checkpoint activity, and neighboring T-cell exhaustion. The SPP1-THBS relay places this TIL-rich yet immune-suppressive niche within a wider myeloid-stromal signaling context. Second, MIs connect local communication to cell state and response: melanoma MIs predicted neighboring T-cell responses to held-out NF-*κ*B perturbations, whereas brain MIs identified a T-cell-associated aging program and captured age-predictive signals that transferred across anatomical regions and platforms. Third, recurrent MI programs can be deployed across heterogeneous tissue contexts, exemplified by the pan-cancer COLLAGEN-linked fibroblast-tumor program. More broadly, recurrence and phenotypic association are distinct aspects of a communication program: the shared MI basis identifies programs that recur across tumors while resolving their context-specific cellular and clinical associations. Together, these findings suggest that MIs are reusable organizational units linking local signaling to higher-order multicellular organization and context-dependent states.

Several limitations remain in this representational framework. First, MI directionality reflects ordered sender-receiver asymmetry rather than causal signal flow. Although MI directionality is informed by LR ordering and asymmetric sender- and receiver-side regulatory programs, static ST data alone cannot establish temporal or biochemical causality. Incorporating pathway and gene-regulatory priors [54, 55] together with perturbation-calibrated data [56] could strengthen the mechanistic interpretation of MIs. Second, the current formulation primarily captures local LR-mediated communication, whereas tissue regulation can also involve longer-range, contact-independent, and non-LR mechanisms [2]. Extending the model across molecular channels and spatial scales could broaden its coverage of tissue communication. Third, MI resolution depends on transcript coverage, available LR annotations, and RNA abundance as an indirect readout of functional signaling. Targeted ST panels may omit communication-relevant genes, while transcript levels may not directly reflect signaling activity. Integration with whole-transcriptome references [57] and spatial multi-omics [58] could improve molecular resolution. Finally, *in silico* perturbation analyses estimate model-predicted sensitivity rather than experimental causal effects, whereas projection of spatially learned MIs into bulk cohorts yields sample-level program abundance without preserving cell-pair resolution. Experimental perturbation of prioritized MI bridges and prospective validation of projected clinical associations will therefore be important for establishing causality and translational relevance.

Overall, SpiderNet reframes spatial communication analysis from cataloging candidate molecular contacts to resolving recurrent, directed MI programs. This intermediate representation complements LR- and pathway-level analyses by organizing molecular signals into reusable programs linked to cell states, multicellular relays, perturbation responses, and tissue phenotypes. As spatial profiling becomes increasingly multimodal and perturbation-rich, MI-based models provide a foundation for comparing, tracing, and experimentally testing how communication programs shape multicellular behavior across biological contexts.

## Methods

### The SpiderNet model

SpiderNet takes as input spatial transcriptomics data comprising each cell’s gene-expression profile, spatial coordinate, and a pre-annotated cell-type label, together with curated ligand-receptor (LR) pairs from public databases [9, 19]. Let 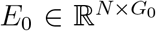 denote the raw transcript-count matrix for *N* cells and *G*_0_ genes. Counts are normalized by each cell’s total count, scaled to 10,000, and log1p-transformed. The top *G* highly variable genes are selected using Scanpy [31], yielding the processed expression matrix *E* ∈ ℝ ^*N* ×*G*^.

To incorporate spatial context, we perform a directed spatial *k*-nearest-neighbor (*k*NN) search by connecting each cell to its k nearest neighbors without symmetrization, defining the resulting edges ***E***= {(*i*→*j*)} as candidate sender-receiver interactions. This directed formulation allows the two directions between the same pair of neighboring cells to carry different communication activities. Let 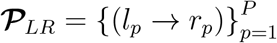 denote the curated LR pairs, where *l*_*p*_ and *r*_*p*_ are the ligand and receptor genes or complexes of the *p*-th LR pair. For each directed neighboring cell pair (*i* → *j*) and LR pair (*l*_*p*_, *r*_*p*_), LR co-expression is defined as:

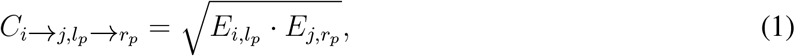

which is the geometric mean of ligand expression in sender cell *i* and receptor expression in receiver cell *j*. For multi-subunit ligands or receptors, we use the arithmetic mean expression of constituent genes present in the input matrix.

#### Encoding directed cell pairs into MI activities

SpiderNet infers *M* latent meta-interaction (MI) dimensions 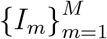, each representing a coordinated directional communication module that links sender-side regulators, LR bridges, and receiver-side target genes. For each directed neighboring pair (sender *i* → receiver *j*), gene expression is encoded by sender- and receiver-specific multilayer perceptrons (MLPs), producing sender and receiver embeddings 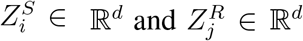. The two role-specific embeddings are concatenated into a pairwise sender-receiver representation:

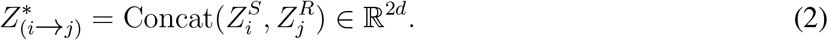

This pairwise representation is passed through a scoring MLP to infer non-negative MI activities:

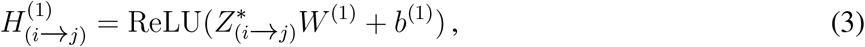

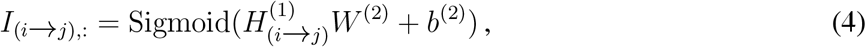

where *I*_(*i*→*j*),*m*_ ∈ [0, 1] denotes the activity of the *m*-th MI from sender cell *i* to receiver cell *j*, and *W* ^(*l*)^ and *b*^(*l*)^, for *l* = 1, 2, are trainable weights and biases.

#### Aggregating pairwise MIs into cell-level sending and receiving profiles

Inferred pairwise MI activities are summarized into cell-level sending and receiving profiles. For each cell *i*, let 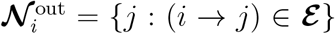 and 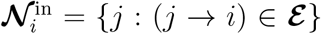 denote its outgoing and incoming neighborhoods, respectively. MI activities are averaged over the corresponding directed pairs:

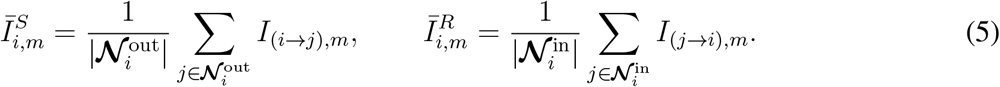

Here, 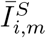 quantifies how strongly cell *i* sends MI *m* to its neighbors, whereas 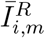 quantifies how strongly cell *i* receives MI *m* from its neighbors. These aggregated profiles provide cell-level representations of local communication state and are used for gene-expression reconstruction and downstream analyses.

#### Interpretable reconstruction of cellular gene expression

To connect inferred MIs to intracellular transcriptional programs, SpiderNet reconstructs the normalized expression of each cell as the sum of three interpretable components:

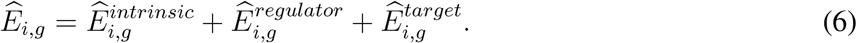

The intrinsic term captures baseline expression associated with annotated cell identity, whereas the regulator- and target-associated terms model variation associated with the aggregated sending and receiving MI profiles, respectively:

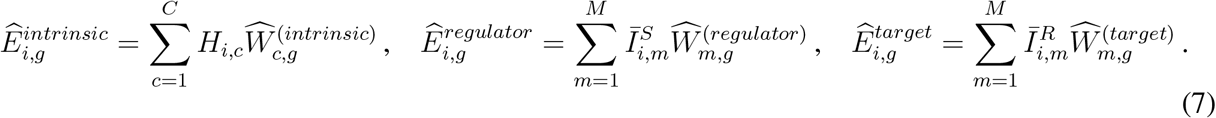

Here, *H* ∈ ℝ*N* ×*C* is the one-hot encoded cell-type matrix, and 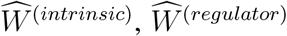, and 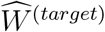 are non-negative gene-loading matrices learned during training. The regulator and target loading matrices link each MI to genes associated with the sender and receiver sides of the inferred communication module, respectively.

#### Interpretable reconstruction of ligand-receptor co-expression

In parallel with gene-expression reconstruction, SpiderNet uses pairwise MI activities to reconstruct LR co-expression for each directed neighboring cell pair. For sender cell *i*, receiver cell *j*, and LR pair (*l*_*p*_ → *r*_*p*_), the reconstructed LR co-expression is

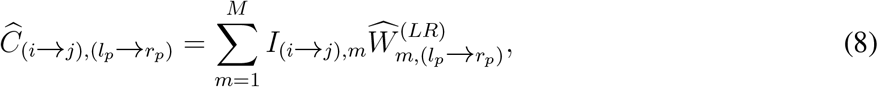

where 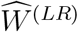 is a learnable non-negative loading matrix that quantifies the contribution of each LR pair to each MI. Together, 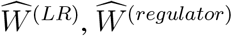, and 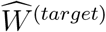 provide molecular anchors for each MI, linking sender-side regulators, LR bridges, and receiver-side target genes.

#### Joint optimization

SpiderNet is trained end-to-end by jointly reconstructing cellular gene expression and directed LR co-expression:

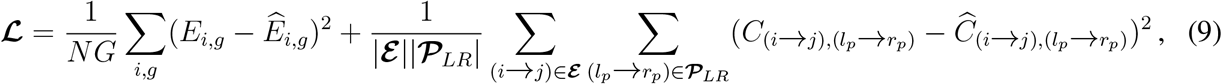

where ***ε*** denotes the set of directed neighboring cell pairs, and |***E***| and |***P***_*LR*_| denote the numbers of neighboring cell pairs and LR pairs, respectively. Each reconstruction term is normalized by its number of entries, and the two terms are assigned equal weights because gene expression and LR co-expression are derived from expression values on the same normalized scale. The model is trained for 20,000 epochs by default using the Adam optimizer with section-level batching and gradient accumulation. The learning rate was set to 5 × 10^−4^ during the initial 10% of training and to 1 × 10^−4^ thereafter, with ReduceLROnPlateau scheduling; weight decay was fixed at 1 × 10^−5^.

### SpiderNet supports diverse downstream applications

SpiderNet outputs cell-cell MI activities and associated loading matrices, which support diverse down-stream applications (**Fig**. 1).

#### MI-associated pathway enrichment

SpiderNet quantifies the contribution of sender regulators, receiver targets, and LR pairs to each MI dimension using the non-negative loading matrices 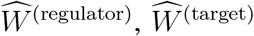, and 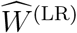. To measure the specificity of each gene and LR pair across MIs, we apply gene-wise and LR-wise normalization:

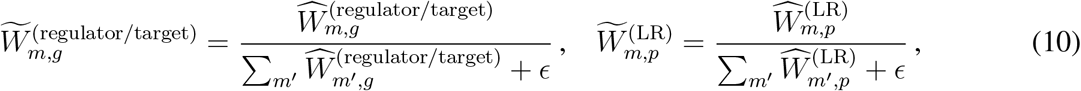

with *ϵ* = 1 × 10^−6^. These normalized loadings reflect how specifically each gene or LR pair associates with a given MI. For each MI *I*_*m*_, genes and LR pairs are ranked by 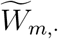, and the top-ranked features are reported as key molecular components.

By default, MI-associated sender regulators and receiver targets are defined as genes with MI-specific normalized loadings greater than 0.2. Gene Ontology biological process enrichment is performed independently for sender- and receiver-side gene sets, and enriched terms are considered significant at Benjamini-Hochberg-adjusted *P<*0.05.

#### MI-informed cell-subtype identification

To derive MI-informed cell subtypes, we use the aggregated MI activities 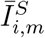 and 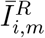 for each cell according to Eq. 5. We define the corresponding sending and receiving MI vectors as 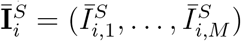 and 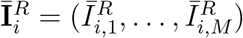, respectively. The aggregated MI profile of each cell is obtained by concatenating these two vectors:

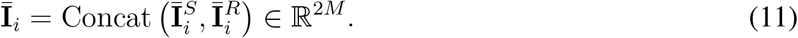

Within each annotated cell class, a *k*-nearest-neighbor graph (*k*=25) was constructed from the aggregated MI profiles using Euclidean distance, followed by Louvain clustering to identify finer-grained MI-related subtypes.

#### In silico spatial perturbation

To examine how model-predicted communication and phenotypes respond to molecular perturbations and changes in local cellular composition, we performed two types of *in silico* perturbations using the trained SpiderNet model. First, **gene perturbation** was implemented by setting the expression of selected genes to zero or partially reducing their expression in specified sender or receiver cells. Second, **cell replacement** was implemented by replacing the expression profile at a given spatial location with that of a tissue-sampled cell from a different cell state, while preserving the original spatial coordinates. In both analyses, the trained SpiderNet model is held fixed and used to recompute MI activities, model-predicted expression, and downstream phenotype scores. Changes in these outputs indicate the sensitivity of the inferred communication architecture and associated phenotypes to specific molecular features and local cell states within the fitted model.

#### MI-cascade detection

Let 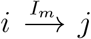 denote an edge from sender cell *i* to receiver cell *j* mediated by MI *I*_*m*_. An **MI-cascade**, defined by an ordered MI pair (*I*_*m*1_, *I*_*m*2_), is any triplet of cells (*i, j, k*) satisfying 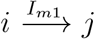 and 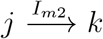 (**Fig**. 1), representing a sequential two-hop communication motif across three cells. To detect such cascades, we first retain strong MI edges with activity*>*0.6 and then, for each ordered MI pair (*I*_*m*1_, *I*_*m*2_), count all three-cell chains 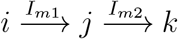 in tissue.

A null distribution of cascade chain counts is obtained by randomly reassigning MI edges among observed cell pairs within each sender-cell-type by receiver-cell-type stratum, recounting cascades, and repeating 100 times. This preserves the observed spatial edges and cell-type-pair composition while disrupting the higher-order ordering of MI edges. For each ordered MI pair, empirical *P* values were calculated as (*b* + 1)*/*(*B* + 1), where *B* = 100 and *b* is the number of permutations with cascade counts at least as large as the observed count. Empirical *P* values are then adjusted across all ordered MI pairs using the Benjamini-Hochberg procedure. Cascades are considered enriched when the BH-adjusted empirical *P<*0.05 and the observed-to-expected count ratio*>*1.5.

#### Phenotype prediction

Aggregated MI sending and receiving activities (Eq. 5) are used to predict cellular phenotypes. For the aging-brain analysis, a cell-type-specific linear regression model is trained using the concatenated sending and receiving MI profiles as predictors and chronological age as the response. Performance is evaluated by the correlation between out-of-fold predicted and observed ages using age-stratified 10-fold cross-validation at the cell level, with cells from each tissue section distributed across folds.

### Initialization

Initial model parameters are obtained as follows. First, we apply non-negative matrix factorization (NMF) to the precomputed cell-cell LR co-expression matrix, with the number of NMF factors set to the MI dimensionality *M*, yielding initial estimates of the cell-cell MI activity matrix *I* and LR loading matrix *W* ^(*LR*)^. Each NMF activity factor was max-normalized across cell-cell edges before encoder pretraining. NMF is implemented using `sklearn.decomposition.NMF`with NNDSVD initialization [59]. Second, aggregated sending and receiving MI activities (*Ī*^*S*^ and *Ī*^*R*^; Eq. 5) are concatenated with the one-hot encoded cell-type matrix *H* and used as covariates in a non-negative linear regression to predict the gene-expression matrix *E*, providing initial estimates of the gene loading matrices (*W* ^(intrinsic)^, *W* ^(regulator)^, and *W* ^(target)^). Finally, the encoder is pretrained to minimize the mean squared error between the NMF-derived MI activity matrix *I* and the encoder output. All model parameters are subsequently optimized jointly using the objective in Eq. 9.

### Selection of major tuning parameters

SpiderNet has two major tuning parameters: (1) the MI dimensionality and (2) the spatial neighborhood size (*K*) used to define candidate cell-cell interactions.

#### MI dimensionality selection

To select an appropriate MI dimensionality for each dataset, we estimated the number of overlapping ligand-receptor (LR) co-variation communities in an LR-LR correlation network, using a maximal-clique merging procedure inspired by clique percolation [60]. We classified datasets with more than 100 LR pairs as large (*P>*100) and those with at most 100 LR pairs as small (*P*≤100). Specifically, we pooled LR co-expression neighborhood features across samples, computed pairwise Spearman correlations between LR pairs, and constructed an undirected LR-LR similarity graph by retaining edges above a correlation threshold (*ρ>*0.3 for large datasets and *ρ>*0.2 for small datasets).

Candidate LR components were defined as maximal cliques in this similarity graph and filtered by a minimum clique size (≥4 for large datasets and ≥3 for small datasets). To reduce redundancy among overlapping cliques, we merged clique pairs with substantial overlap (Jaccard index ≥0.5 for large datasets and ≥0.7 for small datasets) and defined each merged component as the union of its constituent LR pairs. The resulting number of non-redundant LR components was used as a data-driven estimate of the MI dimensionality.

Using this procedure, the MI dimensionality was set to 15 for the HGSOC CosMx SMI dataset, 23 for the human melanoma Perturb-FISH dataset, 30 for the mouse aging brain MERFISH dataset, and 11 for the Human Immuno-Oncology FFPE dataset (**Fig**. S30a). Sensitivity analyses comparing *M* =12, 15, and 18 in HGSOC and *M* =20, 23, and 26 in melanoma showed high correspondence of cell-cell MI activity patterns across nearby dimensionality settings (**Fig**. S30b; **Supplementary Notes**).

#### Spatial neighborhood size selection

The spatial neighborhood size (*K*) defines the local scale for constructing ordered neighboring cell pairs. In the HGSOC dataset, cell-cell MI activity patterns were highly concordant across *K*=5, 8, and 10 (**Fig**. S30c; **Supplementary Notes**). In practice, we set *K*=10 for datasets with fewer than 300,000 cells and *K*=5 for datasets with more than 300,000 cells to balance spatial resolution and computational cost.

### Statistics and reproducibility

Statistical tests and significance thresholds are described in the figure legends and Supplementary Notes. Where families of hypotheses were evaluated, multiple-testing correction was applied as specified for the corresponding analysis. In box plots, center lines indicate medians, box limits show the first and third quartiles, and whiskers extend 1.5 times the interquartile range.

### Empirical running time

On a CentOS 7 machine with six CPU cores and one NVIDIA RTX A6000 GPU, SpiderNet required 2 minutes to run on an HGSOC CosMx SMI slice (slice ID: SMI_T10_F001), containing 5,660 cells, 979 genes, and 81 ligand-receptor pairs, representing a typical single-cell spatial transcriptomics analysis. Peak memory usage was 2.05 GB CPU RAM and 688 MiB GPU memory. This run used the NMF-based initialization described above to initialize MI activities and loading matrices.

### SpiderNet-Interactive web tool

To facilitate interactive exploration of tissue communication patterns identified by SpiderNet, we developed SpiderNet-Interactive, a web-based visualization interface that enables users to explore SpiderNet outputs through an integrated workflow (**Fig**. S32). SpiderNet-Interactive organizes key downstream analyses into four workflows: (1) basic communication exploration, which combines *in situ* MI visualization with cell-type-pair and pathway enrichment analyses, (2) MI-informed subtype discovery, (3) multi-hop MI cascade detection, and (4) *in silico* spatial perturbation analysis based on the trained SpiderNet model. By allowing users to select tissue slices, adjust analysis-specific parameters, and move between complementary analyses within the same workspace, the interface supports rapid interrogation of spatial communication programs across slices and experimental conditions. SpiderNet-Interactive runs as a local Flask server and is distributed with the SpiderNet package on GitHub at https://github.com/ma-compbio-lab/SpiderNet.

### Generation and analysis of simulated data

We generated simulated ST datasets with ground truth specified at both the directed cell-pair and molecular-program levels. Each dataset contained 2,000 cells assigned to three cell types in two-dimensional space. Candidate interactions were defined by directed spatial neighbors around a ring-like structure, with each candidate pair assigned to MI-1, MI-2, or a non-interacting state. Gene expression was simulated for 80 genes, including cell-type marker genes and MI-associated ligand-receptor pairs, sender regulators, and receiver targets. For each MI, ligand and sender-regulator expression increased with the cell’s out-going MI activity, whereas receptor and receiver-target expression increased with incoming MI activity. Additive Gaussian noise with mean zero and standard deviations of *σ*=0.4, 0.7, and 1.0 was introduced to gene expression, and sparsity was introduced by randomly setting 10%, 30%, or 50% of expression values to zero. Each condition was evaluated using 30 independently generated replicates.

For simulation benchmarking, method-specific outputs were converted into MI-1- and MI-2-specific directed edge scores, and AUPRC was evaluated against the corresponding ground-truth edge maps. Molecular recovery was assessed by ranking MI-associated ligand-receptor pairs, sender regulators, and receiver targets in the learned loadings. Detailed simulation parameters, output construction, and evaluation procedures are provided in the **Supplementary Notes**.

### Real-tissue benchmarking of inferred communication programs

For real-tissue benchmarking, we evaluated whether inferred communication programs captured biologically meaningful regulatory structure in HGSOC CosMx SMI and aging mouse brain MERFISH datasets. We benchmarked SpiderNet against NMF-LR, COMMOT, scCChain, and Spacia by summarizing each method’s sender × receiver × communication-feature representation into cell-level sender-side and receiver-side communication profiles. Spacia was evaluated in HGSOC but omitted from the aging-brain analysis owing to computational constraints. Methods that require a user-specified latent dimensionality, including SpiderNet, NMF-LR, and scCChain, were run with the same selected dimensionality within each dataset. COMMOT and Spacia, which produce pathway-level or gene-program-based communication outputs, were evaluated using their native communication-feature sets.

We then assessed regulatory coupling to OmniPath-derived upstream-regulator and scSeqComm-derived downstream-target activities, communication-state coherence by comparing communication-derived and expression-derived cell-cell similarity within each cell type, directionality through edge-reversal and opposite-direction controls, and spatial specificity using cell-type-matched distant pseudo-edges. Detailed database construction, cell-level aggregation, and score definitions are provided in the **Supplementary Notes**.

### Spatial transcriptomics datasets and analysis-specific procedures

We analyzed four public spatial transcriptomics datasets spanning cancer, perturbation, and aging settings. Cell-level gene-set scores were calculated as the mean of per-gene z-scored expression values for measured genes in each signature within the relevant cell population.

#### Human ovarian cancer data (CosMx Spatial Molecular Imager)

We downloaded 48 human ovarian cancer tissue samples from 29 individuals, profiled using CosMx Spatial Molecular Imager technology (https://zenodo.org/records/12613839) [21]. These samples measured spatial gene expression of 979 genes in an average of 5,764 cells per sample. We used the seven cell-type annotations provided by the original study: B cells, endothelial cells, fibroblasts, malignant cells, mast cells, monocytes, and T/NK cells. Further details of the ovarian cancer analysis are provided in the **Supplementary Notes**, including tumor functional-state scoring, cancer-associated fibroblast activation scoring, and MI-10-guided *in silico* perturbation.

#### Human melanoma perturbation data (Perturb-FISH)

We analyzed the Perturb-FISH human melanoma dataset (https://www.ncbi.nlm.nih.gov/geo/query/acc.cgi?acc=GSE221321) [34], which profiles how genetic perturbations in melanoma cells influence local immune responses *in situ*. The dataset comprises spatial transcriptomic measurements for 154,418 melanoma and immune cells across 154 genes. Specifically, 35 NF-*κ*B-related genes were individually perturbed in melanoma cells, and spatial expression was measured in the perturbed cells and surrounding immune cells. Previous analyses identified 11 NF-*κ*B-associated perturbation genes that elicit strongly correlated transcriptional responses in neighboring T cells [34]. Further details of the melanoma perturbation analysis are provided in the **Supplementary Notes**, including leave-one-perturbation model training, counterfactual perturbation analysis, benchmarking against spatially aware baselines, and MI-associated GO enrichment.

#### Mouse aging brain data (MERFISH)

We analyzed 10 mouse coronal brain sections spanning ages 3.4 to 33.2 months, profiled using MERFISH (https://zenodo.org/records/13883177) [22]. This dataset quantified the spatial expression of 300 genes in 712,695 cells, with an average of approximately 71,270 cells per sample. We used the 18 cell-type annotations provided by the original study: astrocytes, B cells, endothelial cells, ependymal cells, macrophages, microglia, neural stem cells (NSCs), neuroblasts, excitatory neurons, inhibitory neurons, medium spiny neurons (MSNs), neutrophils, oligodendrocyte precursor cells (OPCs), mature oligodendrocytes, pericytes, T cells, vascular and leptomeningeal cells (VLMCs), and vascular smooth muscle cells (VSMCs). Further details of the aging brain analysis are provided in the **Supplementary Notes**, including cell-type-specific age prediction, cross-context transfer to sagittal MERFISH and hippocampal Stereo-seq, and MI-29-guided *in silico* perturbation.

#### Human Immuno-Oncology FFPE data (MERSCOPE)

We analyzed the MERSCOPE FFPE Human Immuno-Oncology dataset obtained from the Vizgen Data Release Program (see **Data availability**), which profiles large-area human cancer tissue sections spanning colon, liver, melanoma, ovarian, prostate, lung, breast, and uterine cancers. The dataset measures the spatial expression of 500 genes and contains 4,728,128 cells, with 4,707,751 cells retained after quality control. Because cell-type labels were not provided, we performed marker-based cell-type annotation using canonical marker-gene programs curated in CellMarker 2.0 [61]. Briefly, cells were clustered within each cancer type, assigned to major tumor, immune, and stromal compartments by marker-gene-set activity, and further annotated to resolve major non-malignant populations and T cell subtypes. Further details of the Human Immuno-Oncology FFPE analysis are provided in the **Supplementary Notes**, including cell-type annotation and marker validation, projection of the spatially learned MI basis onto TCGA bulk transcriptomes, and survival and immunotherapy-response association analyses.

## Data availability

The spatial transcriptomics data and related datasets used in this study were obtained from publicly available repositories.

- The high-grade serous ovarian cancer CosMx SMI dataset is available from Zenodo at https://zenodo.org/records/12613839.
- The human melanoma Perturb-FISH dataset is available from GEO under accession GSE221321: https://www.ncbi.nlm.nih.gov/geo/query/acc.cgi?acc=GSE221321.
- The mouse aging brain MERFISH data for both coronal and sagittal atlases are available from Zenodo at https://zenodo.org/records/13883177.
- The mouse aging hippocampal Stereo-seq data are available from STOmicsDB under accession STDS0000247.
- The Human Immuno-Oncology FFPE data are available through the Vizgen Data Release Program: https://info.vizgen.com/ffpe-showcase.
- TCGA bulk RNA-seq (TPM) and corresponding clinical annotations were obtained from the NCI Genomic Data Commons (GDC) Data Portal https://portal.gdc.cancer.gov/, for TCGA-BRCA, TCGA-COAD, TCGA-LIHC, TCGA-LUAD, TCGA-LUSC, TCGA-SKCM, TCGA-OV, TCGA-PRAD, and TCGA-UCEC.
- ICB-treated melanoma and lung cancer bulk RNA-seq data with response annotations were obtained from published cohorts [49–51]; transcriptomic data are available from GEO (GSE78220 and GSE135222) and ENA (PRJEB23709).

The data supporting the documented SpiderNet reproduction workflows are publicly available on Zenodo at https://zenodo.org/records/22801896.

## Code availability

The source code of SpiderNet and the notebooks used to reproduce the analyses in this study are available at https://github.com/ma-compbio-lab/SpiderNet. The repository also includes SpiderNet-Interactive for interactive analysis and visual exploration.

## Acknowledgements

This work was supported, in part, by the National Institutes of Health grants UM1HG011593 (J.M.), R03OD039980 (J.M.), UH3CA268202 (J.M.), R01HG007352 (J.M.), R01HG012303 (J.M.), R21DA061481 (J.M.), and U24HG012070 (J.M.). J.M. was additionally supported by the Ray and Stephanie Lane Professorship, a Guggenheim Fellowship from the John Simon Guggenheim Memorial Foundation, and a Google Research Award. S.L. is a Lane Fellow. The funders had no role in study design, data collection and analysis, decision to publish, or preparation of the manuscript.

## Author Contributions

Conceptualization: J.T. and J.M.; Methodology: J.T., S.L., and J.M.; Software – SpiderNet model implementation: J.T.; Software – SpiderNet-Interactive development: J.T. and W.C.; Investigation: J.T., S.L., S.A., Y.Z., and J.M.; Writing – original draft: J.T., S.L., and J.M.; Writing – review and editing: J.T., S.L., S.A., W.C., Y.Z., and J.M.; Funding acquisition: J.M.

