## Supplemental Information for "A meta-interaction basis for cell-cell communication in tissues"

#### A Supplementary Notes

##### A.1 Comparison and harmonization of CCC representations

As summarized in **Table S1**, we compared SpiderNet with COMMOT [1], scCChain [2], Spacia [3], and an NMF-based ligand-receptor baseline (NMF-LR). To place their native outputs in a common representation, we represented each method as a sender × receiver × communication-feature tensor. The communication features retained their native meaning: latent components for SpiderNet, NMF-LR, and scCChain; LR or pathway features for COMMOT; and receiver-gene or pathway features for Spacia. This harmonization enabled common evaluation metrics without assuming semantic equivalence across methods or feature types. For simulation benchmarking, these outputs were further converted into MI-1- and MI-2-specific edge scores as described in **Supplementary Note A.2**; real-tissue benchmarks retained each method’s native features.

Beyond this output harmonization, the methods differ in three aspects relevant to interpretation: the communication features represented, the molecular information used for inference, and the basis of sender-receiver directionality.

**Table S1: Comparison of spatially resolved communication representations included in benchmarking.** Check marks indicate information used explicitly in model inference or representation definition.

| Method | Communication feature program | Information used |  |  | Directional information used |  |
| --- | --- | --- | --- | --- | --- | --- |
|  |  | Spatial proximity | Curated LR pairs | Non-LR cellular expression | LR ordering | Asymmetric sender/receiver programs |
| NMF-LR | LR co-expression latent factors | ✓ | ✓ | × | ✓ | × |
| COMMOT [1] | LR signaling and pathway summaries | ✓ | ✓ | × | ✓ | × |
| scCChain [2] | LR-derived communication programs | ✓ | ✓ | × | ✓ | × |
| Spacia [3] | Sender-receiver influence on receiver genes/pathways | ✓ | × | ✓ | × | × |
| SpiderNet | Meta-interactions linking regulators, LR bridges, and targets | ✓ | ✓ | ✓ | ✓ | ✓ |

*Note:* Spacia defines direction through prespecified sender and receiver cell populations.

The methods differ first in what constitutes a communication feature. COMMOT represents LR-level signaling with pathway-level summaries, whereas NMF-LR and scCChain derive latent communication features from spatial LR patterns. Spacia models sender-to-receiver transcriptomic influence on receiver genes or pathways. In contrast, SpiderNet learns recurrent MIs that jointly organize sender regulators, LR bridges, and receiver targets into directional communication modules across the tissue.

The molecular information used for inference also differs. COMMOT, NMF-LR, and scCChain primarily use spatial proximity and curated LR information, whereas Spacia models spatially resolved sender-to-receiver transcriptomic associations without restricting inference to curated LR pairs. SpiderNet jointly integrates spatial adjacency, curated LR pairs, and cellular expression profiles through reconstruction of LR co-expression and gene expression.

Finally, the basis of directionality differs. COMMOT, NMF-LR, and scCChain derive direction primarily from ligand-to-receptor ordering, whereas Spacia defines direction through prespecified sender and receiver cell populations. In SpiderNet, directionality is informed jointly by LR ordering and asymmetric sender- and receiver-side transcriptional programs.

#### A.2 Simulation design and recovery evaluation

**Simulation data generation.** We simulated 2,000 cells uniformly distributed in a two-dimensional space and assigned them to three cell types (CT-A, CT-B, or CT-C). For cells near the boundary of a central circular region, directed spatial edges were drawn to their ten nearest neighbors. Each edge was randomly assigned to MI-1, MI-2, or a non-interacting state. Interacting edges had ground-truth MI activity 1, whereas non-interacting edges and all other cell pairs had no MI activity.

To model both cell-intrinsic identity and MI-dependent regulatory effects, we simulated the expression of 80 genes. Each cell type was assigned five marker genes randomly sampled from the full gene set, with elevated expression in the corresponding cell type. For each MI, we specified ten sender regulators, ten receiver targets, and ten ligand-receptor pairs. Ligand and regulator expression scaled with each cell’s total sending activity for a given MI, whereas receptor and target expression scaled with its total receiving activity for the same MI.

Expression values were sampled from truncated normal distributions on $[0, \infty)$. Cell-type markers had mean $\mu=3$ in the corresponding cell type and $\mu=0.6$ otherwise. MI-associated genes had means proportional to sending or receiving MI activity with baseline scale $\mu_0=3$. We introduced expression noise by varying the standard deviation over $\sigma=0.4, 0.7$, and $1.0$, and introduced sparsity by randomly setting $\rho=10\%$, $30\%$, or $50\%$ of expression values to zero. Each setting was evaluated using 30 independently generated simulation replicates.

**Construction of MI-specific edge scores.** For each candidate directed edge, we derived separate scores for MI-1 and MI-2. SpiderNet, NMF-LR, and scCChain were fitted with two latent communication components, which were matched one-to-one to the two ground-truth MIs using Spearman correlation. COMMOT and Spacia outputs were instead aggregated over the ground-truth LR-pair and gene signatures, respectively, that defined each MI. This yielded MI-1- and MI-2-specific edge-score maps for every method.

**Simulation evaluation metrics.** Edge recovery performance was evaluated for each ground-truth MI by thresholding the corresponding inferred communication scores across their full range to generate precision-recall curves for recovering ground-truth interacting edges. Performance was summarized by the area under the precision-recall curve (AUPRC) across replicates for each noise and dropout setting. AUPRC was used because true MI edges were sparse relative to the candidate directed pairs.

Molecular recovery performance for SpiderNet was evaluated by ranking ligand-receptor pairs, sender regulators, and receiver targets according to their loadings in the learned MI corresponding to each ground-truth MI. For each feature class, features were ranked in ascending order of loading, such that larger ranks corresponded to larger loadings, and the mean rank of ground-truth MI-associated features was compared with that of non-associated features. Greater rank separation in favor of MI-associated features indicated better recovery of the defining molecular program.

##### A.3 Real-tissue CCC benchmarking

For all compared methods, outputs were first represented as sender × receiver × communication-feature tensors as described above. For the regulatory-coupling and cell-cell similarity benchmarks, these tensors were aggregated into cell-level sender and receiver profiles: summing over receiver cells yielded each cell’s sender profile, whereas summing over sender cells yielded its receiver profile. The communication-feature dimensions were those produced by each method: 15 for SpiderNet, NMF-LR, and scCChain, 52 for COMMOT, and 14 for Spacia in the HGSOC CosMx dataset; and 30 for SpiderNet, NMF-LR, and scCChain, and 6 for COMMOT in the aging brain MERFISH dataset. The spatial-specificity control used edge-level MI activities directly.

***Sender- and receiver-side regulatory coupling evaluation.*** To test whether inferred communication signals were coupled to expected regulatory programs, we evaluated their association with curated upstream regulators in sender cells and curated downstream targets in receiver cells. For the upstream and downstream benchmarks, we derived ligand-receptor-specific upstream regulator sets from OmniPath [4] and downstream target sets from scSeqComm [5], respectively, both based on curated transcriptional regulatory interactions. All gene sets were restricted to genes measured in the corresponding dataset.

For each curated ligand-receptor pair, upstream-regulator and downstream-target activities were computed as the mean normalized expression of their annotated genes. The best-matching-feature procedure was applied identically to all methods. Sender-side coupling was the Spearman correlation between regulator activity and each dimension of the sender-side communication profile; receiver-side coupling was defined analogously using target activity and the receiver-side communication profile. For each method and ligand-receptor pair, the final coupling score was the maximum Spearman correlation across communication-feature dimensions. Higher values indicate closer alignment between inferred communication signals and curated regulatory programs.

***MI directionality evaluation.*** We evaluated MI directionality using two complementary analyses in the HGSOC CosMx and aging mouse brain MERFISH datasets. First, we tested whether the same neighboring-cell pair showed similar MI activities in both sender-receiver directions. Within each tissue slice, we retained cell pairs represented by edges in both directions and calculated, for each MI, the Pearson correlation between their activities in the two directions. These correlations were summarized across slices.

Second, we tested whether the inferred edge direction was required for regulatory concordance. We constructed an edge-reversed control that retained each edge’s MI activity but exchanged its sender and receiver cells, and repeated the sender- and receiver-side regulatory coupling analyses described above. For each ligand-receptor pair, both conditions used the MI selected under the original edge direction. Coupling values were summarized across slices by their mean and 95% confidence interval.

***Cell-cell similarity evaluation.*** Motivated by the view that transcriptional cell states are embedded in multicellular spatial niches [6], we asked whether inferred communication states retained transcriptionally coherent variation within annotated cell types. We defined a cell-cell similarity concordance score to test whether transcriptionally similar cells also exhibited similar inferred communication states.

For each method, we constructed a communication-state representation for each cell by concatenating its sender-side and receiver-side communication profiles. Within each cell type, Pearson correlations between these representations defined the communication-derived cell-cell similarity matrix, whereas correlations between normalized expression profiles defined the expression-derived similarity matrix.

Concordance was quantified as the Spearman correlation between the vectorized upper triangles of the two matrices. Higher values indicate that the inferred communication representation better preserves transcriptionally coherent relationships among cells.

Across the HGSOC CosMx and aging mouse brain MERFISH datasets, SpiderNet generally achieved higher similarity concordance than the other methods across evaluated cell types (**Fig. S6**), indicating that its inferred MIs preserve coherent within-cell-type variation beyond broad cell identity.

***Spatial-specificity control.*** To assess whether inferred MIs reflected local spatial organization, we performed a cell-type-matched counterfactual spatial-specificity analysis in the HGSOC CosMx and aging mouse brain MERFISH datasets. For each MI, we selected the sender-receiver cell-type context with the highest mean MI activity across observed neighboring edges. Within each tissue slice, every evaluated local edge was matched one-to-one to a distant pseudo-edge by retaining the same sender and sampling a receiver of the same cell type outside the sender’s 1,000 nearest spatial neighbors. With the trained model held fixed, MI activities were recomputed on the pseudo-edges.

Spatial enrichment was summarized as the $\log_2$ fold change in mean MI activity between local and distant edges, with positive values indicating stronger activity among local neighbors. Across the dominant MI contexts, 13/15 HGSOC MIs and 21/30 aging-brain MIs showed positive local-versus-distant enrichment, with median $\log_2$ fold changes of 0.05 in both datasets (**Fig. S7**), supporting modest local enrichment across most evaluated MI contexts.

###### **A.4 Additional details of HGSOC analyses**

For the HGSOC analyses, the malignant-cell hypoxia score used positively correlated ovarian cancer markers from CancerSEA [7]. HIF-1 signaling, PD-1/PD-L1 checkpoint, and ECM-receptor interaction scores used the corresponding KEGG gene sets [8]. The malignant TIL score used the upregulated malignant TIL program defined in the original HGSOC study [9]. Fibroblast CAF scores used established CAF markers [10, 11], and CD8<sup>+</sup> T-cell exhaustion scores used canonical exhaustion and checkpoint markers [12].

***CAF-malignant program coupling evaluation.*** We evaluated whether alternative CCC representations captured the fibroblast→malignant C5 program represented by SpiderNet MI-10. Outputs from NMF-LR, COMMOT, scCChain, and Spacia were mapped to the common communication activity tensor (sender × receiver × communication-feature) and evaluated on the same fibroblast → malignant C5 interactions. Edge-level activities were max-aggregated to sender fibroblasts or receiver malignant C5 cells. To enable comparison across method-specific communication features, we used MI-10 for SpiderNet and selected the fibroblast→malignant C5 feature most strongly coupled to the relevant programs for each baseline. For each baseline, we evaluated every fibroblast→malignant C5 communication feature by splitting cells at its median activity and calculating SMDs for CAF, PD-1/PD-L1 checkpoint, ECM-receptor interaction, and hypoxia programs. The feature with the best average rank across these four SMDs was retained, and coupling was reported as the corresponding SMDs between high- and low-communication cell groups.

***In silico perturbation of top MI-10 LR pairs.*** To assess the model-predicted perturbability of the MI-10 molecular bridge, we performed MI-10-guided *in silico* LR perturbations on fibroblast→malignant C5 edges with the trained SpiderNet model held fixed. Top MI-10 LR pairs were defined as those with normalized LR loadings > 0.5. For each selected pair, ligand expression in fibroblast senders and receptor expression in malignant C5 receivers were set to zero. **KO-TopMI10** knocked out the selected top LR pairs, whereas **KO-Random** knocked out a size-matched random set of non-top LR pairs as a control. Perturbation effects were quantified as post-minus-pre changes in predicted fibroblast CAF and malignant immune-suppressive program scores.

#### A.5 Additional details of melanoma Perturb-FISH analyses

For the melanoma Perturb-FISH analyses, MI-17 melanoma-sender and T-cell-receiver program scores used GO gene sets enriched among the corresponding regulator and target genes, respectively [13].

**Leave-one-perturbation model training.** We benchmarked SpiderNet, Celcomen [14], and a niche-summary linear model on predicting gene-wise T-cell responses to 11 melanoma NF-κB perturbations previously shown to induce correlated responses in neighboring T cells [15]. For each perturbed gene $g$, melanoma cells carrying that perturbation were excluded from training, whereas immune cells, unperturbed melanoma cells, and melanoma cells carrying other perturbations were retained. Each model was fitted to the filtered tissue before counterfactual evaluation using the held-out perturbed melanoma-cell profiles as described below.

**Counterfactual perturbation analysis and T-cell response evaluation.** For each perturbed gene $g$, we constructed a counterfactual slice by replacing the expression profile of each unperturbed melanoma cell with that of a randomly sampled $g$-perturbed melanoma cell, while retaining all other cells and spatial coordinates. To evaluate newly introduced exposure to perturbation $g$, we defined a fixed reference set of T cells that were not adjacent to $g$-perturbed melanoma cells in the original slice. The **predicted T-cell response** was the gene-wise log fold change in mean predicted expression for these same T cells between the counterfactual and original slices. The **observed T-cell response**, used as the reference response, was the corresponding gene-wise log fold change in observed expression between T cells adjacent to $g$-perturbed melanoma cells and the non-adjacent reference T cells in the original slice. Prediction accuracy for each perturbed gene was quantified as the Spearman correlation between predicted and observed response vectors across measured genes.

We compared SpiderNet with two baselines compatible with the same leave-one-perturbation and counterfactual response-prediction framework: Celcomen [14] and a niche-summary linear regression model (LinearModel). Celcomen is a causality-inspired graph neural network that models intercellular regulatory effects to generate counterfactual post-perturbation expression profiles. LinearModel predicted center-cell expression from cell type and local-niche features, including cell-type-pair composition, mean neighboring-cell expression, and mean sending and receiving LR expression.

**Perturbation-responsive program coupling evaluation.** For SpiderNet, NMF-LR, and scCChain, we identified the melanoma-to-T-cell communication feature with the largest mean activity increase across the 11 perturbations using the same counterfactual replacement framework. Mean outgoing activity across melanoma-to-T-cell edges defined sender-cell (melanoma) communication activity, whereas mean incoming activity defined receiver-cell (T-cell) communication activity. Melanoma cells and T cells were stratified by their respective median communication activities, and coupling was quantified as the SMD in the corresponding sender or receiver program scores between high- and low-communication groups.

**In silico perturbation of top MI-17 LR pairs.** To assess the model-predicted perturbability of the MI-17 molecular bridge, we performed MI-17-guided *in silico* LR perturbations on melanoma→T-cell edges with the trained SpiderNet model held fixed. Top MI-17 LR pairs were defined as those with normalized

LR loadings $>0.5$. For each selected pair, ligand expression was perturbed in melanoma senders and receptor expression in neighboring T-cell receivers. **KO-TopMI17** set these genes to zero, whereas **50%-TopMI17** reduced their expression by 50%. A size-matched random set of non-top LR pairs was knocked out as a control. Perturbation effects were quantified as post-minus-pre changes in predicted MI-17-associated melanoma-sender and T-cell-receiver program scores.

#### A.6 Additional details of aging mouse brain analyses

For the aging mouse brain analyses, aging scores were computed from aging-associated markers reported in the original MERFISH atlas [16]. Genes with a mean-expression log fold change $>0.2$ in old versus young samples were retained.

**Cell-type-specific cellular age prediction.** To evaluate whether each representation captured aging-relevant cellular variation, we trained cell-type-specific linear models to predict tissue-section age from each cell’s representation. All methods were evaluated using the same age-stratified 10-fold cross-validation framework. Within each cell type, cells were divided into folds such that every test fold contained cells from each age group. For SpiderNet, NMF-LR, COMMOT, and scCChain, method-specific sending and receiving activities were aggregated over spatial neighbors to obtain cell-level features, whereas Banksy used weighted neighborhood-aggregated expression features. Out-of-fold predictions were pooled to calculate the Pearson correlation between predicted and observed section age. Cell types with fewer than ten cells were excluded.

**Cross-context transfer to sagittal MERFISH and hippocampus Stereo-seq.** To assess cross-context transfer, we applied models trained on the coronal aging brain MERFISH dataset to sagittal sections from the same atlas [16] and to an independent aging hippocampus Stereo-seq dataset [17], using the shared 300-gene MERFISH panel. For SpiderNet, target-dataset MIs were inferred with the pretrained encoder before applying the pretrained cell-type-specific age models. For baseline methods, target-dataset representations were constructed in the corresponding coronal-trained feature space and directly supplied to the pretrained cell-type-specific age models. We evaluated whether age-related signals were preserved by comparing predicted ages between young and old samples within each cell type. Samples older than 19 months were defined as old in sagittal MERFISH, whereas the original labels were used for hippocampal Stereo-seq.

**T-cell-associated aging-program coupling evaluation.** We evaluated whether alternative CCC representations captured the T-cell-associated aging program represented by SpiderNet MI-29. Outputs from NMF-LR, COMMOT, and scCChain were mapped to the common communication activity tensor (sender × receiver × communication-feature) and evaluated on the same T-cell → neighbor interactions. Edge-level activities were max-aggregated to sender T cells. To enable comparison across method-specific communication features, we used MI-29 for SpiderNet and selected the T-cell-sending feature most strongly coupled to the aging program for each baseline. For each baseline, T cells were split by the median strength of each sending feature, and the feature with the largest SMD in T-cell aging score was selected. Coupling was reported as the SMD in aging module scores between the resulting high- and low-communication T-cell groups.

**MI-29-guided in silico gene perturbation and cell replacement.** To examine model-predicted responses to perturbation of the T-cell-derived MI-29 program, we performed MI-29-guided *in silico* gene perturbation and cell replacement with the trained SpiderNet model held fixed. Top MI-29 LR pairs, sender regulators, and receiver targets had normalized loadings $>0.5$, whereas non-top controls had loadings in $[0.25, 0.5]$. Ligands and regulators were perturbed in T-cell senders, and receptors and targets in neighboring receivers. **KO-TopMI29** set top-feature genes to zero, **50%-TopMI29** reduced their expression by 50%, and **KO-NonTopMI29** and **KO-Random** knocked out non-top or size-matched random features, respectively. For cell replacement, a cell’s expression profile was replaced with that of a sampled T cell, astrocyte, or endothelial cell while retaining its coordinates. Perturbation effects were quantified as post-minus-pre changes in predicted neighbor age using the pretrained age models. Cell-replacement effects additionally included changes in neighbor-received MI-29 activity.

#### A.7 Additional details of pan-cancer FFPE atlas analyses

For the pan-cancer FFPE atlas analyses, tumor-cell invasion, angiogenesis, and hypoxia scores used the corresponding CancerSEA gene sets [7].

**Cell-type annotation and marker validation.** Because the MERSCOPE FFPE atlas lacked cell-type labels, we annotated cells using unsupervised clustering and canonical marker-gene programs. Given the targeted 500-gene panel, annotations were interpreted primarily at the major-compartment level for pan-cancer MI analyses. For each cancer type, we first applied library-size normalization followed by log transformation, computed a PCA embedding using the top 50 principal components, and performed Louvain clustering. Major cell types were assigned using canonical marker genes curated in CellMarker 2.0 [18]. Cancer-type-specific tumor populations were annotated after excluding clusters enriched for immune, endothelial, fibroblast, or other stromal markers. To resolve T cell subtypes, we subset T cells, reclustered them, and annotated them as $CD4^+$ T, $CD8^+$ T/NK, or Treg cells using established subtype markers [18]. The final atlas contained eight cancer-specific tumor populations and 11 non-malignant populations: B cells, $CD4^+$ T cells, $CD8^+$ T/NK cells, dendritic cells, endothelial cells, epithelial cells, fibroblasts, macrophages, mast cells, Treg cells, and a low-expression group. Marker-set activity broadly supported the major-compartment assignments (Fig. S22).

**Sub-slice pan-cancer analysis.** Because each large-area tissue section contained hundreds of thousands of cells and spanned a broad spatial field, we partitioned each section into 20 non-overlapping spatial sub-slices to support scalable SpiderNet training. All sub-slices from eight cancer types were jointly modeled, with each treated as a separate tissue patch for model fitting.

**Fibroblast-tumor malignant-program coupling evaluation.** We evaluated whether alternative CCC representations captured the fibroblast→tumor malignant-program coupling represented by SpiderNet MI-4. Outputs from COMMOT and scCChain were mapped to the common communication activity tensor (sender × receiver × communication-feature) and evaluated on the same fibroblast→tumor interactions within each cancer type. Edge-level activities were max-aggregated to receiver tumor cells to obtain each cell’s incoming fibroblast→tumor communication activity. To enable comparison across method-specific communication features, we used MI-4 for SpiderNet and selected the fibroblast→tumor feature most strongly coupled to the three malignant programs for each baseline. For each baseline, tumor cells were split by the median activity of each receiving feature, and the feature with the strongest mean coupling to invasion, angiogenesis, and hypoxia across cancer types was selected. Coupling was reported as SMDs in the three malignant-program scores between the resulting high- and low-communication tumor-cell groups.

***In silico* perturbation of top MI-4 LR pairs.** To assess the model-predicted perturbability of the MI-4 molecular bridge, we performed MI-4-guided *in silico* LR perturbations on fibroblast→tumor edges with the trained SpiderNet model held fixed. Top MI-4 LR pairs were defined as those with normalized LR loadings $>0.5$. For each selected pair, ligand expression was perturbed in fibroblast senders and receptor expression in neighboring tumor receivers. **KO-TopMI4** set these genes to zero, whereas **50%-TopMI4** reduced their expression by 50%. A size-matched random set of non-top LR pairs was knocked out as a control. Perturbation effects were quantified as post-minus-pre changes in predicted invasion, angiogenesis, and hypoxia scores in receiver tumor cells.

#### A.8 Bulk projection and clinical association of the pan-cancer MI basis

**Bulk MI projection.** To extend the spatially learned MI programs to clinical cohorts, we projected the pan-cancer MI basis learned from the Human Immuno-Oncology FFPE spatial transcriptomics data onto tumor bulk RNA-seq profiles. Because bulk transcriptomes lack spatial cell-pair resolution, this projection estimates sample-level MI abundance rather than cell-pair-level communication activity.

For each bulk sample $j$, we inferred $\tilde{\mathbf{I}}_{j,\cdot} \in [0, 1]^M$, representing sample-level abundances of the $M$ MI programs. Bulk projection fixed the non-negative intrinsic, regulator, target, and LR loading matrices learned by the pan-cancer spatial model. The intrinsic loading matrix $\widehat{\mathbf{W}}^{(\text{intrinsic})} \in \mathbb{R}^{C \times G}$ captures cell-intrinsic programs, whereas the regulator, target, and LR loading matrices, $\widehat{\mathbf{W}}^{(\text{regulator})} \in \mathbb{R}^{M \times G}$, $\widehat{\mathbf{W}}^{(\text{target})} \in \mathbb{R}^{M \times G}$, and $\widehat{\mathbf{W}}^{(\text{LR})} \in \mathbb{R}^{M \times P}$, anchor each MI to sender regulators, receiver targets, and LR bridges, respectively (Eqs. 7 and 8). Sample-level MI abundances were inferred by jointly reconstructing bulk gene expression and a bulk-derived LR co-expression proxy. The derivation below translates the cell-level expression model to the bulk-sample level, introduces the LR co-expression proxy as a complementary molecular readout, and combines both components in a joint constrained optimization across cohorts.

First, we derived a sample-level analogue of the SpiderNet expression model by averaging the cell-level decomposition over the cells represented by bulk sample $j$:

$$\begin{aligned} E_{j,g}^{\text{bulk}} &\approx \frac{1}{N_j} \sum_{i=1}^{N_j} \left[ \sum_{c=1}^C H_{i,c} \widehat{W}_{c,g}^{(\text{intrinsic})} + \sum_{m=1}^M \bar{I}_{i,m}^S \widehat{W}_{m,g}^{(\text{regulator})} + \sum_{m=1}^M \bar{I}_{i,m}^R \widehat{W}_{m,g}^{(\text{target})} \right] \\ &= \sum_{c=1}^C \tilde{H}_{j,c} \widehat{W}_{c,g}^{(\text{intrinsic})} + \sum_{m=1}^M \tilde{I}_{j,m}^S \widehat{W}_{m,g}^{(\text{regulator})} + \sum_{m=1}^M \tilde{I}_{j,m}^R \widehat{W}_{m,g}^{(\text{target})}, \end{aligned} \quad (\text{S1})$$

where $\tilde{H}_{j,c} = N_j^{-1} \sum_i H_{i,c}$ is the sample-level abundance of intrinsic program $c$, whereas $\tilde{I}_{j,m}^S = N_j^{-1} \sum_i \bar{I}_{i,m}^S$ and $\tilde{I}_{j,m}^R = N_j^{-1} \sum_i \bar{I}_{i,m}^R$ are the sample-level sending and receiving abundances of MI $m$. Here, $\bar{I}_{i,m}^S$ and $\bar{I}_{i,m}^R$ are the cell-level aggregated MI profiles defined in Eq. 5. Thus, bulk expression is decomposed into sample-level intrinsic-program abundance and MI-associated sending and receiving components.

At the sample level, each directed interaction contributes once to the sending side and once to the receiving side of the same MI. Thus, for bulk projection, we approximate each MI's sample-level sending and receiving abundances with a shared coefficient $\tilde{I}_{j,m}$, yielding

$$E_{j,g}^{\text{bulk}} \approx \sum_{c=1}^C \tilde{H}_{j,c} \widehat{W}_{c,g}^{(\text{intrinsic})} + \sum_{m=1}^M \tilde{I}_{j,m} \widehat{W}_{m,g}^{(\text{MI})}, \quad \widehat{\mathbf{W}}^{(\text{MI})} = \widehat{\mathbf{W}}^{(\text{regulator})} + \widehat{\mathbf{W}}^{(\text{target})}, \quad (\text{S2})$$

where the combined loading $\widehat{W}^{(\text{MI})}$ summarizes the sender- and receiver-side transcriptional signatures of each MI for bulk reconstruction.

Second, we constructed an LR co-expression proxy from bulk RNA-seq. For each of the $P$ LR pairs used in SpiderNet training, the bulk LR proxy $\mathbf{Z}^{\text{bulk}} \in \mathbb{R}^{J \times P}$ was defined as the geometric mean of measured ligand- and receptor-side expression (Eq. 1). Although this is not a direct measurement of spatial LR co-expression, it provides an expression-derived proxy for the LR pairs. Using the same sample-level MI abundances, we approximated

$$\mathbf{Z}_{j,p}^{\text{bulk}} \approx \sum_{m=1}^M \tilde{I}_{j,m} \widehat{W}_{m,p}^{(\text{LR})}. \quad (\text{S3})$$

The same sample-level MI abundances were used to reconstruct both bulk readouts, so that the inferred MI profile was supported jointly by gene-expression and LR-derived evidence.

Let $q = 1, \dots, Q$ index bulk cohorts, with $\mathbf{E}^{\text{bulk},(q)} \in \mathbb{R}^{J_q \times G}$ and $\mathbf{Z}^{\text{bulk},(q)} \in \mathbb{R}^{J_q \times P}$ denoting the gene expression and LR proxy for cohort $q$. Across cohorts, we kept the spatially learned loading matrices fixed and jointly estimated MI abundances $\tilde{\mathbf{I}}^{(q)} \in \mathbb{R}^{J_q \times M}$, intrinsic program abundance $\tilde{\mathbf{H}}^{(q)} \in \mathbb{R}^{J_q \times C}$, cohort-specific gene-expression offset $\boldsymbol{\alpha}^{(q)} \in \mathbb{R}^{1 \times G}$, and LR-proxy offsets $\boldsymbol{\delta}^{(q)} \in \mathbb{R}^{1 \times P}$ by minimizing:

$$\begin{aligned} \min_{\{\tilde{\mathbf{H}}^{(q)}, \tilde{\mathbf{I}}^{(q)}, \boldsymbol{\alpha}^{(q)}, \boldsymbol{\delta}^{(q)}\}_{q=1}^Q} \quad & \sum_{q=1}^Q \frac{1}{J_q} \left[ \underbrace{\frac{1}{G} \left\| \left( \mathbf{E}^{\text{bulk},(q)} - \tilde{\mathbf{H}}^{(q)} \widehat{\mathbf{W}}^{(\text{intrinsic})} - \tilde{\mathbf{I}}^{(q)} \widehat{\mathbf{W}}^{(\text{MI})} - \mathbf{1}_{J_q \times 1} \boldsymbol{\alpha}^{(q)} \right) \boldsymbol{\Omega} \right\|_F^2}_{\text{bulk gene-expression reconstruction}} \right. \\ & \left. + \underbrace{\frac{1}{2P} \left\| \left( \mathbf{Z}^{\text{bulk},(q)} - \tilde{\mathbf{I}}^{(q)} \widehat{\mathbf{W}}^{(\text{LR})} - \mathbf{1}_{J_q \times 1} \boldsymbol{\delta}^{(q)} \right) \boldsymbol{\Psi} \right\|_F^2}_{\text{bulk-derived LR-proxy reconstruction}} \right] \quad (\text{S4}) \end{aligned}$$

$$\begin{aligned} \text{s.t.} \quad & \tilde{\mathbf{H}}^{(q)} \mathbf{1}_{C \times 1} = \mathbf{1}_{J_q \times 1}, \quad q = 1, \dots, Q, \\ & \tilde{H}_{j,c}^{(q)} \geq 0, \quad \forall j, c, \quad q = 1, \dots, Q, \\ & 0 \leq \tilde{I}_{j,m}^{(q)} \leq 1, \quad \forall j, m, \quad q = 1, \dots, Q. \end{aligned}$$

Here, the cohort-specific offsets account for systematic differences between bulk data and the ST-derived signatures. The diagonal weight matrices $\boldsymbol{\Omega} = \text{diag}(w_1, \dots, w_G)$ and $\boldsymbol{\Psi} = \text{diag}(v_1, \dots, v_P)$ assign higher weights to genes and LR pairs whose loadings are concentrated in a small subset of MIs, and lower weights to features shared broadly across MIs. Weights were quantified by the Kullback-Leibler divergence between each feature's normalized loading profile and a uniform distribution across MIs. The constrained optimization problem was solved by alternating optimization, with $\tilde{\mathbf{H}}$ and $\tilde{\mathbf{I}}$ updated using OSQP [19] and cohort-specific offsets updated in turn until convergence.

**Pseudo-bulk validation.** To validate the bulk projection, we generated pseudo-bulk samples from the Human Immuno-Oncology FFPE spatial dataset by aggregating gene expression within non-overlapping sub-slices. We first validated the LR co-expression proxy used in the MI projection objective. The bulk-derived LR proxy showed high Spearman correlations across sub-slices with spatial LR co-expression averaged within the corresponding sub-slices, evaluated separately for each LR pair (Fig. S25a). We then projected each pseudo-bulk sample and, across all MI-sub-slice combinations, compared its estimated MI abundance with observed sub-slice-average MI abundance derived from cell-pair-level MI activities. Projected MI abundance increased monotonically across tertiles of observed abundance (Fig. S25b), indicating that the projection preserved relative MI abundance ordering despite the loss of spatial cell-pair information.

**Clinical associations in TCGA cohorts.** To assess associations between projected MI abundance and patient overall survival, we analyzed nine TCGA cohorts. Within each TCGA cohort, GDC-defined tumor samples were stratified into low-, intermediate-, and high-MI-abundance groups for Kaplan-Meier visualization. For statistical inference, we fit cohort-wise Cox proportional hazards models using standardized continuous MI abundance, with age, sex, and tumor stage included as clinical covariates. Hazard ratios and Wald $P$ values were reported, with $HR > 1$ indicating increased hazard per standard-deviation increase in MI abundance and an association with poorer survival.

Because fibroblast→tumor MI-4 was associated with malignant invasion, angiogenesis, and hypoxia programs, we tested whether it provided prognostic information beyond these canonical tumor programs. Within each TCGA cohort, we compared nested Cox models containing clinical covariates and bulk invasion, angiogenesis, and hypoxia scores, with or without projected MI-4 abundance. This analysis quantified the incremental prognostic value of MI-4 after accounting for the canonical malignant programs with which it was associated. Cohorts with an MI-4 $HR > 1$ and likelihood-ratio-test $P < 0.05$ were considered to show incremental adverse prognostic information beyond that captured by the clinical covariates and MI-4-linked tumor programs (Fig. S26b).

**Clinical associations in immunotherapy cohorts.** To test whether projected MI-4 abundance was associated with immune-checkpoint blockade (ICB) response, we projected bulk RNA-seq profiles from published ICB-treated melanoma and lung cancer cohorts onto the pan-cancer MI basis using the joint gene-expression and LR decomposition (Supplementary Note A.8). Within each cohort-treatment setting, harmonized responder and non-responder groups were compared using Wilcoxon rank-sum tests. Associations were summarized as signed $-\log_{10} P$ values, with positive and negative values indicating responder and non-responder enrichment, respectively. Within each cohort-treatment setting, candidate features were ranked by non-response association, and the resulting ranks were averaged across the four settings to obtain an overall mean within-setting rank (Fig. S26c).

#### A.9 Model component ablations

To assess the contribution of each model component, we compared the full SpiderNet model with gene-expression-reconstruction-only, LR-co-expression-reconstruction-only, and no-intrinsic-component variants. These variants probe the contributions of intracellular expression modeling, LR-mediated intercellular anchoring, and separation of communication-associated expression from broad cell-type baseline variation, respectively. We evaluated them on two complementary tasks: regulatory coupling and held-out perturbation-effect prediction.

**Regulatory coupling under component ablation.** Using the sender- and receiver-side regulatory-coupling benchmark (Supplementary Note A.3), all variants were evaluated in the HGSOC CosMx and aging brain MERFISH datasets. The full model maintained strong coupling across both datasets and communication directions (Fig. S29a,b). The gene-expression-only variant remained strong in several settings but showed substantially weaker receiver-side coupling in the aging brain, whereas the LR-co-expression-only variant generally showed weaker sender-side coupling. Removing the intrinsic component also reduced coupling on both sides, consistent with its intended role in separating MI-associated regulatory variation from baseline cellular identity.

**Held-out perturbation prediction under component ablation.** Using the leave-one-perturbation and counterfactual response-evaluation workflow (Supplementary Note A.5), we evaluated all variants on T-cell perturbation-effect prediction in melanoma Perturb-FISH. Prediction accuracy was quantified by the Spearman correlation between observed and predicted T-cell responses across measured genes. The full and no-intrinsic models showed broadly comparable performance. The single-objective variants were generally weaker or more variable, although both retained predictive signal for some perturbations (Fig. S29c).

Together, these results show that joint gene-expression and LR reconstruction support regulatory organization and perturbation prediction, whereas the intrinsic component contributes more clearly to regulatory coupling than to held-out predictive accuracy.

#### A.10 MI-basis stability and robustness

**Stability across MI dimensionality.** To assess whether the inferred MI structure was stable across nearby MI dimensionalities rather than tied to a specific dimension, we compared cell-cell MI activities across $M=12, 15$, and $18$ in the HGSOC CosMx dataset and across $M=20, 23$, and $26$ in the melanoma Perturb-FISH dataset. Corresponding MI programs showed high Spearman correlations of cell-cell MI activities across adjacent dimensionality settings (Fig. S30b), indicating that the dominant MI activity patterns were stable across the tested values of MI dimensionality.

**Robustness to spatial neighborhood size.** Because the number of nearest neighbors ($K$) determines the spatial scale over which ordered cell-cell pairs are constructed, we assessed the sensitivity of the inferred MI patterns to this choice. We retrained SpiderNet on the HGSOC CosMx dataset using $K=5, 8$, and $10$ nearest neighbors and compared cell-cell MI activities across models. MI activities were highly concordant between adjacent settings (Fig. S30c), indicating limited sensitivity of the dominant MI activity patterns to neighborhood size.

**Robustness to cell-type label misannotation.** Because spatial transcriptomics annotations may contain errors or ambiguous cell-type assignments, we assessed the sensitivity of inferred MIs to cell-type-label misannotation. In the HGSOC CosMx dataset, we randomly selected 10%, 20%, or 30% of cells, re-assigned their annotations to an incorrect cell type, and retrained SpiderNet under each misannotation rate. MI activities remained highly concordant with those from the original-label model even at 30% mis-annotation (Fig. S31a). This high concordance under increasing misannotation rates indicated limited sensitivity to moderate cell-type-label misannotation.

**Stability across random initialization.** To assess reproducibility across stochastic initialization, we retrained SpiderNet on the HGSOC CosMx dataset using ten different random seeds and aligned MIs one-to-one based on Spearman correlations of cell-cell MI activities. Edge-level MI activities, LR loadings, sender-regulator loadings, and receiver-target loadings showed high Pearson correlations with the reference model (Fig. S31b), indicating stable MI activities and molecular signatures across initializations.

**Stability with reduced training data.** To assess whether the learned MI basis remained stable with fewer training samples, we retrained SpiderNet on ten random subsets, each containing 70% of the HGSOC slices. Each subsampled fit was aligned with the full-data reference based on cell-cell MI activity correlations. The subsampled models retained high concordance in edge-level MI activities and molecular loadings, particularly MI activities and LR loadings (Fig. S31c), supporting the stability of the learned

MI basis under a moderate reduction in training sample size.

#### B Supplementary Figures

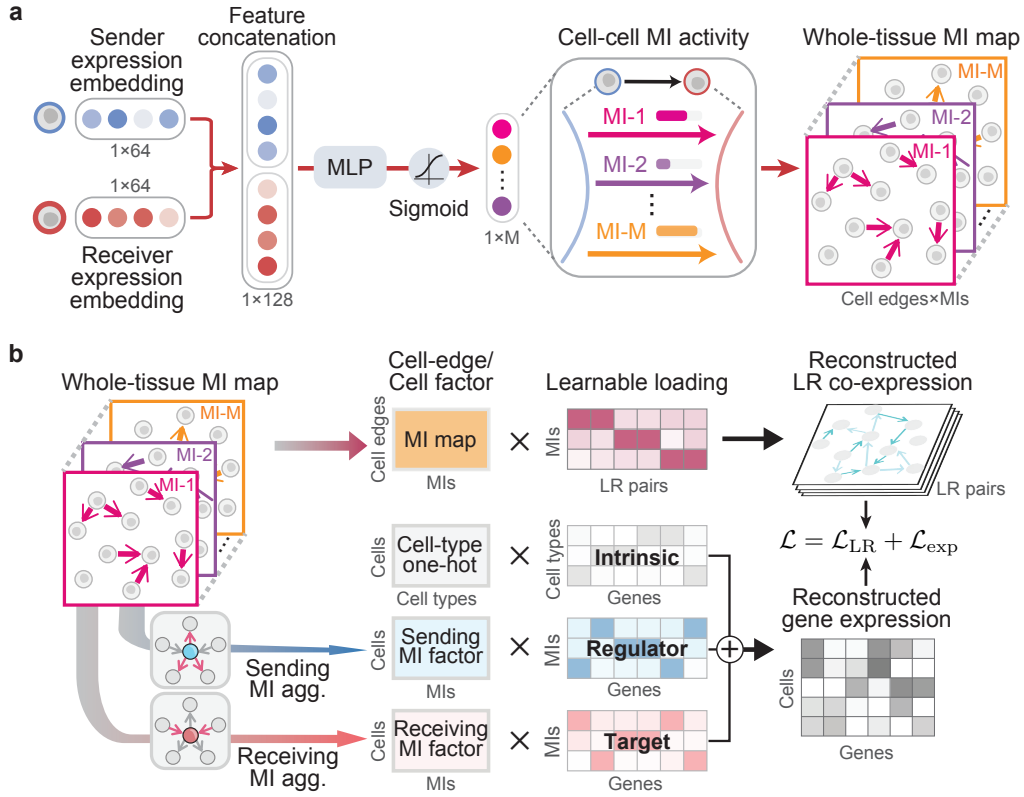

**Figure S1: Implementation details of SpiderNet.** **a**, For each ordered sender-receiver cell pair, SpiderNet concatenates sender and receiver expression embeddings and uses an MLP to infer directional MI activities across multiple MI dimensions, producing a tissue-wide cell-cell MI map. **b**, MI activities are molecularly anchored by reconstructing LR co-expression through MI-specific LR loadings and gene expression through intrinsic and role-specific loadings after MI aggregation into sending and receiving cell factors.

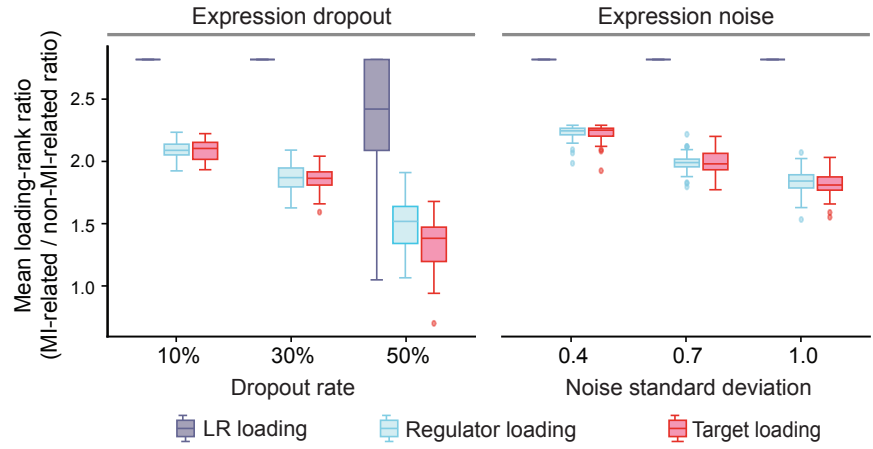

**Figure S2: Recovery of molecular anchors defining simulated meta-interactions.** Recovery of MI-associated LR pairs, sender regulators, and receiver targets across dropout and noise settings in 30 independent simulation replicates. Boxplots show the ratio of mean loading ranks for MI-associated versus non-associated features (larger ranks indicate higher loadings); values above 1 indicate preferential recovery. Ratios exceeded 1 for all loading types and settings (two-sided one-sample $t$-test against 1, all $P < 1 \times 10^{-8}$). Center lines indicate medians; boxes, interquartile ranges; whiskers, $1.5 \times \text{IQR}$; and points, outliers.

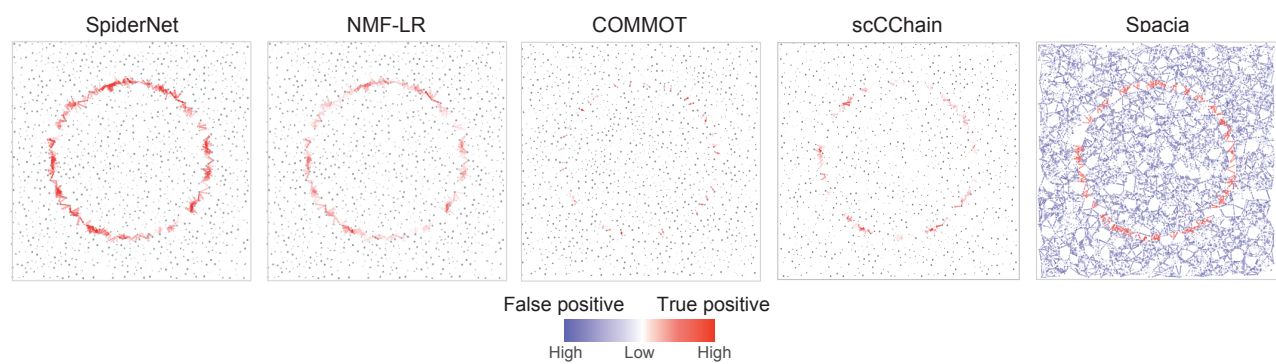

**Figure S3: *In situ* recovery of simulated meta-interaction structure across methods.** Spatial distributions of predicted interacting edges in a representative simulation replicate, shown after combining the MI-1 and MI-2 edge maps. Red indicates predicted edges assigned to the correct ground-truth MI, whereas blue indicates false positives, including non-interacting edges and edges assigned to the incorrect MI.

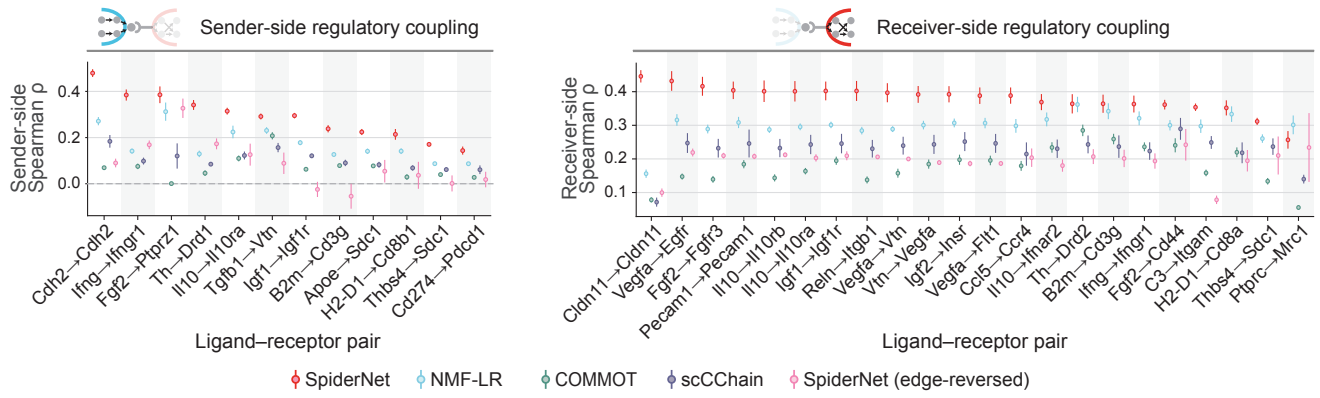

**Figure S4: Regulatory concordance of inferred communication features in the aging mouse brain MERFISH dataset.** Sender- and receiver-side regulatory coupling across curated ligand-receptor pairs. Coupling definitions, the *SpiderNet* (*edge-reversed*) control, and plotting conventions follow Fig. 2e. Spacia was omitted owing to computational constraints.

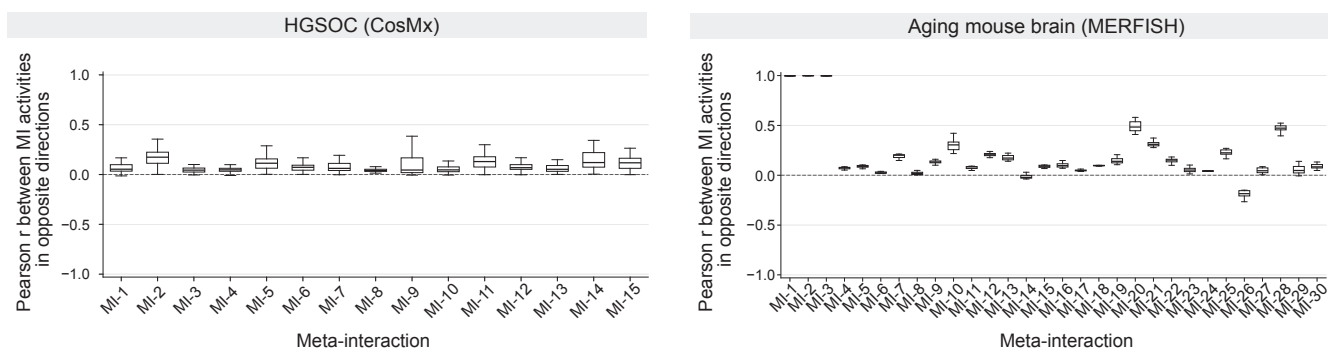

**Figure S5: Directional specificity of SpiderNet meta-interactions in HGSOc and aging mouse brain.** Within-slice Pearson correlations between MI activities inferred for the two directions of the same neighboring-cell pair in HGSOc CosMx (left) and aging mouse brain MERFISH (right). Only cell pairs with edges in both directions were included. Boxplots summarize correlations across tissue slices.

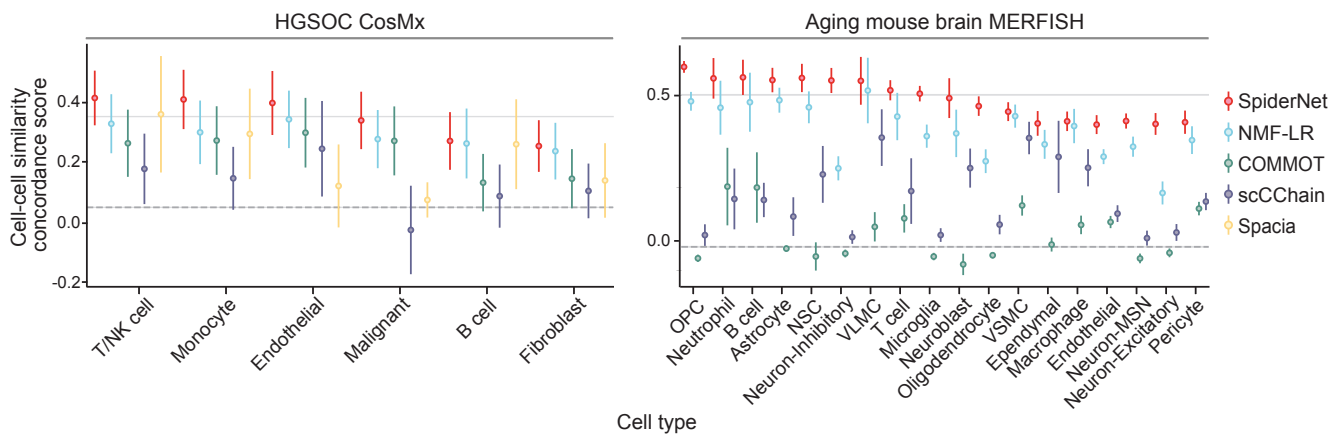

**Figure S6: Concordance between inferred communication states and transcriptional similarity.** Cell-cell similarity concordance scores for each cell type in HGSOC CosMx (left) and aging mouse brain MERFISH (right). Within each tissue slice, concordance was quantified as the Spearman correlation between cell-cell similarities derived from inferred communication states and normalized gene-expression profiles. Higher concordance indicates better preservation of within-cell-type transcriptional structure. Points and error bars show the mean ± s.d. across tissue slices. Spacia was omitted from the aging-brain analysis owing to computational constraints.

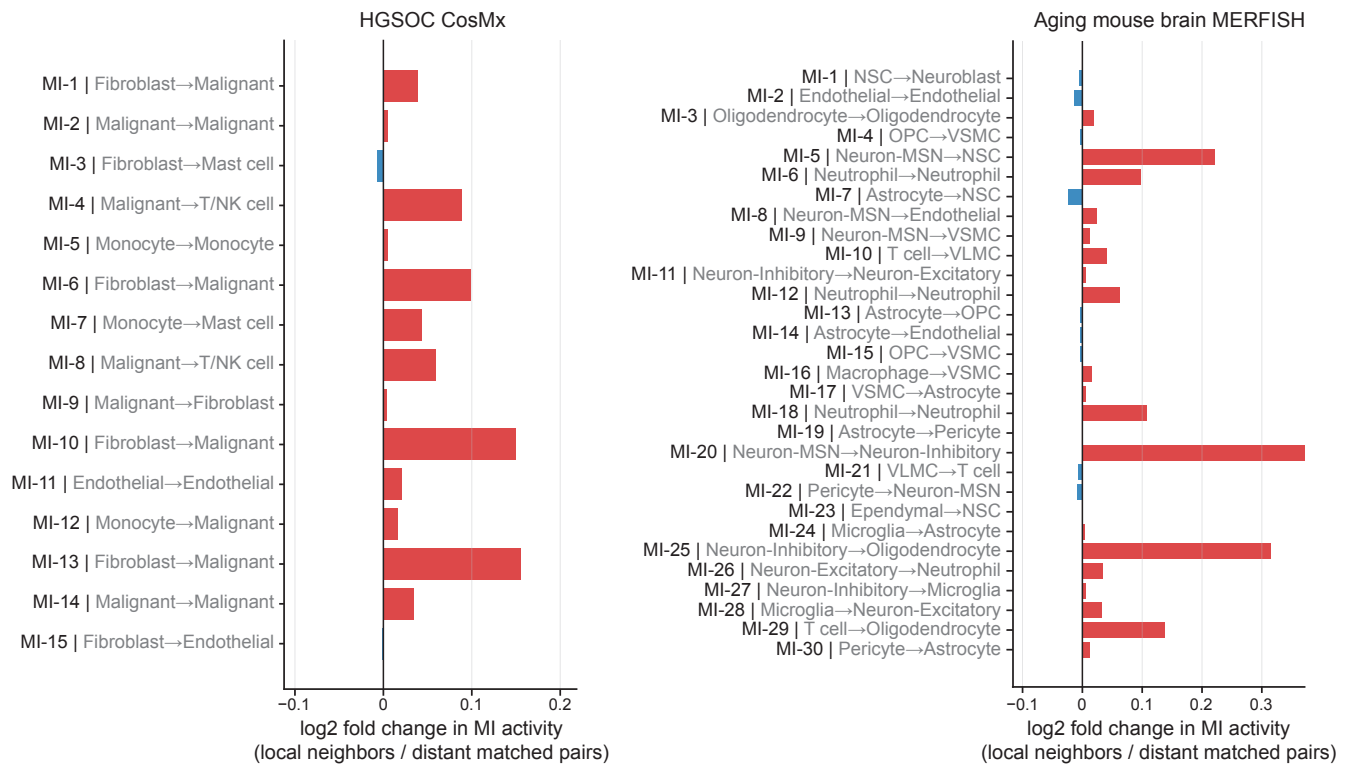

**Figure S7: Spatial specificity of dominant MI contexts in real tissues.** For each MI dimension, we selected the sender-receiver cell-type context with the highest mean MI activity and compared local neighboring cell pairs with matched distant pairs of the same cell types. Bars show the log<sub>2</sub> fold change in mean MI activity between local and distant pairs; positive values indicate enrichment among local neighboring cell pairs.

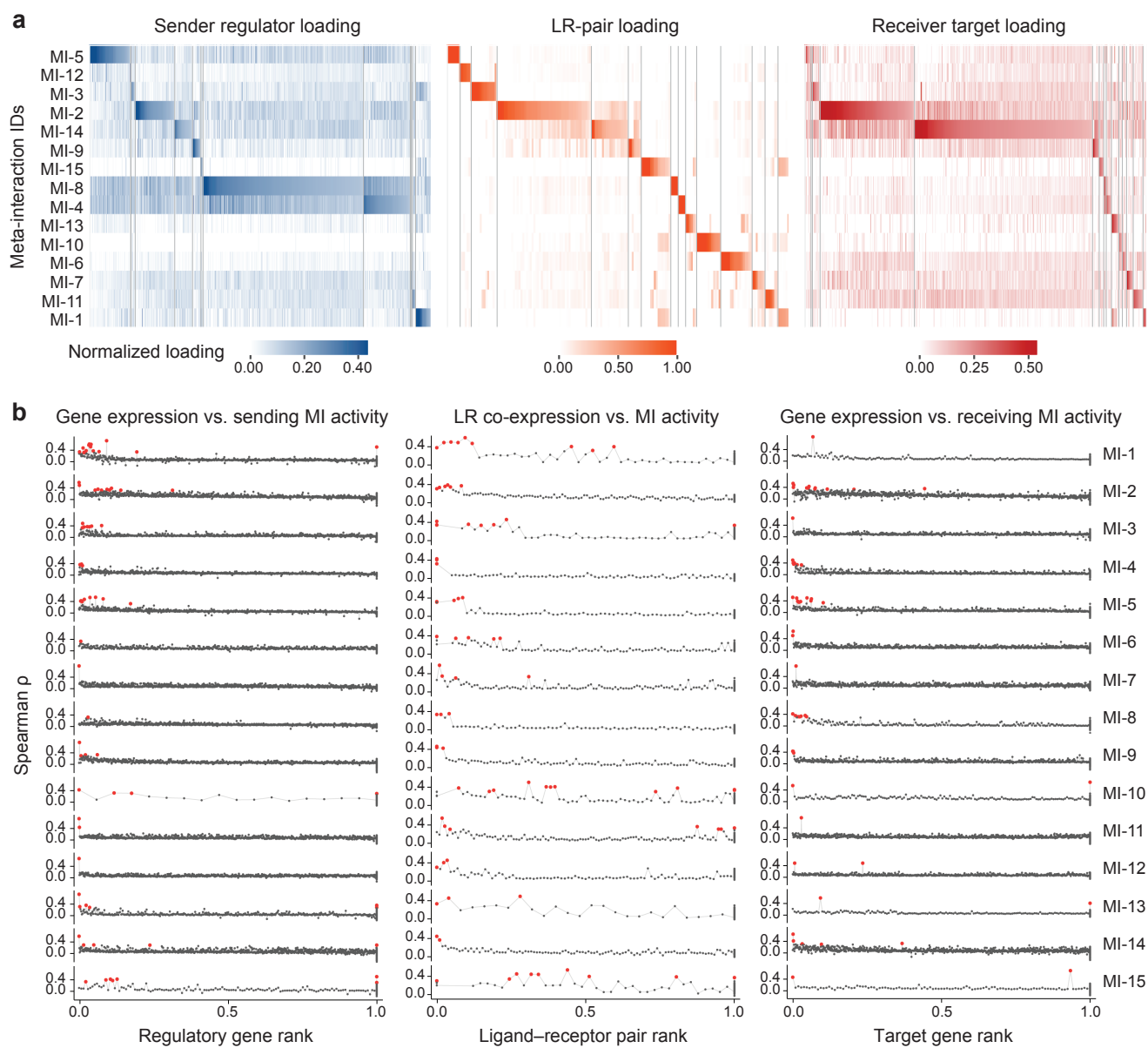

**Figure S8: Molecular anchoring and co-expression concordance of HGSOC meta-interactions.** **a**, Normalized sender-regulator, ligand-receptor (LR)-pair, and receiver-target loadings across MIs. Higher loadings indicate molecular features that more strongly define each sender-LR-receiver program. **b**, Concordance between MI activity and its molecular components. Shown are Spearman correlations of regulator-gene expression with sending MI activity (left), LR-pair co-expression with edge-level MI activity (middle), and target-gene expression with receiving MI activity (right), with features ordered by their corresponding loadings. Correlations greater than 0.4 are highlighted in red. The enrichment of stronger correlations among top-ranked features supports the molecular anchoring of SpiderNet MIs to coherent sender programs, LR bridges, and receiver programs.

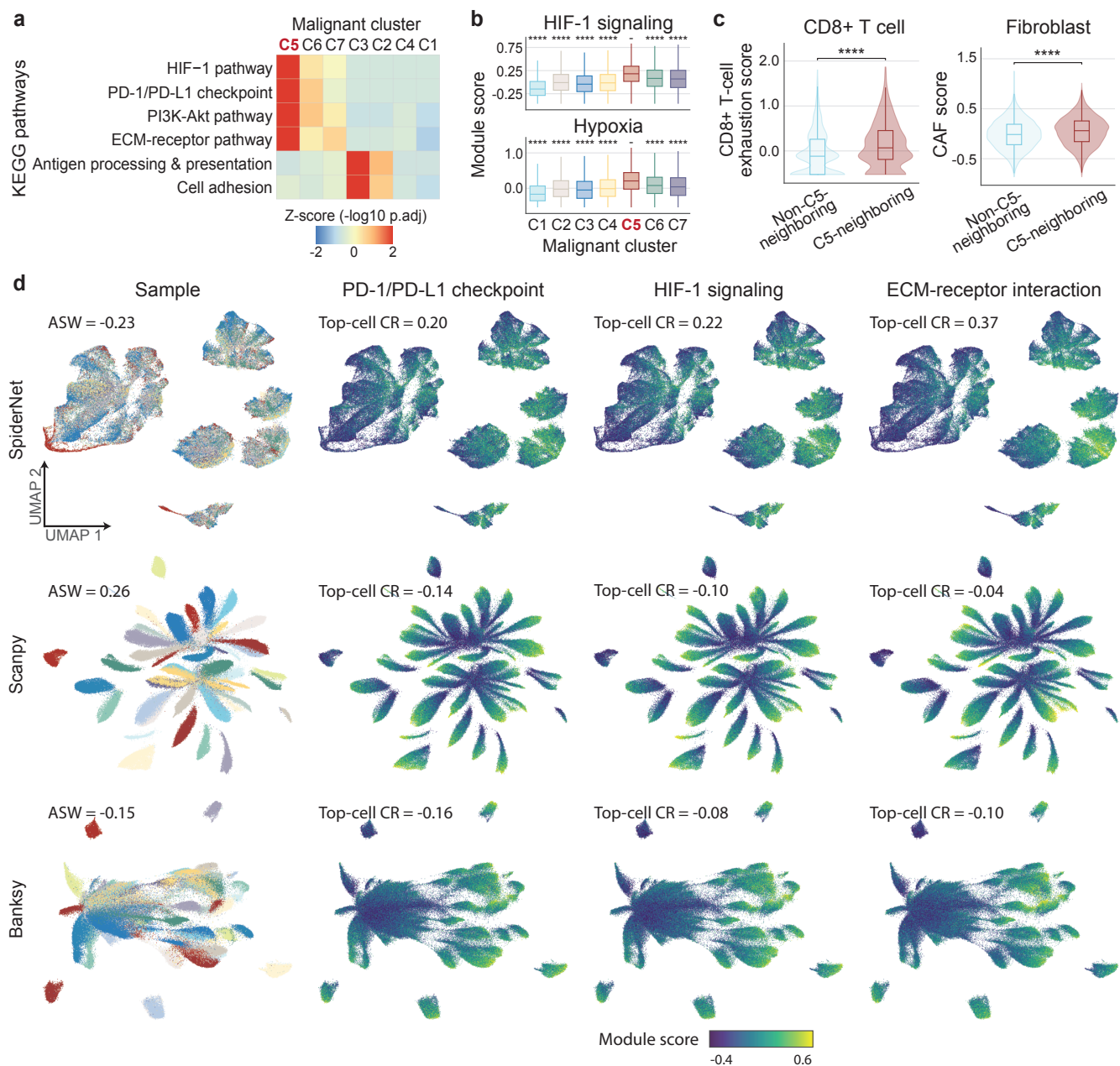

**Figure S9: Immune-suppressive features and embedding organization of the MI-defined malignant C5 state.** **a**, KEGG pathway enrichment of genes upregulated across MI-defined malignant clusters, shown as pathway-wise z scores of $-\log_{10}$ adjusted $P$ values. **b**, HIF-1 signaling and hypoxia scores across malignant clusters, with C5 compared with each other cluster. **c**, CD8<sup>+</sup> T-cell exhaustion and fibroblast CAF scores in cells neighboring versus not neighboring malignant C5 cells. **d**, Malignant-cell embeddings generated by SpiderNet, Scanpy, and Banksy. The first column shows sample identity; sample-based average silhouette width (ASW) quantifies sample-driven separation, with lower values indicating weaker sample dominance. The remaining columns show immune-suppressive program scores; top-cell compactness ratio (CR) is defined as $\log_2(D_{all}/D_{top})$, where $D_{top}$ and $D_{all}$ are the mean pairwise embedding distances among top-scoring and all cells, respectively; higher values indicate greater concentration of top-scoring cells. SpiderNet showed lower sample ASW and higher top-cell CRs than Scanpy and Banksy. For **b** and **c**, $P$ values are from one-sided Wilcoxon rank-sum tests testing for higher scores in C5 or C5-neighboring cells. Significance levels are denoted as \*$P < 0.05$; \*\*$P < 0.01$; \*\*\*$P < 0.001$; and \*\*\*\*$P < 0.0001$.

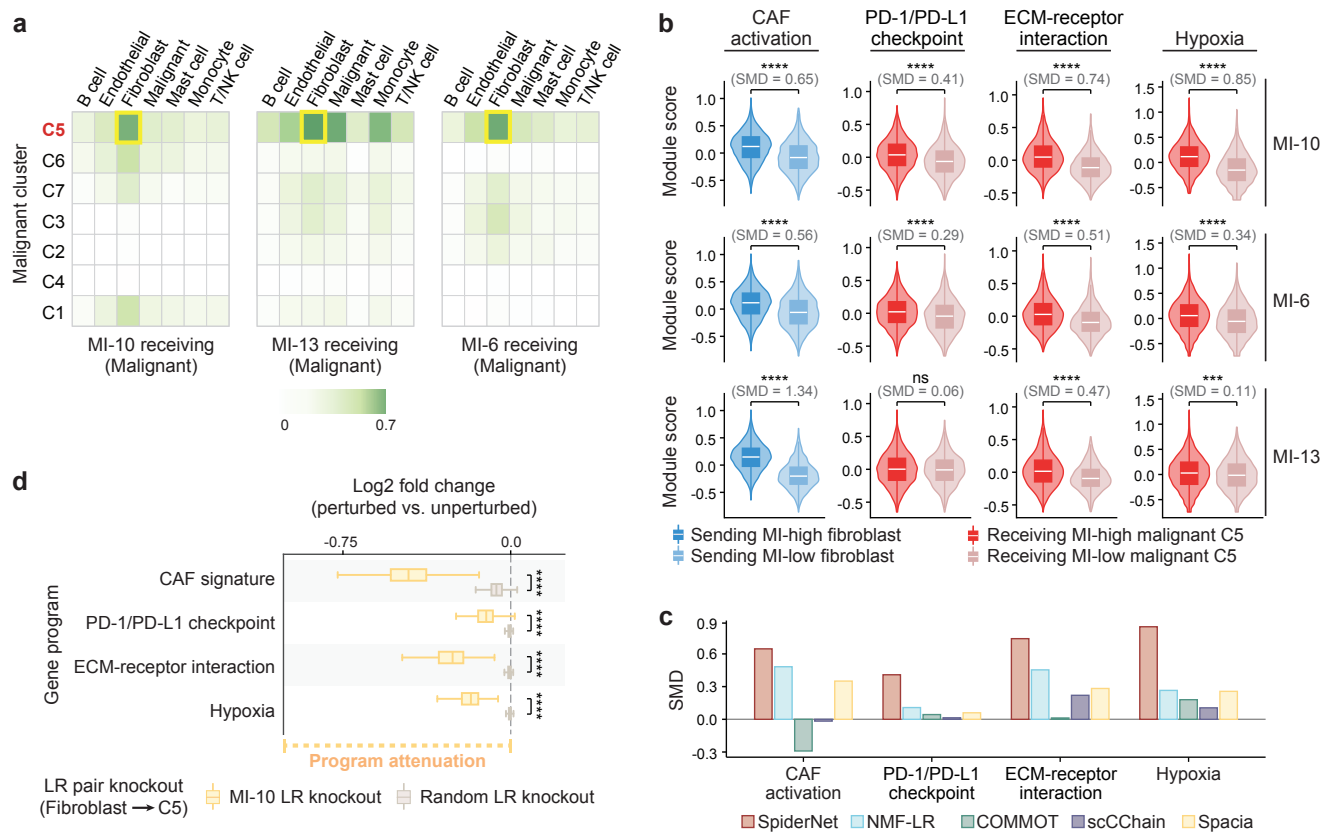

**Figure S10: Fibroblast-malignant MIs associated with the immune-suppressive malignant C5 state and fibroblast CAF activation.** **a**, Mean receiving strengths of MI-10, MI-13, and MI-6 across malignant clusters and sender cell types. Yellow boxes highlight fibroblast→C5 interactions. **b**, Fibroblast CAF and malignant program scores stratified by fibroblast→malignant activity of MI-10, MI-6, and MI-13. Fibroblasts were stratified by sending MI activity, whereas malignant C5 cells were stratified by receiving MI activity. $P$ values are from one-sided Wilcoxon rank-sum tests for higher scores in MI-high groups; standardized mean differences (SMDs) are shown. **c**, Comparison of CCC representations for the fibroblast→malignant C5 program. Bars show SMDs in CAF activation and malignant immune-suppressive programs for the best-matching communication feature from each method. **d**, *In silico* knockout of top MI-10 LR pairs in fibroblast→malignant C5 interactions, showing changes in predicted CAF activation and malignant immune-suppressive programs relative to random LR knockout. $P$ values are from two-sided Wilcoxon rank-sum tests comparing cell-level program changes between top MI-10 LR and random LR knockouts. Significance levels are denoted as ns, $P > 0.05$; \*\*\*$P < 0.001$; and \*\*\*\*$P < 0.0001$.

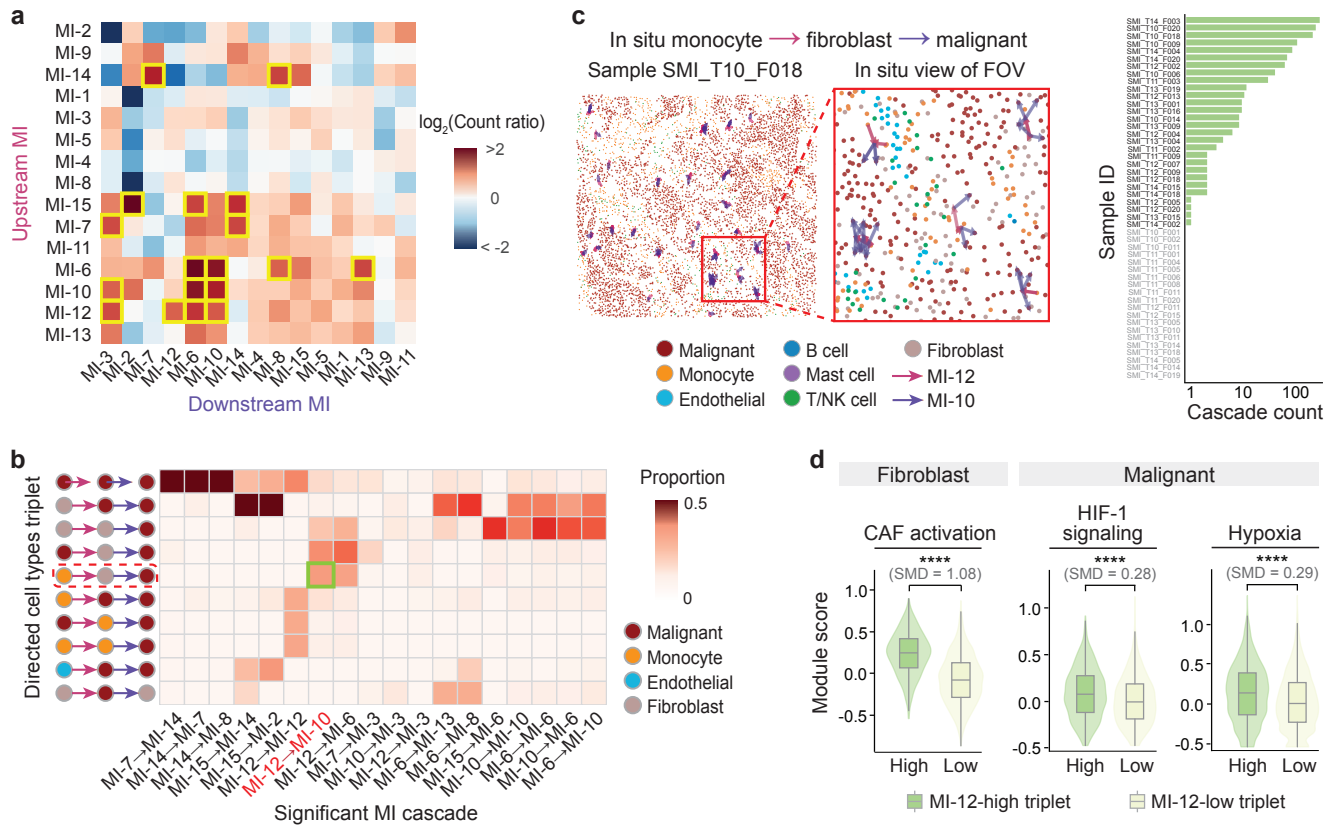

**Figure S11: MI-cascade analysis resolves an SPP1-THBS monocyte-fibroblast-malignant cascade.** **a**, Heatmap of $\log_2$ observed-to-expected cascade abundance ratios for upstream-downstream MI pairs. Cascades satisfying BH-adjusted empirical $P < 0.05$ and observed-to-expected count ratio $> 1.5$ are highlighted by yellow boxes. **b**, Cell-type composition of significant cascades, highlighting the monocyte→fibroblast→malignant MI-12→MI-10 cascade. **c**, Spatial and sample-level distribution of the monocyte→fibroblast→malignant MI-12→MI-10 cascade. Left, *in situ* distribution in representative sample SMI\_T10\_F018 with an enlarged field of view. Right, sample-wise counts of detected cascade triplets; the cascade was detected in 28 of 48 HGSOC samples. **d**, Topology-matched comparison of MI-12-high and MI-12-low monocyte-fibroblast-malignant triplets, defined by upstream monocyte→fibroblast MI-12 activity. Violin/box plots compare fibroblast CAF activation and malignant immune-suppressive programs. $P$ values are from two-sided Wilcoxon rank-sum tests, and standardized mean differences (SMDs) are shown. Significance is denoted as \*\*\*\* $P < 0.0001$.

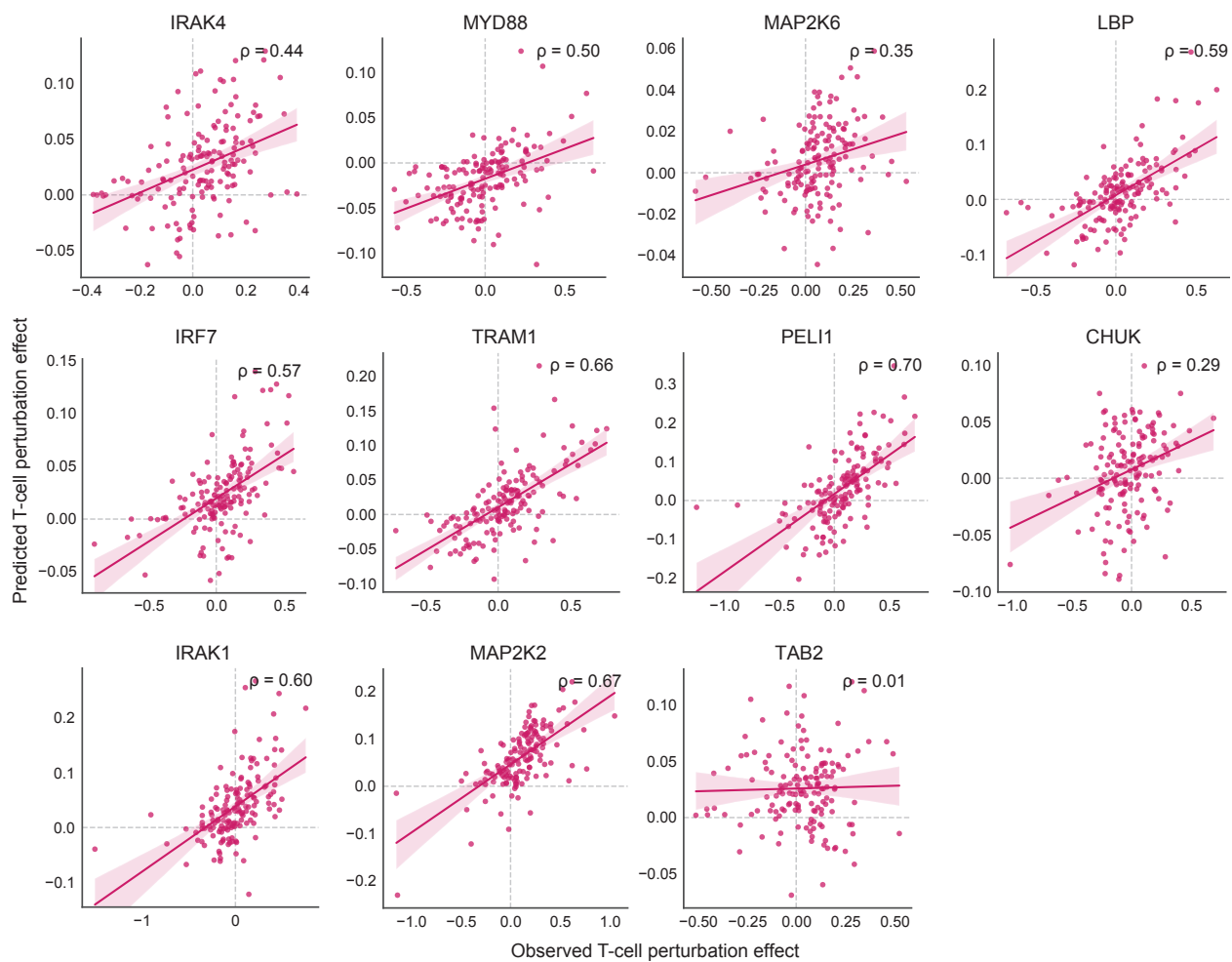

**Figure S12: Gene-level prediction of T-cell responses to held-out melanoma NF-κB perturbations.** Scatter plots compare observed and SpiderNet-predicted T-cell responses across measured genes for each held-out melanoma perturbation. Points represent measured genes; lines show linear fits with confidence intervals, and $\rho$ denotes Spearman correlation.

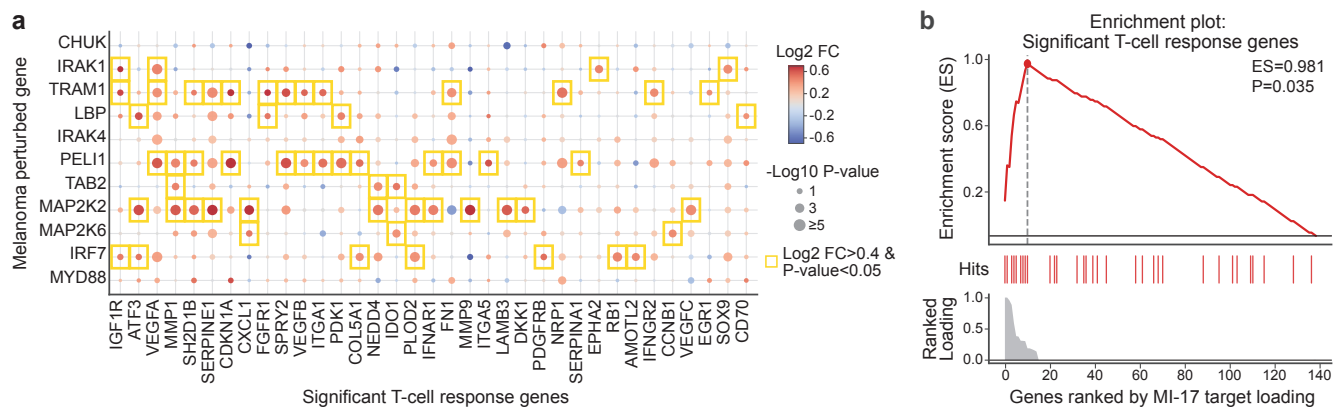

**Figure S13: MI-17 receiver-target loadings prioritize T-cell response genes induced by melanoma NF-κB perturbations.** **a**, T-cell response genes across melanoma-cell NF-κB perturbations, comparing T cells neighboring melanoma cells carrying each perturbation with T cells not neighboring such cells. Dot color indicates $\log_2$ fold change, and dot size indicates $-\log_{10} P$ from two-sided Wilcoxon rank-sum tests; yellow boxes mark genes with $\log_2$ fold change > 0.4 and $P < 0.05$. **b**, Gene set enrichment analysis (GSEA) of significant T-cell response genes among genes ranked by MI-17 receiver-target loading, supporting alignment of MI-17 with the perturbation-induced T-cell transcriptional program.

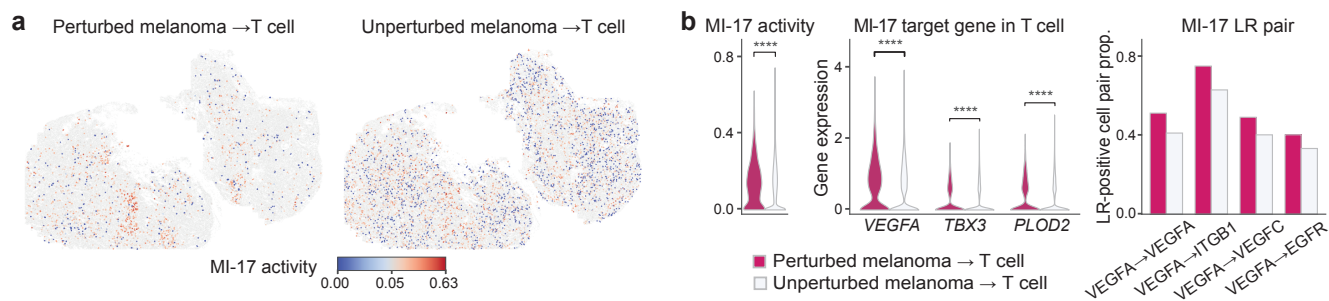

**Figure S14: Observed enrichment of MI-17 activity and associated molecular programs in NF-κB-perturbed melanoma-T-cell interactions.** **a**, *In situ* distribution of melanoma→T-cell MI-17 interactions involving NF-κB-perturbed or unperturbed melanoma cells. **b**, Comparison between perturbed and unperturbed melanoma-T-cell pairs in MI-17 activity, MI-17 receiver-target expression in T cells, and the proportion of cell pairs with positive co-expression for each top MI-17 LR pair. *P* values are from one-sided Wilcoxon rank-sum tests for higher values in perturbed pairs; \*\*\*\**P*<0.0001.

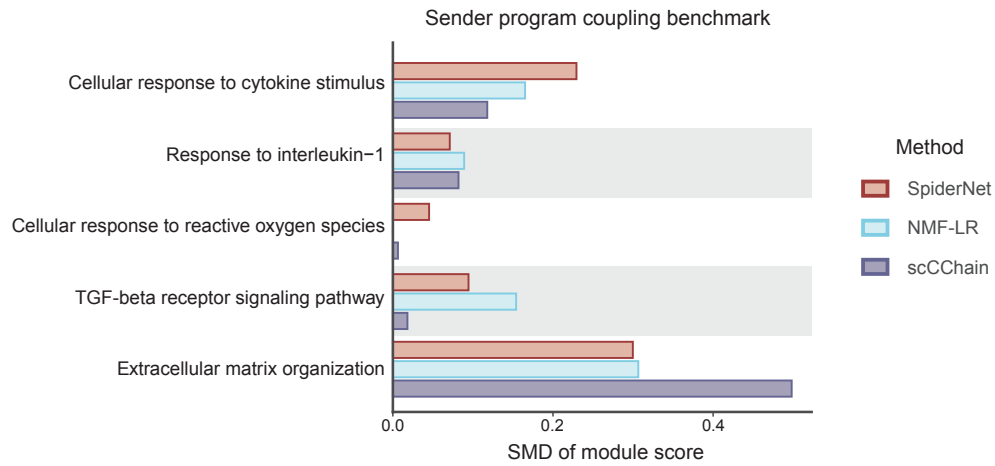

**Figure S15: Sender-program coupling for each method's most perturbation-responsive melanoma-to-T-cell communication feature.** For each method, the communication feature with the largest mean increase across the 11 NF-κB perturbations was selected. Coupling to MI-17-associated melanoma sender programs was quantified as the standardized mean difference (SMD) in module scores between high- and low-communication-activity cells; positive values indicate stronger coupling.

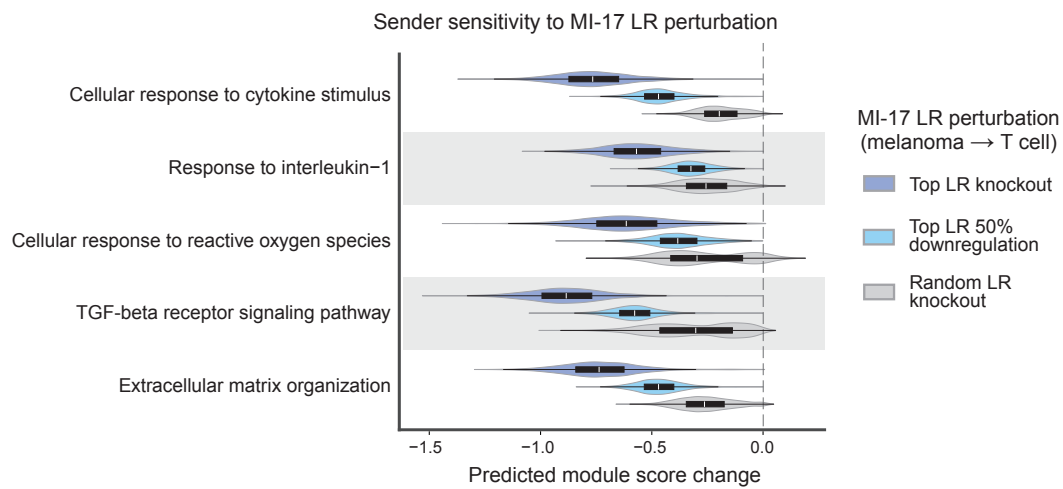

**Figure S16: Model-predicted sender sensitivity of MI-17 sender programs in melanoma cells to perturbation of top ligand-receptor (LR) pairs.** Top MI-17 LR pairs were knocked out or downregulated by 50% and compared with size-matched random LR knockouts; negative values indicate reduced predicted sender-program scores.

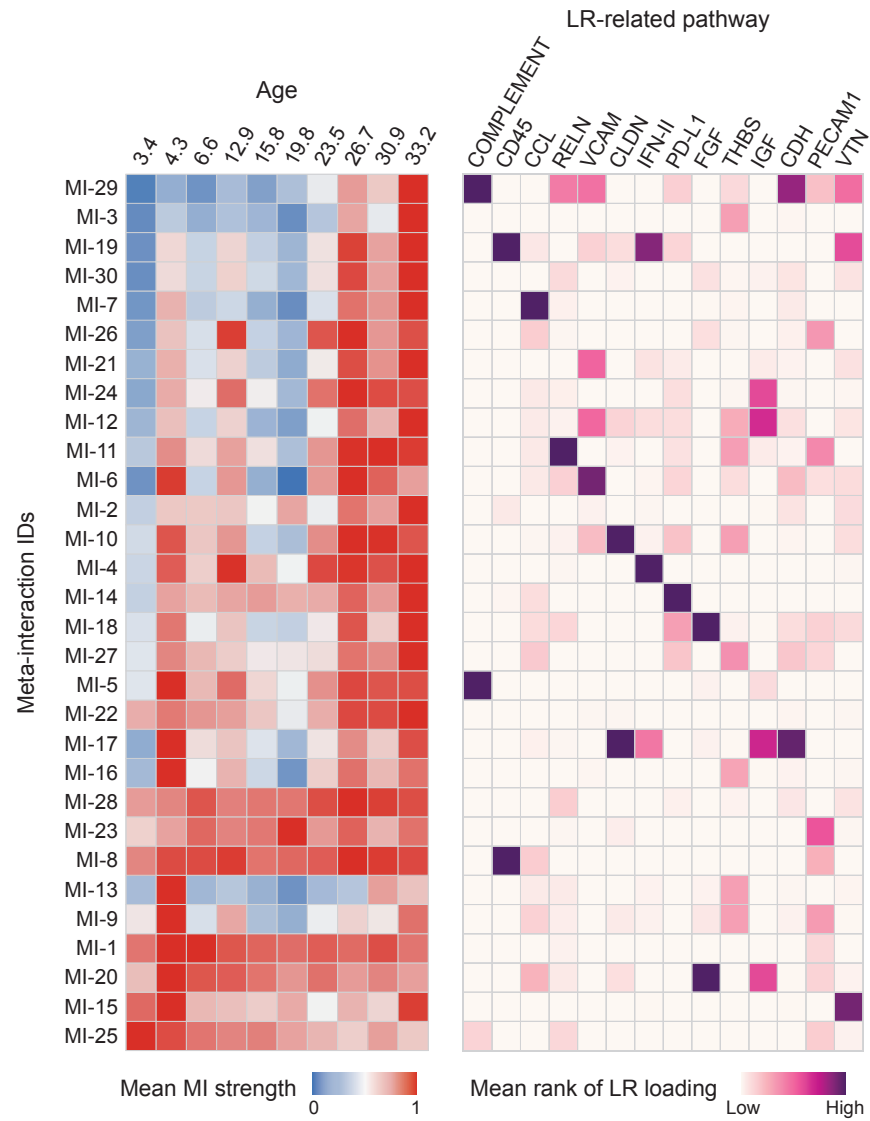

**Figure S17: Age-associated patterns and ligand-receptor pathway enrichment of meta-interactions in the aging mouse brain.** Left, mean MI activity across samples ordered by age. Right, mean rank of LR loadings summarized by CellChat pathways for each MI (abbreviations from CellChat). Color denotes the mean rank of pathway-related LR loadings within each MI.

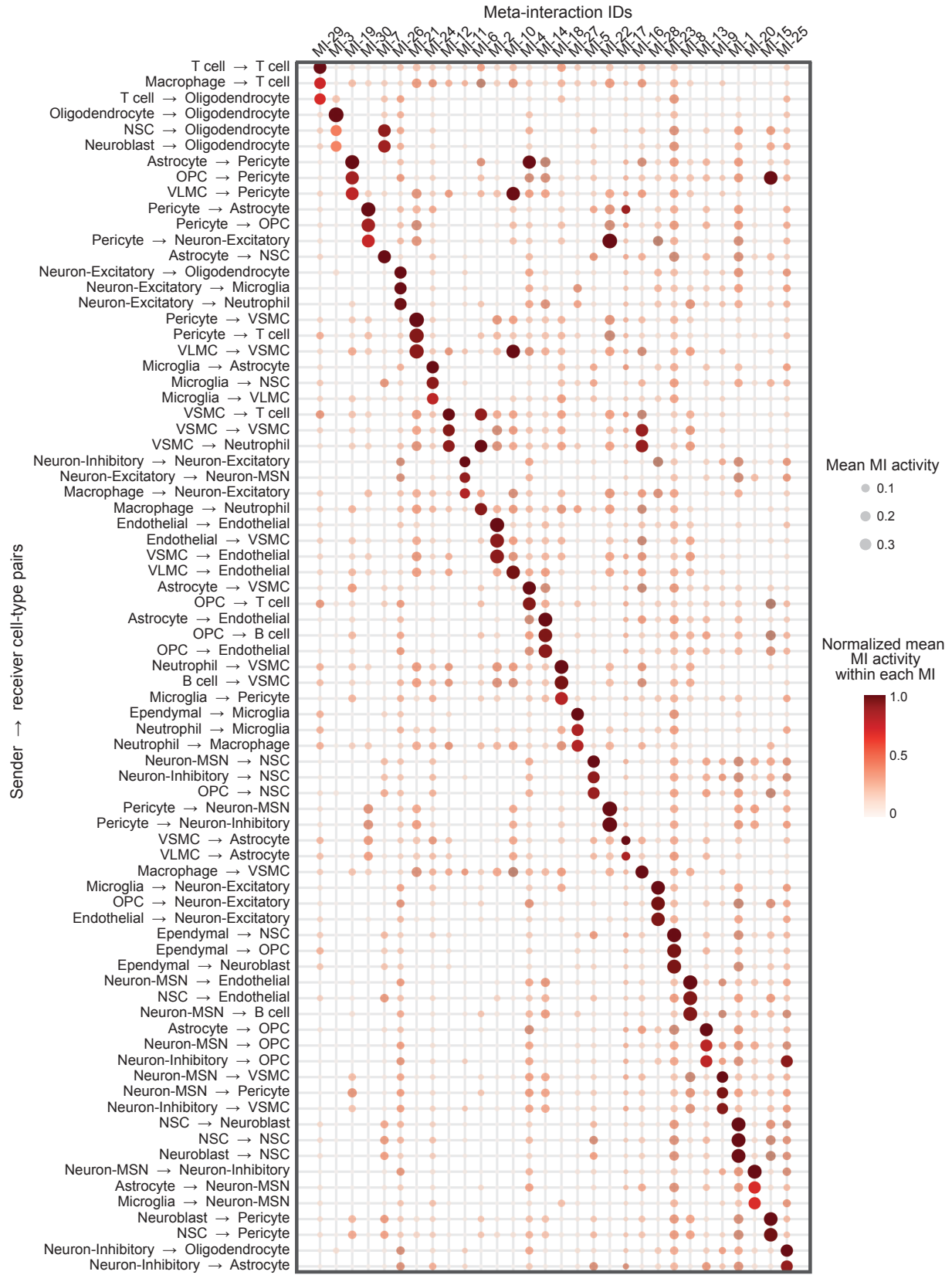

**Figure S18: Cell-type-pair specificity of aging-brain meta-interactions.** Mean MI activity across directed sender-receiver cell-type pairs in the coronal aging mouse brain atlas. Dot size indicates raw mean activity, whereas color indicates activity normalized within each MI. The top three highest-activity cell-type pairs are highlighted for each MI.

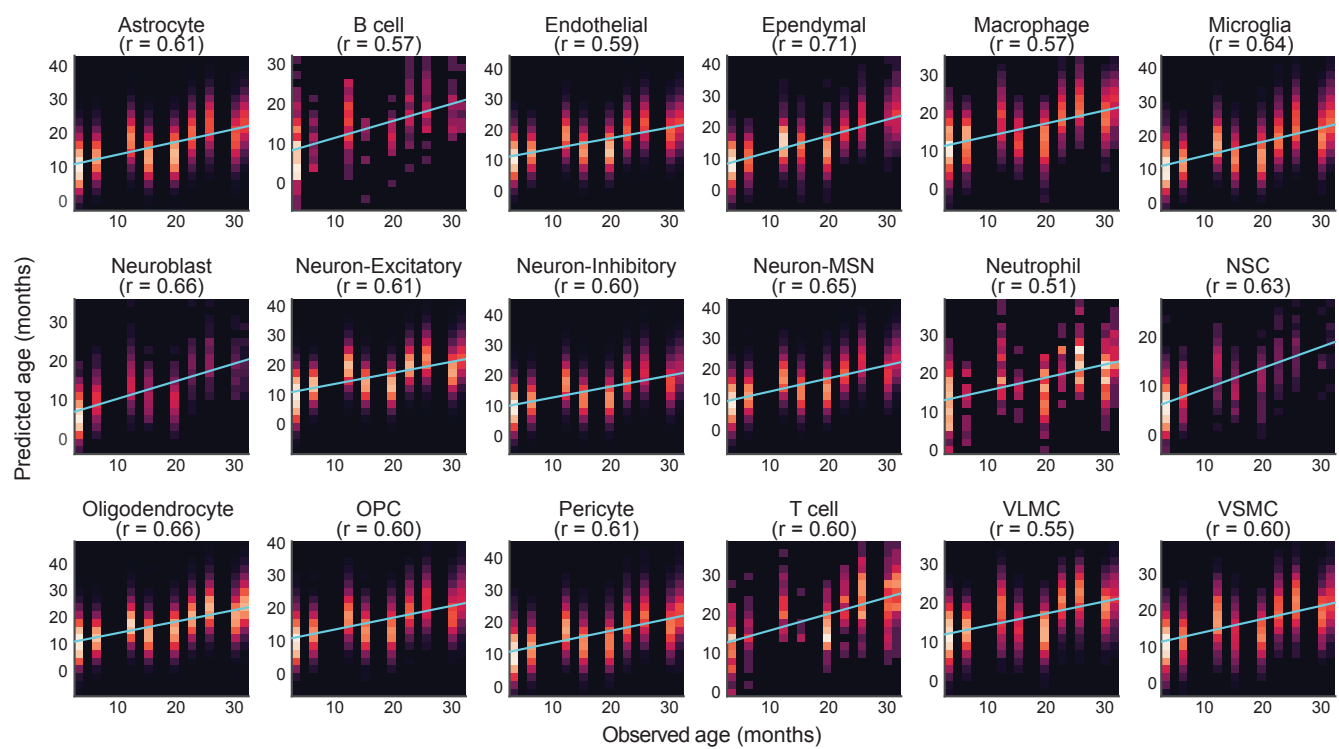

**Figure S19: Cell-type-specific, cell-level age prediction from SpiderNet MI profiles in the MERFISH coronal brain atlas.** Each panel compares observed and predicted age for one cell type. Predictions were generated by 10-fold cross-validation using cell-type-specific linear models fitted to SpiderNet cell-level MI profiles. Heatmaps show cell density, light-blue lines indicate least-squares fits, and $r$ denotes the Pearson correlation between observed and predicted age.

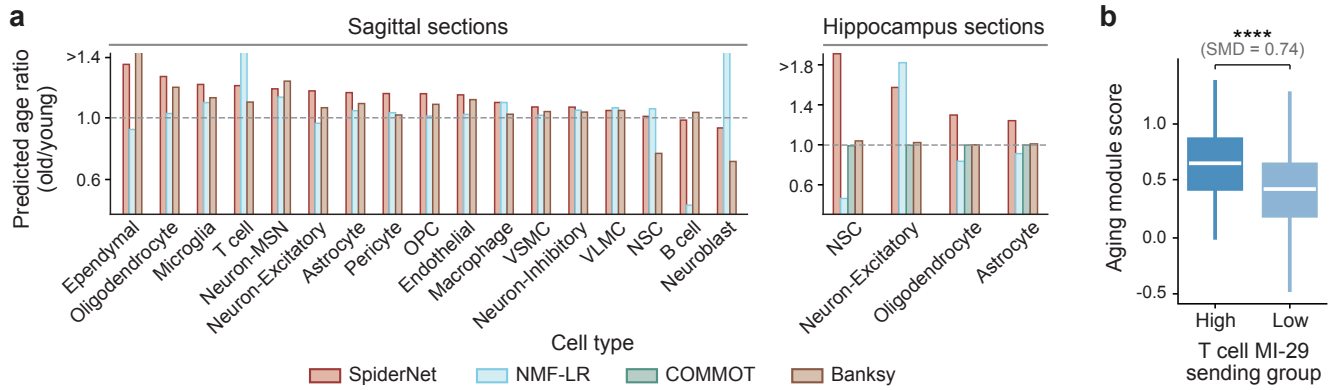

**Figure S20: Cross-context transfer of age-prediction models and MI-29-associated T-cell aging signals.** **a**, Old-to-young ratios of mean predicted age across cell types in sagittal MERFISH (left) and hippocampus Stereo-seq (right). Age-prediction models trained on coronal MERFISH were applied without retraining; ratios above 1 indicate higher predicted age in old samples. Young and old groups were defined as $\leq 19$ and $> 19$ months for sagittal MERFISH and by the original study labels for Stereo-seq. **b**, T-cell aging scores stratified by high versus low MI-29 sending activity in sagittal sections. $P$ values are from two-sided Wilcoxon rank-sum tests; standardized mean differences (SMDs) and significance levels (\*\*\*\*$P < 0.0001$) are shown. COMMOT was omitted for sagittal MERFISH owing to computational constraints, and scCChain was omitted because pretrained application to new datasets was not supported.

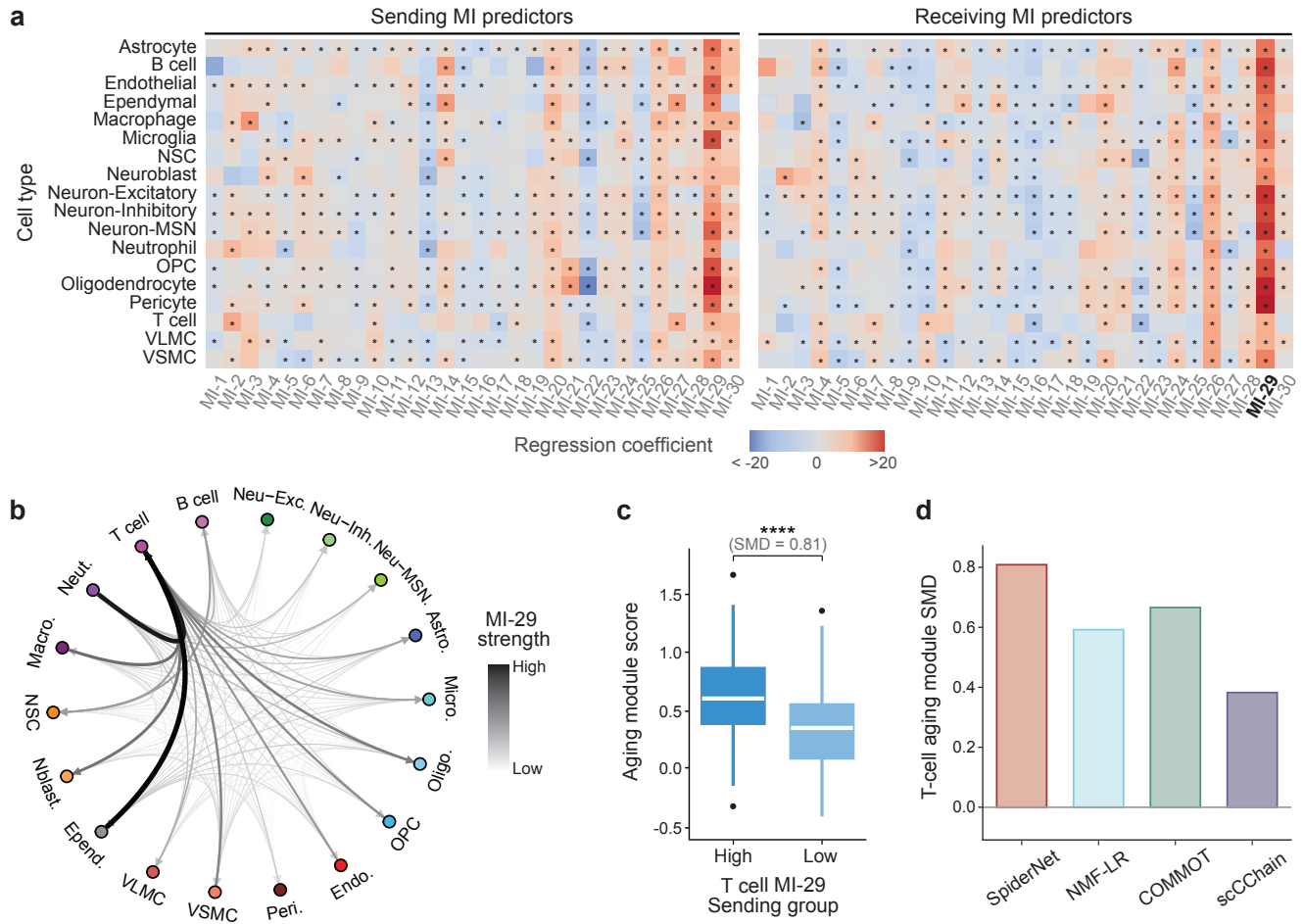

**Figure S21: MI-29-mediated T-cell communication is associated with local tissue aging across brain cell types.** **a**, Regression coefficients from cell-type-specific linear models predicting cellular age from aggregated sending and receiving MI features. Asterisks mark significant MI predictors ($P < 0.05$). Receiving MI-29 showed consistently positive age associations across cell types. **b**, Circle plot showing MI-29 signals between cell-type pairs. **c**, T-cell aging scores in high- and low-MI-29-sending T cells, defined by median MI-29 sending activity. $P$ values are from two-sided Wilcoxon rank-sum tests; standardized mean differences (SMDs) are shown. \*\*\*\*$P < 0.0001$. **d**, Benchmark of T-cell aging-program coupling across CCC representations. For each method, T cells were stratified by the T-cell-sending communication feature most strongly associated with the aging score. Bars show standardized mean differences (SMDs) between high- and low-communication T cells; for SpiderNet, this feature is MI-29.

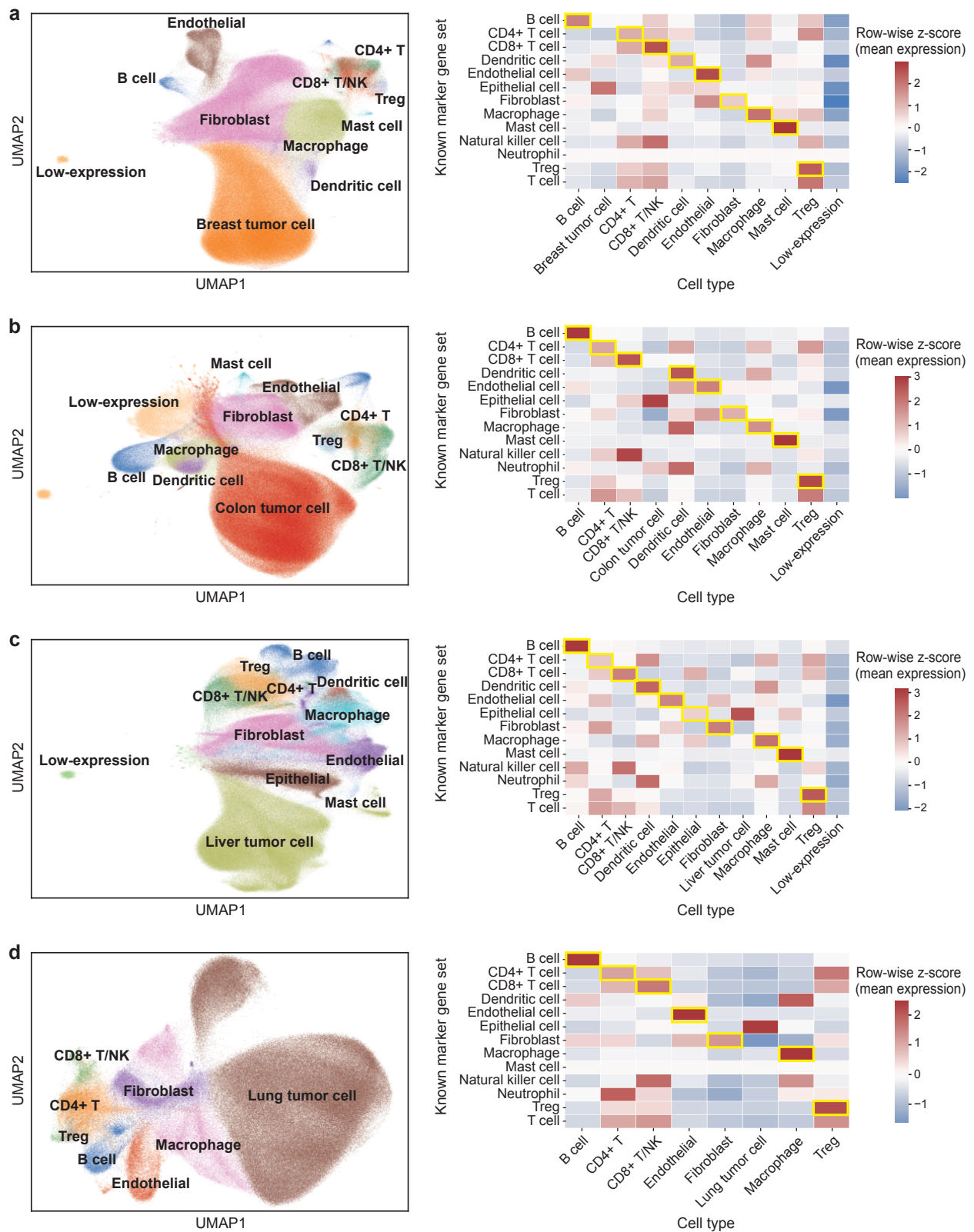

**Figure S22: Cell-type annotation and marker validation across eight cancer types.** a-d, Cell-type annotations in breast, colon, liver, and lung cancers. UMAPs show cells colored by annotated cell type (left), and heatmaps show row-wise z-scored mean expression of canonical marker-gene sets across cell types (right). Yellow boxes mark the expected correspondence between each marker-gene set and its annotated cell type.

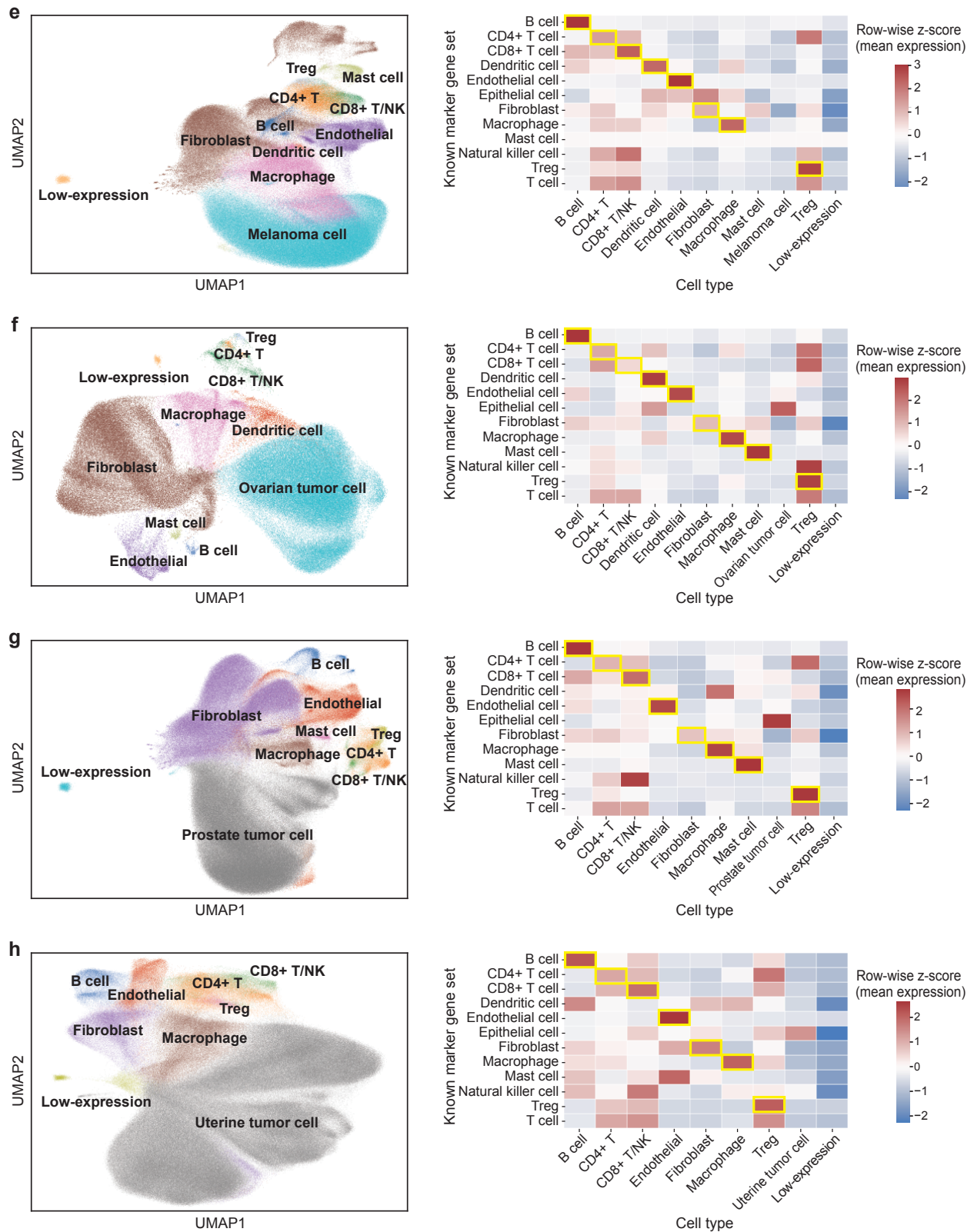

**Figure S22: Cell-type annotation and marker validation across eight cancer types (continued).** e-h, Cell-type annotations in melanoma, ovarian, prostate, and uterine cancers. UMAPs show cells colored by annotated cell type (left), and heatmaps show row-wise z-scored mean expression of canonical marker-gene sets across cell types (right). Yellow boxes mark the expected correspondence between each marker-gene set and its annotated cell type.

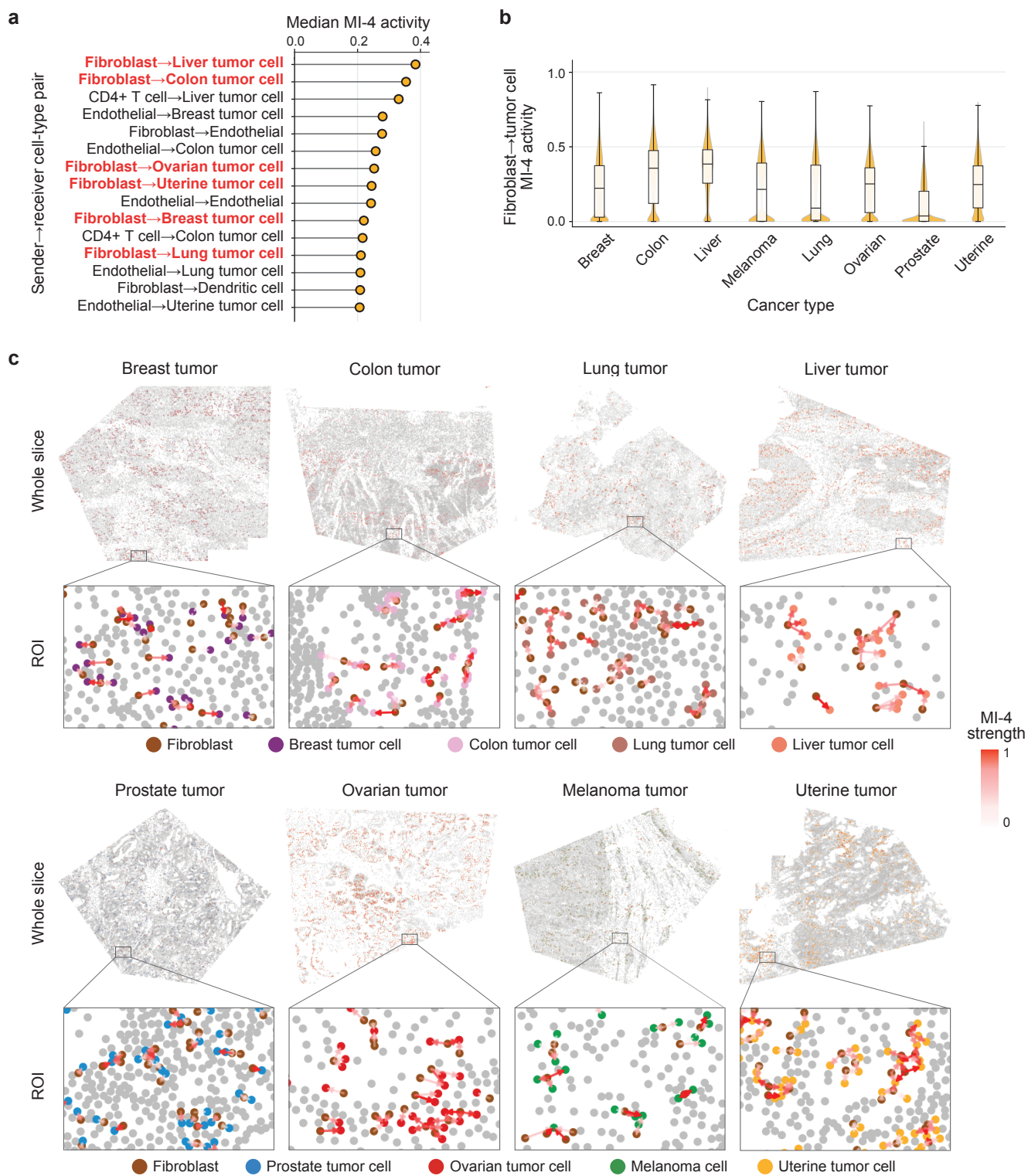

**Figure S23: Pan-cancer recurrence and *in situ* organization of fibroblast-to-tumor MI-4.** **a**, Top directed cell-type pairs ranked by their median MI-4 activity across spatial sub-slices. Fibroblast→tumor pairs are highlighted in red. **b**, Distribution of fibroblast→tumor MI-4 activity across cancer types. **c**, *In situ* distribution of fibroblast→tumor MI-4 interactions in representative sub-slices and enlarged regions of interest. Edges are colored by MI-4 activity, with fibroblasts and tumor cells highlighted.

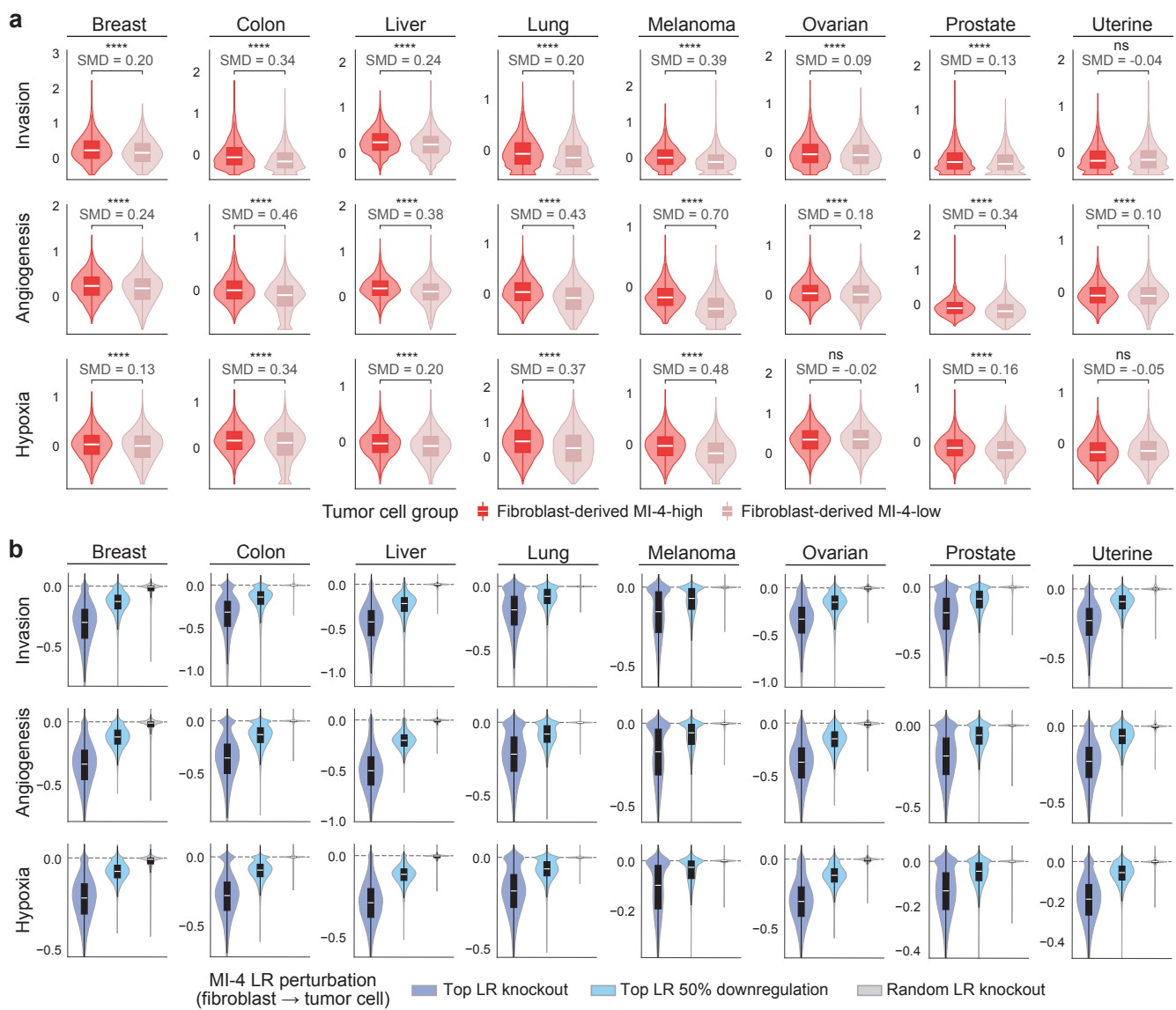

**Figure S24: Fibroblast-derived MI-4 is associated with malignant programs and shows fixed-model counterfactual sensitivity across cancer types.** **a**, Tumor cells were stratified within each cancer type by aggregated incoming fibroblast→tumor MI-4 activity. Violin/box plots compare invasion, angiogenesis, and hypoxia scores between MI-4-high and MI-4-low tumor cells. $P$ values are from Wilcoxon rank-sum tests; standardized mean differences (SMDs) are shown. Significance is denoted as ns, $P>0.05$; \*\*\*\*$P<0.0001$. **b**, Fixed-model *in silico* perturbation of fibroblast→tumor MI-4 LR pairs. Plots show changes in receiver tumor-cell program scores after knockout or 50% downregulation of top MI-4 LR pairs, compared with size-matched random LR-pair knockout, across cancer types. Top LR knockout produced greater predicted program attenuation than random LR knockout across all cancer-type-program comparisons (two-sided Wilcoxon signed-rank tests, all $P<0.0001$).

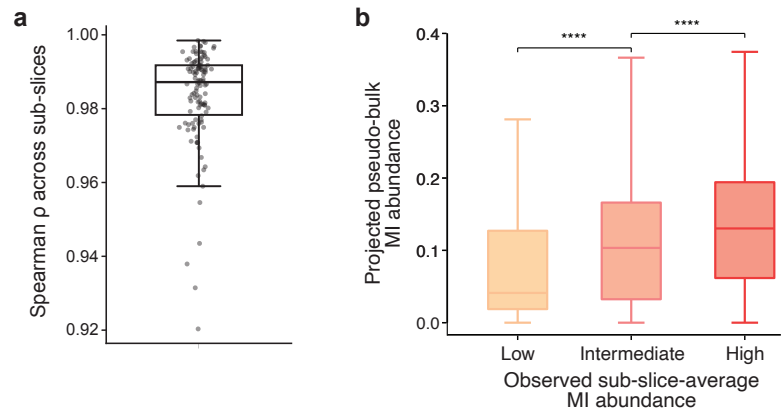

**Figure S25: Pseudo-bulk validation of projected MI abundance.** **a**, Spearman correlations between pseudo-bulk LR co-expression proxies and spatial LR co-expression averaged over cell-cell edges across sub-slices. Each point represents one LR pair. **b**, Projected pseudo-bulk MI abundance across tertiles of observed sub-slice-average MI abundance, pooling all MI–sub-slice combinations. $P$ values indicate two-sided Wilcoxon rank-sum tests between adjacent tertiles. \*\*\*\*$P < 0.0001$.

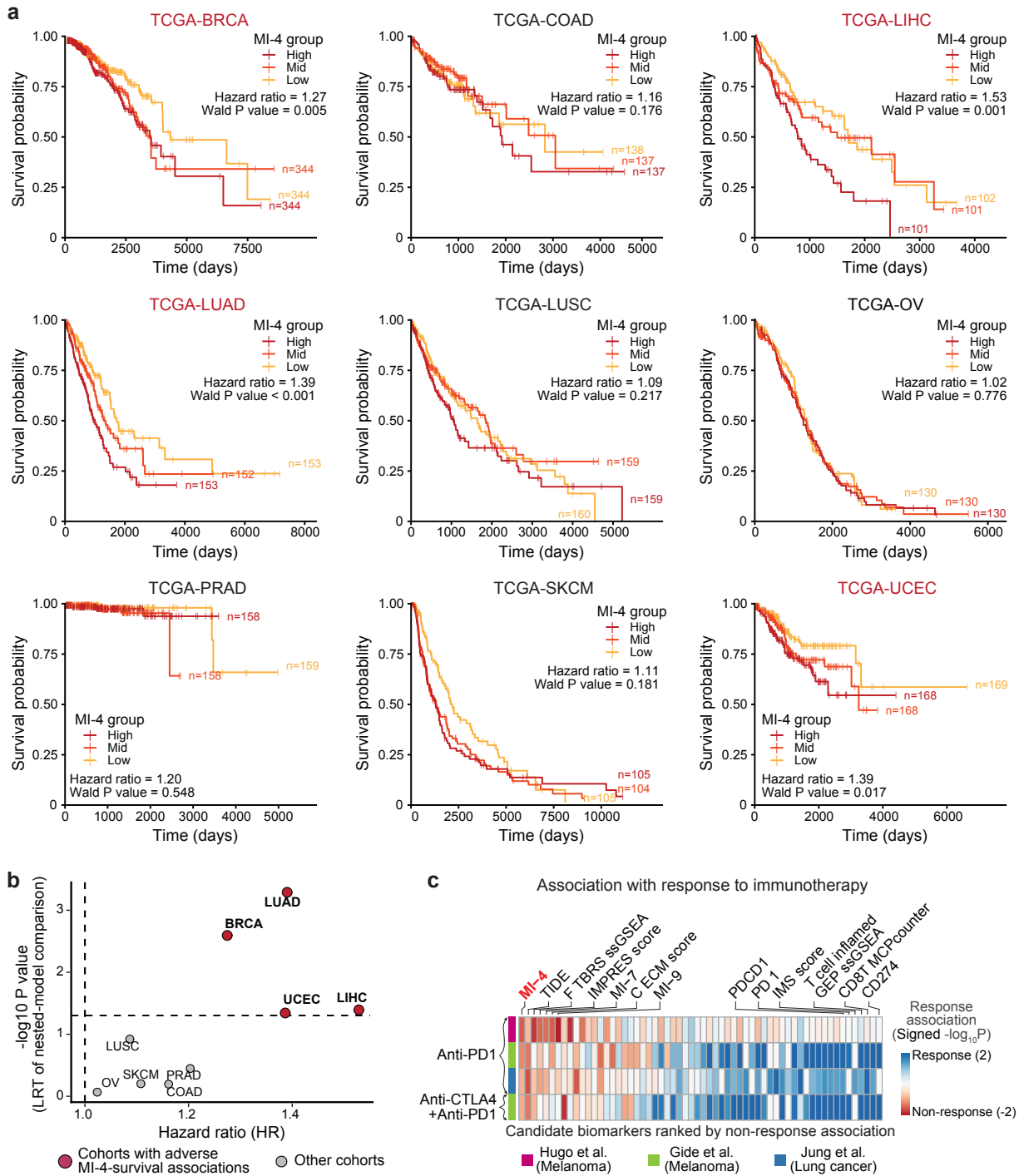

**Figure S26: Projected MI-4 abundance is associated with survival and immunotherapy response across cancer cohorts.** **a**, Kaplan-Meier curves of overall survival across nine TCGA cohorts, with GDC-defined tumor specimens stratified by projected MI-4 abundance. Multivariable Cox models tested associations between standardized MI-4 abundance and overall survival while adjusting for clinical covariates. Cohorts with adverse associations ($HR > 1$, $Wald P < 0.05$) are highlighted in red. **b**, Likelihood-ratio tests evaluating whether MI-4 adds prognostic information beyond clinical covariates and MI-4-associated invasion, angiogenesis, and hypoxia programs. Points show MI-4 hazard ratios and nested-model significance; dashed lines indicate $HR = 1$ and $P = 0.05$. **c**, Associations of projected MI-4 abundance and established immunotherapy biomarkers with checkpoint-blockade response in melanoma and lung cancer cohorts [20–22]. Within each cohort-treatment setting, responder and non-responder groups were compared using two-sided Wilcoxon rank-sum tests, with associations shown as signed $-\log_{10} P$ values. Blue indicates responder-enriched associations, whereas red indicates non-responder-enriched associations. Candidates are ordered by their mean within-setting rank of non-response association across treatment settings.

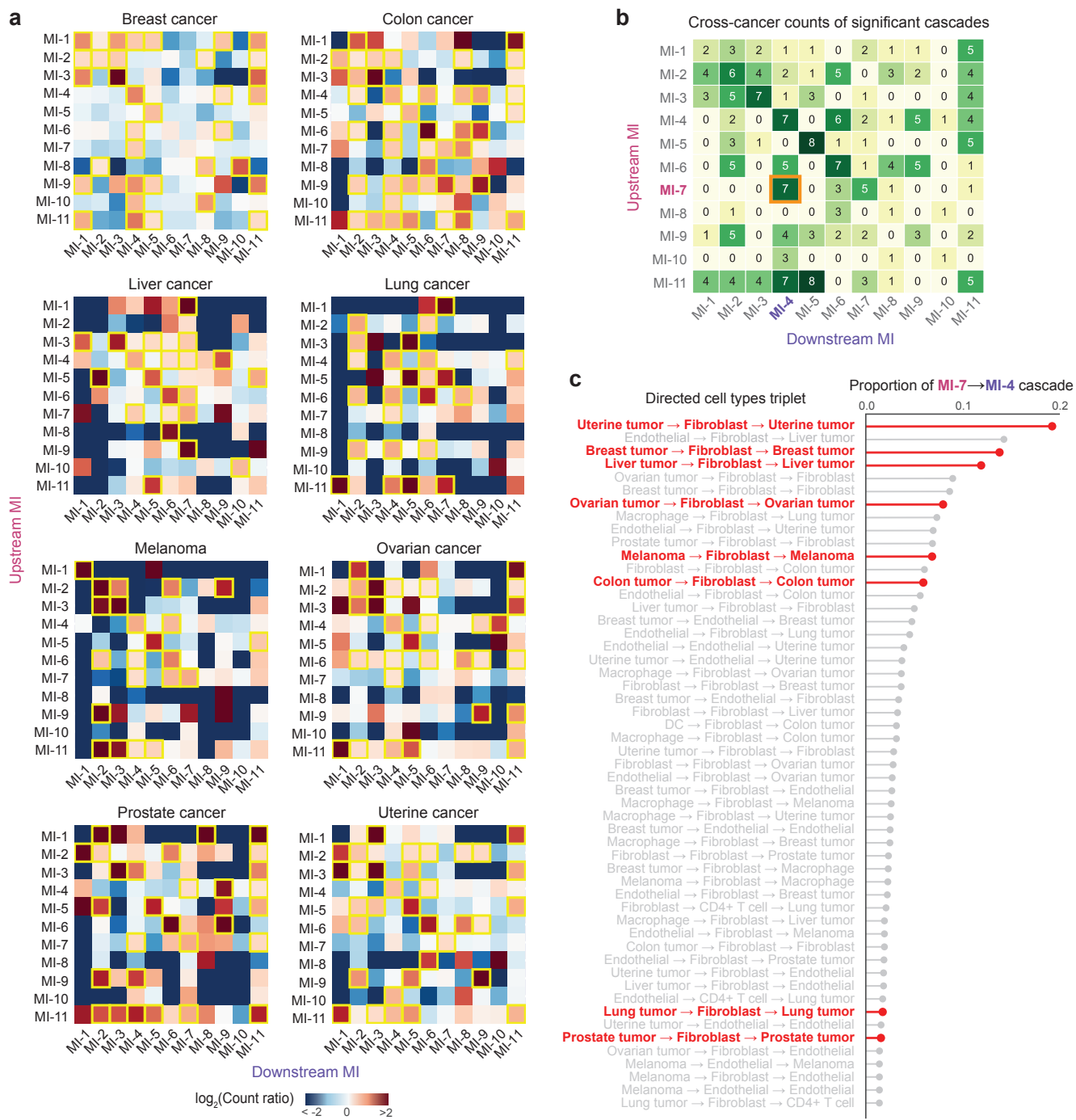

**Figure S27: MI cascades reveal recurrent higher-order signaling and a pan-cancer tumor-fibroblast-tumor relay.** **a**, Heatmaps showing the $\log_2$ ratio of observed-to-expected cascade counts for each upstream-downstream MI pair across eight cancer types. Cascades satisfying BH-adjusted empirical $P < 0.05$ and observed-to-expected count ratio $> 1.5$ are highlighted by yellow boxes. **b**, Number of cancer types in which each upstream-downstream MI pair was significant in panel **a**. The recurrent MI-7→MI-4 cascade is outlined in orange. **c**, Ranked proportions of MI-7→MI-4 cascades across directed cell-type triplets. Recurrent tumor→fibroblast→tumor relays across cancer types are highlighted in red.

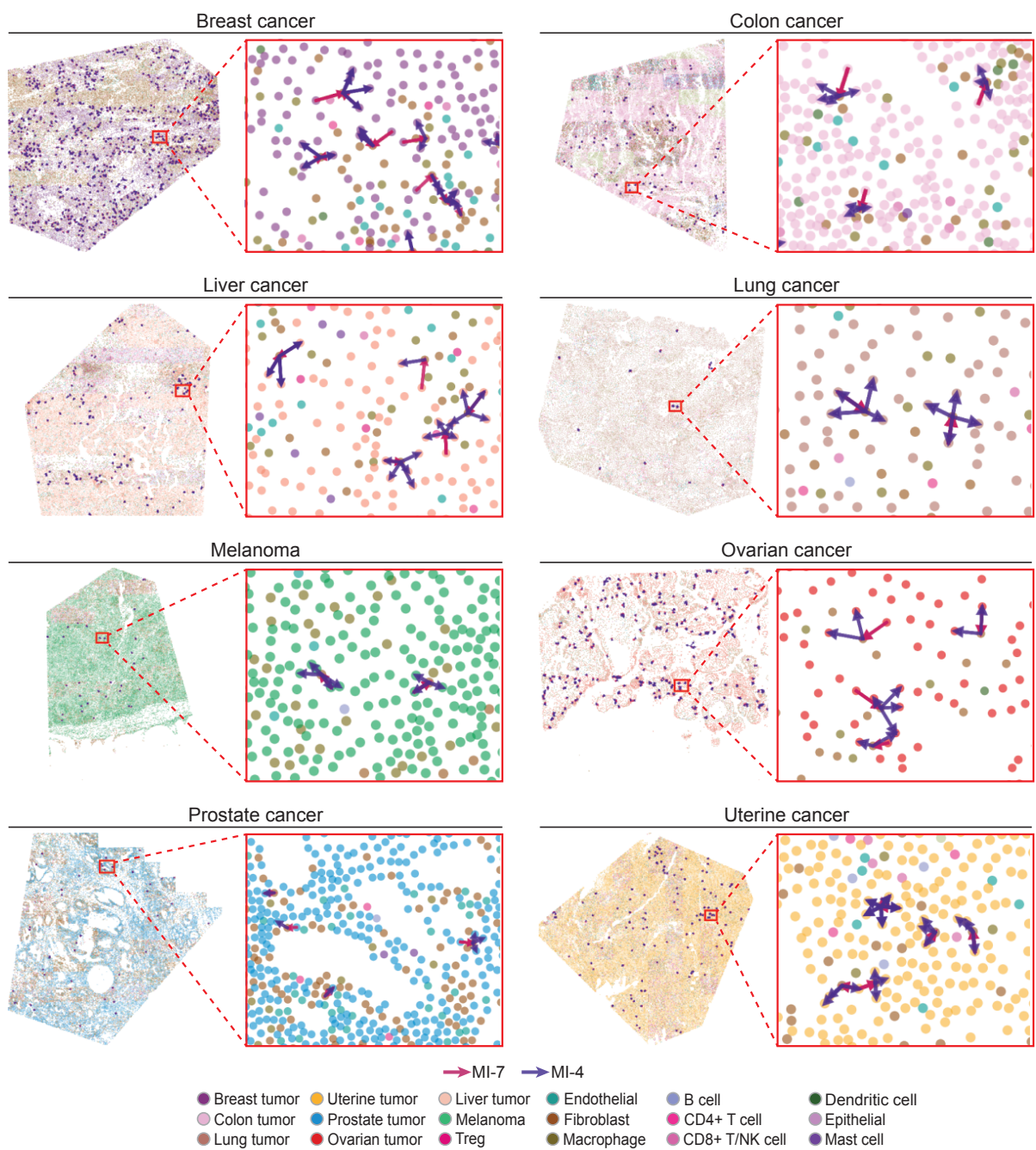

**Figure S28: *In situ* organization of recurrent tumor→fibroblast→tumor MI-7→MI-4 cascades across cancer types.** Representative sub-slices and enlarged regions of interest show directed tumor→fibroblast→tumor triplets comprising upstream MI-7 followed by downstream MI-4 across eight cancer types.

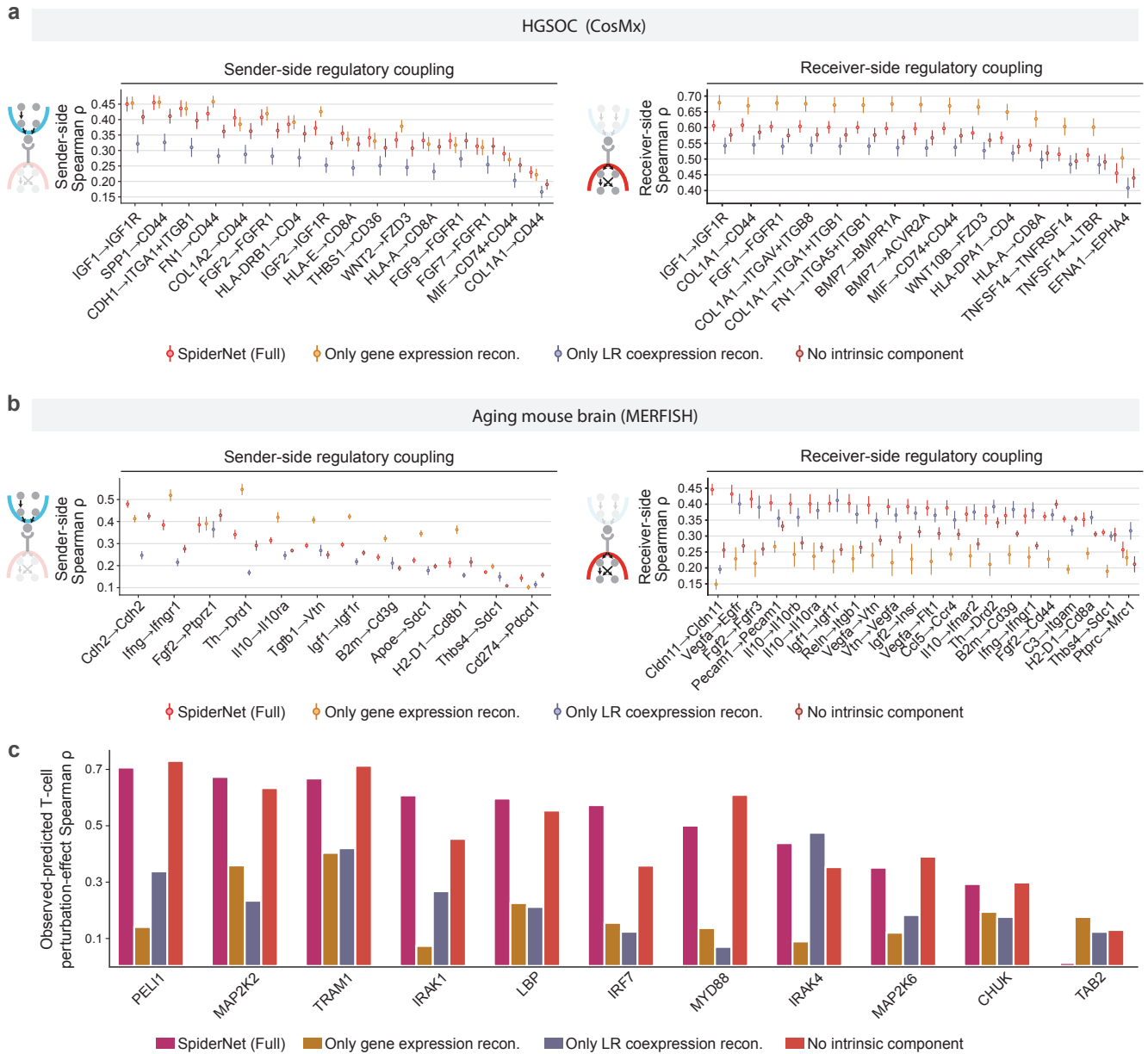

**Figure S29: Component ablation analysis of SpiderNet across regulatory coupling and perturbation prediction.** **a**, Regulatory-coupling benchmark in the HGSOC CosMx dataset. Full SpiderNet was compared with gene-expression-reconstruction-only, LR-co-expression-reconstruction-only, and no-intrinsic variants. Sender- and receiver-side coupling quantify concordance with curated upstream-regulator and downstream-target activities, respectively. **b**, The same regulatory-coupling benchmark in the aging mouse brain MERFISH dataset. For **a** and **b**, points show mean Spearman correlations across tissue slices for each curated LR pair; error bars indicate 95% confidence intervals. **c**, Held-out T-cell perturbation-effect prediction in melanoma Perturb-FISH using the same leave-one-perturbation-out workflow as in Fig. 4b. Bars show Spearman correlations between observed and predicted T-cell perturbation effects across measured genes for each perturbation and model variant.

**Figure S30: MI dimensionality selection and robustness to model hyperparameters.** **a**, Pairwise Spearman correlations between tissue-level LR-pair co-expression profiles in HGSOc CosMx, melanoma Perturb-FISH, aging mouse brain MERFISH, and the Immuno-Oncology MERSCOPE atlas. Yellow boxes mark clique-induced LR-pair components used to determine the MI dimensionality for each dataset. **b**, Stability of cell-cell MI activities across adjacent dimensionality settings in HGSOc ($M=12, 15$, and $18$) and melanoma Perturb-FISH ($M=20, 23$, and $26$). **c**, Stability of HGSOc cell-cell MI activities across spatial neighborhood sizes, comparing $K=5$ with $K=8$ and $K=8$ with $K=10$. In **b** and **c**, heatmaps show Spearman correlations between MI activities inferred under the indicated settings.

**Figure S31: Robustness of SpiderNet outputs to cell-type label misannotation, random seed, and slice subsampling in HGSOc CosMx.** **a**, Pearson correlations of edge-level MI activities with those from the original-label model after randomly misannotating 10%, 20%, or 30% of cell-type labels. Points represent individual MIs. Adjacent misannotation rates were compared using two-sided Wilcoxon rank-sum tests; ns, $P > 0.05$. **b**, Random-seed robustness. SpiderNet was retrained with ten random seeds on all slices and compared with a reference model after one-to-one MI alignment. Points show repeat-level mean Pearson correlations for MI activities and LR-pair, sender-regulator, and receiver-target loadings. **c**, Slice-subsampling robustness. SpiderNet was retrained on ten random 70% subsets of slices and compared with the model trained on all slices. Points show repeat-level mean correlations across aligned MIs. In all panels, center lines indicate medians, boxes indicate the interquartile range, and whiskers extend $1.5 \times \text{IQR}$.

INTERACTIVE ANALYSIS

### Four modules, *one workflow*

MODULE 01

#### Basic analysis

*In situ* MI visualization, top LR / sender / receiver loadings per MI, and pathway and cell-type-pair enrichment.

PICK A DATASET →

MODULE 02

#### Subtype discovery

MI-guided clustering of one cell type with UMAP, DEGs, and GO / KEGG enrichment per subcluster.

PICK A DATASET →

MODULE 03

#### MI cascade

Significant MI cascade pairs, cell-type triplets, in-situ rendering, and DEG / GO at each cascade position.

PICK A DATASET →

MODULE 04

#### Spatial perturbation

*In silico* gene knockdown or cell-type replacement, paired DEG vs. baseline, GO / KEGG enrichment.

PICK A DATASET →

**b**

**c**

**Figure S32: Overview of SpiderNet-Interactive.** **a**, SpiderNet-Interactive provides a web-based workflow for exploring SpiderNet outputs through four linked analysis workflows: basic MI analysis, MI-guided subtype discovery, MI cascade analysis, and *in silico* spatial perturbation. **b**, Example interface of the basic analysis module for visual exploration of MI patterns. **c**, Example interface of the subtype discovery module for visual exploration of MI-guided cell states.
